# Complementary environmental analysis and functional characterization of a plastid diatom lower glycolytic-gluconeogenesis pathway

**DOI:** 10.1101/2022.09.08.507166

**Authors:** Richard G. Dorrell, Youjun Zhang, Yue Liang, Nolwenn Gueguen, Tomomi Nonoyama, Dany Croteau, Mathias Penot, Sandrine Adiba, Benjamin Bailleul, Valérie Gros, Juan José Pierella Karlusich, Nathanaël Zweig, Alisdair R. Fernie, Juliette Jouhet, Eric Maréchal, Chris Bowler

## Abstract

Organic carbon fixed in chloroplasts through the Calvin Cycle can be diverted towards different metabolic fates, including cytoplasmic and mitochondrial respiration; gluconeogenesis; and synthesis of diverse plastid metabolites via the pyruvate hub. In plants, pyruvate is principally produced via cytoplasmic glycolysis, although a plastid-targeted lower glycolytic pathway is known in non-photosynthetic tissue. Here, we characterize a lower plastid glycolytic-gluconeogenesis pathway in diatoms, ecologically important marine algae distantly related to plants. We show that two reversible enzymes required to complete diatom plastid glycolysis-gluconeogenesis, Enolase and PGAM (*bis-* phospho-glycerate mutase), originated through duplications of mitochondria-targeted respiratory isoforms. Through CRISPR-Cas9 mutagenesis, integrative ‘omic analyses, and measured kinetics of expressed enzymes in the diatom *Phaeodactylum tricornutum*, we present evidence that this pathway diverts plastid glyceraldehyde-3-phosphate into the pyruvate hub, and may also function in the gluconeogenic direction. Considering experimental data, we show that this pathway has different roles dependent in particular on day length and environmental temperature, and show that it is expressed at elevated levels in high latitude oceans where diatoms are abundant. Our data provide evolutionary, meta-genomic and functional insights into a poorly understood yet evolutionarily recurrent plastid metabolic pathway.

## Introduction

Each year, over 250 gigatonnes of atmospheric carbon dioxide is assimilated through photosynthesis, with effectively equal contributions from terrestrial plants and aquatic algae (Friedlingstein, Jones et al. 2022). This activity is essential for maintaining planetary climate homeostasis, supporting the entire Earth ecosystem. Carbon assimilated through photosynthesis via the Calvin cycle is diverted into multiple metabolic fates (Raines 2003). In plants, these include gluconeogenesis of glucose-6-phosphate directly in plastids (or chloroplasts), which can then be used in leaf tissue for starch storage (Scialdone, Mugford et al. 2013). Additional metabolites including fatty acids and lipids, amino acids, chlorophyll and carotenoid pigments are synthesised directly in the plastid (Tanaka and Tanaka 2007, Bromke 2013, Maréchal and Lupette 2020, Bai, Cao et al. 2022) (**Fig. 1A**). Many of these plastid metabolic reactions utilize pyruvate, or its adjacent metabolic precursor phospho-*enol*-pyruvate (or PEP), and are referred to collectively as the pyruvate hub (Shtaida, Khozin-Goldberg et al. 2015). In addition, plant photosynthate is exported from the plastids to the cytosol for subsequent glycolysis and respiration in the mitochondria (Moog, Rensing et al. 2015), or for transport to non-photosynthetic tissue (Carrera, George et al. 2021) (**Fig. 1A**).

**Fig. 1.**
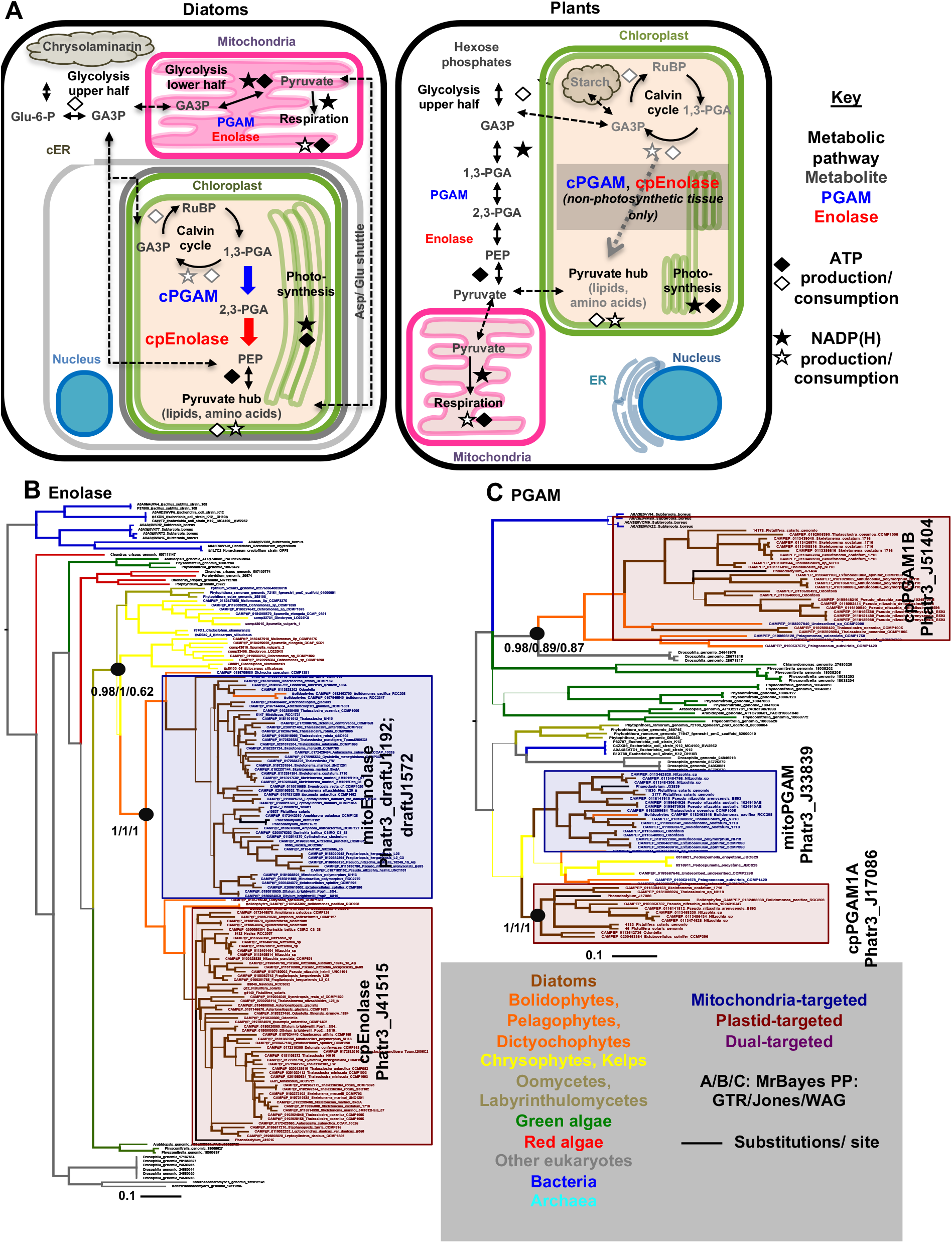
Metabolic context and evolution of the lower half of diatom plastid glycolysis-gluconeogenesis. **A:** schematic comparison of diatom and plant core carbon metabolism, highlighting the localization and functions of two enzymes in the lower half of glycolysis-gluconeogenesis (phospho-glycerate mutase, and enolase) whose localization to the chloroplast can connect endogenous enzymes in the Calvin cycle and pyruvate hub to create a complete glycolytic-gluconeogenic-gluconeogenic pathway. Abbreviations: GA3P-glyceraldehyde-3-phosphate; 1,3-PGA and 2,3-PGA-1,3 and 2,3 bis-phosphoglycerate; Glu-6-P-glucose-6-phosphate; PEP-phospho-enol-pyruvate; RuBP-ribulose *bis*-phosphate; PGAM-phospho-glycerate mutase; cER-chloroplast: ndoplasmic reticulum. **B, C:** consensus MrBayes topologies realised with three substitution matrices (GTR, Jones, WAG) of a 163 taxa x 413 aa alignment of organelle-targeted enolase and 105 taxa x 220 aa alignment of selected organelle-targeted PGAM1 enzymes from diatoms and their closest relatives, identifying recent duplications and recruitments of respiratory glycolytic-gluconeogenic enzymes from the mitochondria to plastid in diatoms and their closest relatives. Branch lines correspond to the frequency with which a given branching relationship was recovered, with thick branches identified by all three substitution matrices.

Plants are classically thought to generate PEP and pyruvate through glycolysis in the cytoplasm, then reimport these metabolites into the plastids (**Fig. 1A**) (Moog, Nozawa et al. 2020). Alongside this, certain plants may synthesize pyruvate hub substrates directly from the Calvin cycle inside the plastid. This conversion is performed by two enzymes, plastid-targeted phospho-glycerate mutase and enolase (henceforth referred to as cpPGAM and epEnolase), which allow the conversion of 1,3-bis-phosphoglycerate from the Calvin cycle to PEP (**Fig. 1A**) (Raines 2003, Andriotis, Kruger et al. 2010). Both Enolase and PGAM have been shown experimentally to be fully reversible enzymes, with bidirectional functions that we henceforth refer to as glycolysis-gluconeogenesis, contrasting with glycolysis and gluconeogenesis to signify enzymatic activities in one direction only (Sutherland, Posternak et al. 1949). Documented plant cpEnolase and cpPGAM enzymes are associated with non-photosynthetic tissues such as seeds and roots (Prabhakar, Löttgert et al. 2009, Fukayama, Masumoto et al. 2015, Troncoso-Ponce, Rivoal et al. 2018). *Arabidopsis thaliana* cpEnolase and cpPGAM knockout lines have limited phenotypes under replete growth conditions (Prabhakar, Löttgert et al. 2009, Andriotis, Kruger et al. 2010, Anoman, Flores-Tornero et al. 2016), raising questions of their overall function.

Diatoms are a eukaryotic algal group that is distantly related to plants, with over one billion years of evolutionary separation between the nuclear and mitochondrial genomes of each species (Nonoyama, Kazamia et al. 2019, Strassert, Irisarri et al. 2021). In contrast to the primary plastids of plants, surrounded by two membranes and of bacterial origin, diatoms possess complex plastids surrounded by four membranes and derived from a eukaryotic red alga, which is likewise ancient (Nonoyama, Kazamia et al. 2019, Liu, Storti et al. 2022). Diatoms are extraordinarily successful in the modern ocean, comprising nearly half of total algal abundance e.g., in environmental sequence data from the *Tara* Oceans expedition (Malviya, Scalco et al. 2016, Behrenfeld, Halsey et al. 2021). Diatoms are particularly abundant in high-latitude and temperate oceans (i.e., the North Atlantic, North Pacific and Southern Oceans) that are characterised by stresses including low temperatures and elongated photoperiods (long days in the summer, and long nights in the winter) (Gilbertson, Langan et al. 2022, Joli, Concia et al. 2024). Previous studies, particularly of the transformable coastal and mesophilic species *Phaeodactylum tricornutum*, have identified multiple strategies that allow diatoms to tolerate photo-stress, including complex inter-organelle metabolite trafficking (Bailleul, Berne et al. 2015, Broddrick, Du et al. 2019, Smith, Dupont et al. 2019) and extensive photoprotective capabilities (reviewed in (Lepetit, Campbell et al. 2022). These data are further supported by extensive environmental (meta-genomic) sequence data such as those of the *Tara* Oceans mission. While the data from these studies relate fundamentally to different species to *Phaeodactylum* (i.e., open-ocean diatoms, including from polar habitats) they may allow us to understand how individual diatom chloroplast proteins function at ecosystem scales, as well as under laboratory conditions (Kazamia, Sutak et al. 2018, Liu, Storti et al. 2022).

Diatom carbon metabolism is highly different to that of plants (Kroth, Chiovitti et al. 2008). Differences include the storage of sugars in cytoplasmic vacuoles (as chrysolaminarin) as opposed to plastidial starch, and the synthesis of most lipid groups (e.g., galactolipids and part of triacylglycerol pathway) directly in the plastid (Zhu, Shi et al. 2016, Huang, Pan et al. 2024). Diatom plastids furthermore possess no known plastid hexose phosphate transporters, which in plants are implicated in plastidial sugar import in storage tissue. Diatoms are instead inferred to exchange sugars with the cytoplasm via triose phosphates only (Moog, Nozawa et al. 2020, Liu, Storti et al. 2022) (**Fig. 1A**). The lower half of respiratory glycolysis-gluconeogenesis in diatoms occurs in the mitochondria, as opposed to the cytoplasm (Kroth, Chiovitti et al. 2008, Río Bártulos, Rogers et al. 2018); and a complete plastid lower half glycolysis-gluconeogenesis, including cpEnolase and cpPGAM proteins, has been inferred from sequenced diatom genomes (Kroth, Chiovitti et al. 2008, Smith, Abbriano et al. 2012, Hippmann, Schuback et al. 2022) (**Fig. 1A**). As diatoms are unicellular and colonial species, plastid glycolysis presumably occurs organelles that perform photosynthesis, contrasting with its predominant association with non-photosynthetic tissues in plants (**Fig. 1A**).

Here, we profile sequence datasets from cultivated and environmental diatoms, perform characterization of *P. tricornutum* CRISPR-CAS9 knockout mutants and measure kinetic activities of expressed enzymes, to infer possible functions of diatom cpEnolase and cpPGAM enzymes. We demonstrate that the genes encoding these enzymes arose from diatom mitochondria-targeted and respiratory isoforms in a common ancestor of all species, contrasting to other algae and plants in which it has a sporadic distribution. We further show that the genes encoding these proteins are most highly expressed at high latitudes in environmental sequence data from *Tara* Oceans, and indeed their expression is induced in *Phaeodactylum* in response to continuous light and low temperature. From *Phaeodactylum* knockout phenotypes, we present evidence that this pathway may have different functions in cells grown under continuous illumination as opposed to light-dark cycling, and at low compared to moderate temperature. We use mutant phenotypes and measured kinetic activities to propose metabolic functions of diatom cpEnolase and cpPGAM under different illumination and temperature regimes. Overall, our data position lower half glycolysis-gluconeogenesis as a modulator of diatom plastid metabolic poise, providing insights into its physiological roles for photosynthetic organisms beyond plants.

## Results

### Distribution and phylogeny of cpEnolase and cpPGAM across photosynthetic eukaryotes

To evaluate the occurrence of plastid-targeted glycolysis across the algal tree of life, we searched for plastid-targeted homologues of *Phaeodactylum tricornutum* and *Arabidopsis thaliana* enolase and PGAM enzymes in 1,673 plant and algal species, considering genomes from JGI PhycoCosm, and transcriptomes from the MMETSP (Marine Microbial Eukaryotic Transcriptome Sequencing Project) and OneKp (One Thousand Plant Transcriptomes) initiatives (Keeling, Burki et al. 2014, Initiative 2019, Grigoriev, Hayes et al. 2021). Plastid-targeting sequences were inferred using both PFAM domain presence and the combined *in silico* predictions of HECTAR, ASAFind, WolfPSort, TargetP and PredAlgo (Emanuelsson, Brunak et al. 2007, Horton, Park et al. 2007, Gschloessl, Guermeur et al. 2008, Tardif, Atteia et al. 2012) (**Table S1,** sheet 1). Plastid lower glycolysis-gluconeogenesis was frequently inferred in diatoms, with 60/101 (59%) libraries with identified enolase and PGAM sequences possessing plastid-targeted versions of each. A lower occurrence (22/69 libraries, 32%) was found amongst close relatives in the stramenopiles (e.g., pelagophytes, dictyochophytes) and other algae with secondary red plastids (cryptomonads, haptophytes; 25/94 libraries, 27%) (**Fig. S1A**). Within primary plastid-harbouring lineages, only angiosperms were inferred to frequently possess plastid-targeted copies of both enzymes (47/537 libraries, 9%). Notably, only 4/127 (3%) occurrences were inferred in primary green algae and none in primary red algae, suggesting that diatom plastid glycolysis does not derive from the secondary red chloroplast ancestor (**Fig. S1A**).

Considering collection sites, diatom species with either plastid glycolysis enzyme typically derive from higher latitudes (mean absolute latitude 45.6°, standard deviation 13.5°, n = 81) than ones that possess neither (mean absolute latitude 38.9°, standard deviation 24.3°, n = 10; one-way ANOVA P = 0.19; **Fig. S1B**). This difference was deemed to be significant for certain diatom groups (e.g., araphid pennate diatoms, **Supplemental Dataset S1**, sheet 1; one-way ANOVA P = 0.012), but was not observed in other algal groups.

Next, we explored the specific origins of *P. tricornutum* plastid Enolase and PGAM sequences from diatoms by building phylogenies of the closest orthologs obtained from other diatoms, the broader taxonomic group to which they belong, the stramenopiles, and two other algal groups, the cryptomonads and haptophytes. These lineages all possess plastids of secondary red endosymbiotic origin, surrounded by four membranes, which are likely to be closely related to one another (Strassert, Irisarri et al. 2021), but also contain non-photosynthetic members (e.g., oomycetes in stramenopiles) which only possess respiratory (i.e., mitochondria-targeted) lower half glycolytic enzymes (Río Bártulos, Rogers et al. 2018). Single-gene trees were made for the conserved domains of all organelle-targeted Enolase and PGAM sequences from 289 cryptomonad, haptophyte and stramenopile genomes and transcriptomes, plus all orthologs from 85 further genomes selected from across the tree of life, based on a previously defined pipeline (**Supplemental Dataset S1**, sheet 2-9). **Figs. 1B** and **1C** show consensus MrBayes trees realised with GTR, Jones and WAG substitution matrices for species with both identifiable plastid- and mitochondria-targeted orthologs of each protein.

The obtained topologies revealed multiple evolutionary origins for plastid Enolase and PGAM sequences from mitochondria-targeted (respiratory) enzymes, with diatom plastid isoforms typically having recent and/or diatom-specific evolutionary origins. Diatom cpEnolase sequences resolve in a well-supported clade with plastid-targeted enzymes from bolidophytes, dictyochophytes and pelagophytes, which are sisters to diatoms in the stramenopiles (Río Bártulos, Rogers et al. 2018, Nonoyama, Kazamia et al. 2019), followed by mitochondria-targeted proteins from these groups (MrBayes PP = 1.0 under all studied matrices, **Fig. 1B**), other photosynthetic (chrysophytes) and non-photosynthetic stramenopiles (oomycetes; MrBayes PP = > 0.95 under GTR and Jones matrices, **Fig. 1B**). This indicates a duplication and recruitment of the host-derived mitochondria-targeted protein to the plastid within a common ancestor of the diatoms, pelagophytes and dictyochophytes. A broader evaluation of cpEnolase distribution suggests further duplications and plastid retargeting of mitochondria-targeted enolase proteins in both the chrysophytes and cryptomonads (**Fig. S2**).

The PGAM phylogeny revealed at least two closely-related families of plastid-targeted diatom enzymes, both likely derived from host mitochondrial isoforms. The cpPGAM1A clade (typified by the *P. tricornutum* protein Phatr3_J17086) was closely related to mitochondrial-targeted proteins found across the stramenopiles (MrBayes PP = 1.0 under all studied matrices, **Fig. 1C**), followed by plastid-targeted proteins from chrysophytes and mitochondria-targeted oomycete proteins. Similarly, the cpPGAM1B (Phatr3_J51404) clade included mitochondrial-targeted proteins from pelagophytes and dictyochophytes (MrBayes > = 0.85 under all studied matrices, **Fig. 1C**), and plastid-and mitochondria-targeted enzymes from the chrysophytes (**Fig. S3**). Further duplications and plastid recruitments of mitochondria-targeted PGAM proteins were again visible in the haptophytes and cryptomonads (**Fig. S3**).

A final plastid-targeted protein annotated as PGAM in the version 3 *P. tricornutum* genome (Rastogi, Maheswari et al. 2018), hereafter termed PGAM2, was identified exclusively in diatoms, pelagophytes, and haptophytes (**Fig. S4**), with limited homology to PGAM1 (BLASTp e-value > 1.0 in pairwise protein-protein searches). Only PGAM1 contain an annotated phospho-glyceromutase active site (IPR005952) per InterProScan, while both PGAM1 and PGAM2 contain the same PFAM (histidine phosphatase, PF03000) per PFAMscan (Jones, Binns et al. 2014, Mistry, Chuguransky et al. 2020). PGAM2 enzymes were predominantly mitochondria-targeted, with plastid- or dual-targeted isoforms amongst diatoms only identified in *P. tricornutum* (Phatr3_J37201, and a more divergent copy Phatr3_J47096) and three species in which it is inferred to have evolved independently (**Fig. S4**).

### Plastidial localisation and expression dynamics of Phaeodactylum lower glycolysis enzymes

To confirm plastid localization of *P. tricornutum* cpEnolase and cpPGAM, eGFP-tagged copies of three proteins (Phatr3_J41515, cpEnolase; Phatr3_J17086, cpPGAM1A; Phatr3_J37201, cpPGAM2) were expressed in *P. tricornutum* Pt1.86 cells via biolistic transformation. The observed GFP fluorescence patterns were coincident with chlorophyll autofluorescence, consistent with *in silico* targeting predictions in each case and confirming plastid localization (**Figs. 1D**, **S5**). We note that both cpEnolase (named per its previous annotation Phatr2_J56418) and cpPGAM1B (named Phatr2_J42857) have been independently localised with GFP to the *P. tricornutum* plastid in a separate study (Río Bártulos, Rogers et al. 2018).

Next, we considered previously published experimental proteomic data of plastid-enriched and *Phaeodactylum* total cellular fractions, following (Huang, Pan et al. 2024). Both cpEnolase and cpPGAM1A were detected in multiple plastid-enriched and total cell proteome samples, respectively forming 0.040% and 0.0033% of the mean plastid-enriched total proteome (**Fig. S6A**). These abundances were analogous to the total abundances and plastid enrichment ratios found for other Calvin cycle and plastidial carbon metabolism enzymes, e.g. pyruvate kinase (Phatr3_J22404), ribose-5-phosphate isomerase (Phatr3_J13382), and ribulose-5-phosphate epimerase (Phatr3_J53395). Both mtEnolase (Phatr3_draftJ1572) and mtPGAM (Phatr3_J33839) were also detected in plastid-enriched fractions, which may relate to close associations observed between the *Phaeodactylum* plastid and mitochondria, (Bailleul, Berne et al. 2015, Uwizeye, Decelle et al. 2020). No other organelle-associated enolase or PGAM enzymes were detected, including cpPGAM1B and cpPGAM2, suggesting that they are present at low abundances in the *Phaeodactylum* cell (**Fig. S6A; Supplemental Dataset S2,** sheet 1).

We further considered the transcriptional dynamics of *Phaeodactylum* lower plastid glycolysis proteins, using a pooled and ranked dataset of normalised microarray and RNAseq data to identify genes that are co-expressed with one another and which may perform linked cellular functions (Ashworth, Turkarslan et al. 2016, Ait-Mohamed, Novák Vanclová et al. 2020, Liu, Storti et al. 2022) (**Fig. S6B**; **Supplemental Dataset S2**, sheet 2-3). From these data, cpEnolase and cpPGAM1A showed strong, positive coregulation to one another (r = 0.868, P < 10^-05^), with cpPGAM1A the second most strongly coregulated gene to cpEnolase across the entire *Phaeodactylum* genome. Other cpPGAMs showed much weaker coregulation to both cpEnolase and cpPGAM1A, including cpPGAM1B (cpEnolase *r*= 0.432; cPGAM1A *r* = 0.473) and cpPGAM2 (cpEnolase *r*= 0.490; cPGAM1A *r* = 0.478; **Fig. S6B**). The coexpression of cpEnolase and cpPGAM1A and accumulation of both encoded proteins in the plastid suggest that they possess linked metabolic functions.

We then explored under what conditions cpEnolase and cpPGAM genes are likely to be highly expressed, considering RNAseq (**Fig. S7A-C**) and microarray (**Fig. S7D**) data (**Supplemental Dataset S2**, sheet 4). These data suggested that nutrient limitation does not directly induce the expression of the chloroplast-targeted glycolysis proteins, with the ratio of expression of genes encoding chloroplast-versus mitochondria-targeted copies of each enzyme either remaining unchanged in published nitrate limitation (nitrate reductase knockout) and iron limitation data (**Fig. S7A, B;** one-way ANOVA P < 0.05 (Smith, Gillard et al. 2016, McCarthy, Smith et al. 2017)). In an analogous study of phosphate limitation, we even observed a lower ratio of plastid to mitochondrial glycolysis gene expression in phosphate-starved versus -replete and -replenished cell lines (ptEnolase/ mtEnolase ratio one way ANOVA P = 0.009, ptPGAM/ mtPGAM ratio one-way ANOVA P = 0.049 (**Fig. S7C**; (Cruz de Carvalho, Sun et al. 2016)).

In contrast, we observed clear impacts of light quality and daylength on plastidial glycolysis gene expression. In an RNAseq study of the effects of the Circadian cycle on Fe-limited and -replete *Phaeodactylum* cells (Smith, Gillard et al. 2016), a much higher ratio of plastid to mitochondrial Enolase gene expression was identified in samples harvested twelve hours post-illumination than other time points (**Fig. S7B**; one-way ANOVA P = 4 x 10^-05^). From a similar meta-analysis of microarray data (Ashworth, Turkarslan et al. 2016), cpEnolase showed greatest relative fold-change in RNA samples (one-way ANOVA P < 10^-05^) collected between 8 and 16h after the light onset, and were strongly suppressed following 30 minutes of white, red, green and blue light treatment (**Fig. S7D**). cpPGAM showed the same trends albeit with lower expression in normal light 10.5 h after the light induction period than 16h (**Fig. S7D; Supplemental Dataset S2,** sheet 4). Both cpEnolase and cPGAM1A showed strong suppression in microarray data obtained following four hours compared to thirty minutes dark incubation (one-way ANOVA, P 0.05); and two days dark incubation compared to two-days high light treatment (one-way ANOVA P < 10^-05^; **Fig. S7D**) suggesting that these effects relate to light perception.

Finally, we performed quantitative RT-PCR of cpEnolase and cpPGAM1A genes from wild-type *Phaeodactylum* cells under different conditions (**Fig. 2B**). qRT-PCRs were performed on RNA collected from late exponential phase cells at the subjective day mid-points at 19°C and 12 h: 12 h light: dark cycling (19C LD); the same time but for cells grown under 19°C and 24 h continuous light (19C CL); and the same time but for cells grown under 8°C and 24 h continuous light (8C CL), considering *Tara* Oceans sampling data (see below; **Supplemental Dataset S2**, sheet 4). qRT-PCRs were performed using two RT-PCR amplicons for each gene and two normalisation references (RPS, and TBP) previously shown to have invariant expression under Circadian cycles in *Phaeodactylum* (Sachse, Sturm et al. 2013).

**Fig. 2.**
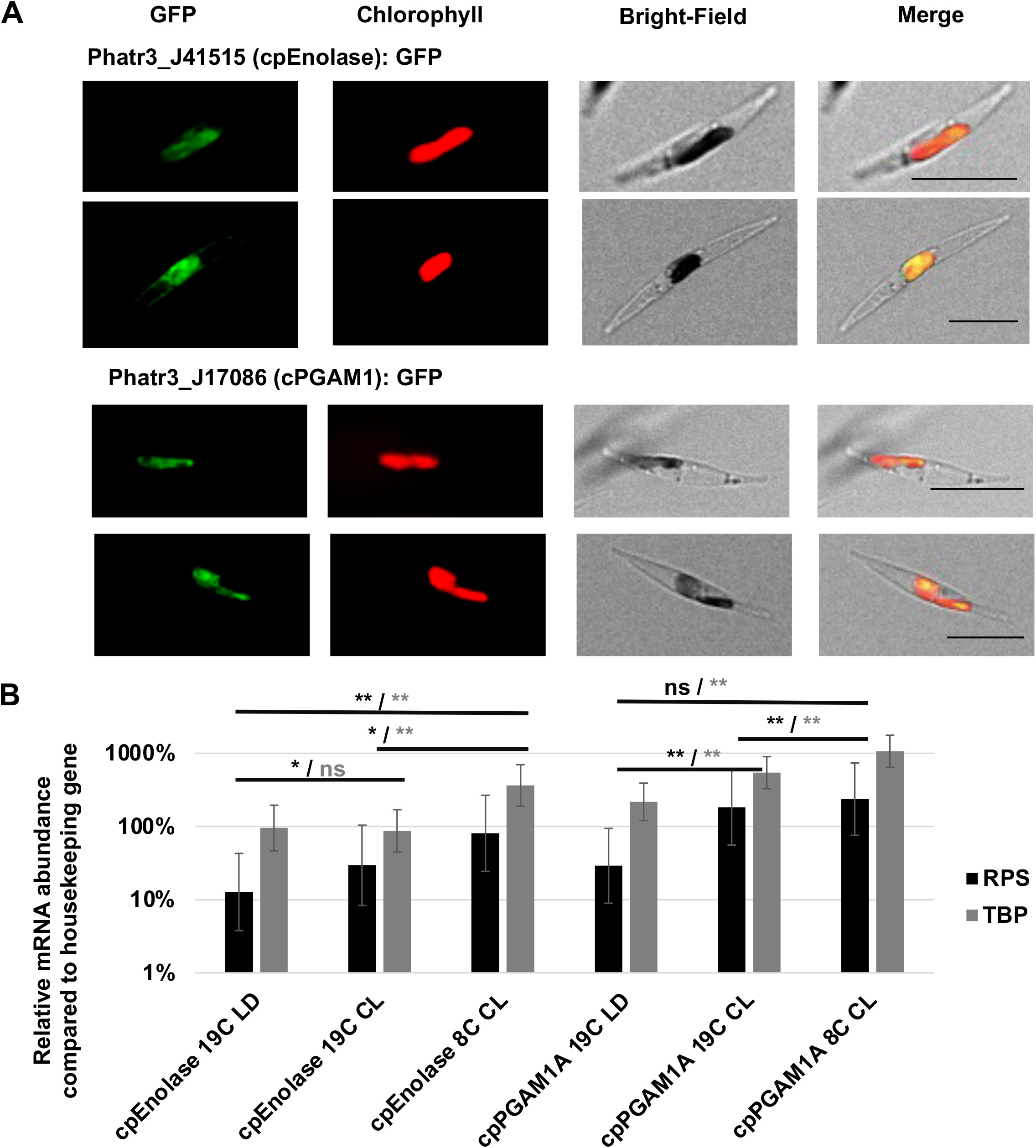
Localisation and expression of *Phaeodactylum tricornutum* cpEnolase and cpPGAM1A. **A:** individual channel and overlay images of GFP-tagged full-length cpEnolase (Phatr3_J41515) and cpPGAM1A (Phatr3_J17086) constructs (green), chlorophyll (red) and bright-field images of transformant *P. tricornutum* lines. Scale bar: 10 μm. **B:** quantitative RT-PCR (qRT-PCR) of cpEnolase and cpPGAM1A expression in mid-exponential phase cells harvested at the mid-point of a 19C 12h light: 12h dark cycle (19LD); at the same timepoint but under 24h continuous light (19CL); and under 8C and continuous light (8CL). Relative expression levels were normalised against two housekeeping genes (RPS, RNA polymerase subunit 1; TBP, Tata binding protein) that show invariant expression in response to light cycles in *Phaeodactylum* per (Sachse, Sturm et al. 2013). *: significantly different expression levels, one-way ANOVA, P < 0.05; **, the same, P < 0.001.

Both cpEnolase and cpPGAM1A showed transcriptional responses to light and temperature, with different responses dependent on normalisation reference. The expression of cpEnolase was inferred to be increased in 19C CL relative to 19C LD when normalised to RPS (fold-change: 2.31, one-way ANOVA P = 0.028) although no difference was measured by normalisation to TBP. In contrast, the expression of cpEnolase was found to be significantly higher under 8C CL than 19C CL conditions considering both RPS (fold-change: 2.74, P = 0.015) and TBP (fold-change: 4.17, P = 0.001; **Fig. 2B**). cPGAM1A expression was inferred to be increased in 19C CL relative to 19C LD conditions normalised to both RPS (fold-change: 6.33, P = 0.003) and TBP (fold-change: 2.50, P = 0.002); but was only inferred to increase in 8C CL relative to 19 CL conditions normalised to TBP (fold-change: 1.96, P = 0.005; **Fig. 2B**). In total these data suggest additive effects of both continuous light and low temperature on *Phaeodactylum* cpPGAM1A and cpEnolase expression.

### Environmental roles of diatom cpEnolase and cpPGAM inferred from meta-genomics

Next, we considered general patterns of transcriptional co-regulation of diatom cpEnolase and cpPGAM sequences in environmental sequence data from *Tara* Oceans. First, we used a previously benchmarked pipeline, based on combined hmmer, reciprocal BLAST and phylogenetic filtration (Liu, Storti et al. 2022) to identify *Tara* Oceans meta-genes that reconcile exclusively with plastid-targeted proteins from cultured diatom species, to the exclusion of non-diatom and non-plastid homologs (**Fig. S8A**). Amongst the retained meta-genes likely to be N-terminally complete (BLAST homology within the first 40 residues of a *P. tricornutum* sequence), a majority have consensus plastid-targeting sequences (enolase: 38/ 78-49%, PGAM: 58/ 97-60%). Only a very small number (one enolase, 10 PGAM) possess mitochondrial or endomembrane localizations, suggesting that they principally correspond to plastid-targeted environmental homologs of each protein (**Fig. S8B, Supplemental Dataset S3**, sheet 11).

Within *Tara* Oceans data, the greatest relative abundances of diatom cpEnolase and cpPGAM1 were observed in meta-transcriptome (metaT) data in stations from both high northern and southern latitudes (**Fig. 3**). We observed these trends concordantly in both surface and deep chlorophyll maximum (DCM) samples from 0.8 - 2000 µm size filtered (**Fig. 3A**); and in individual size fractions (0.8-3/5 µm, 5-20 µm, 20-180 µm, 180-2000 µm (**Fig. S9**), suggesting broad reproductibiilty across diatoms independent of cell size and depth. These levels were notably greater than equivalent levels in meta-genome (metaG) data (**Figs. 3B**, **S9**).

**Fig. 3.**
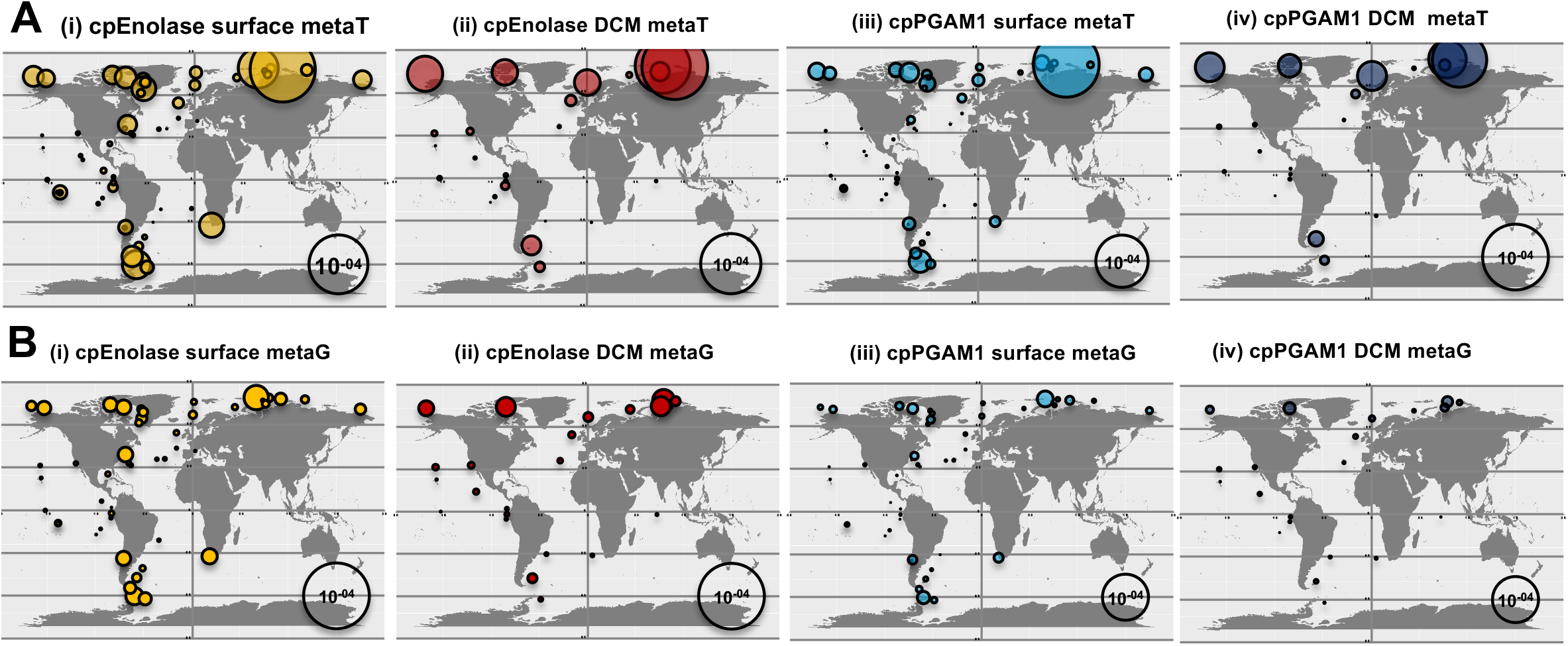
Environmental distributions of diatom plastidial lower half glycolysis-gluconeogenesis meta-genes. Total transcriptome **(A)** and genome **(B)** relative abundances, sampled from all (0.8-2000µm) size fractions and surface layer **(i,iii)** or DCM **(ii,iv)** stations for *Tara* Oceans meta-genes phylogenetically resolved to diatom cpEnolase **(i,ii)** and cpPGAM1 **(iii,iv).** These data provide a global overview of *Tara* Oceans meta-gene abundances, in complement to data from individual size fractions shown in **Fig. S9.** These data demonstrate higher meta-transcript abundance without commensurate increases in meta-gene abundance at high northern and southern latitudes.

To confirm that this was due to a greater expression of cpPGAM and cpEnolase genes, as opposed to being purely driven by the greater relative abundance of diatoms in high latitude *Tara* Oceans stations, we performed multiple normalization tests (**Fig. S10**; **Supplemental Dataset S3**, sheet 10). First, metaT abundances calculated for each gene in the 0.8-2000 μm size fraction were divided by the total relative abundance of all diatom metaT sequences, providing the total proportion of each diatom meta-transcriptome occupied by cpEnolase and cpPGAM. These normalisations showed positive correlations to latitude in both surface and DCM depth fractions, with the greatest relative abundances (> 0.1% total diatom mapped transcripts) typically occurring in stations > 60° (**Fig. S10A**). The observed Pearson correlations to latitude were significantly positive ( surface cpEnolase R^2^ = 0.18, P < 10^-05^, cpPGAM1A R^2^ = 0.23, P < 10^-05^; DCM cpEnolase R^2^ = 0.53, P < 10^-05^, cpPGAM1A R^2^ = 0.59, P < 10^-05^) (**Supplemental Dataset S3**, sheet 10). More broadly across, the metaT-normalised relative abundance levels showed clearest positive correlations to daylength and negative correlations to temperature. No other parameters (e.g., nutrient concentrations) showed as clear correlations to chloroplast glycolysis metaT relative abundances (**Supplemental Dataset S3**, sheet 10).

Alongside this, the metaT abundances obtained for diatom cpEnolase and cpPGAM genes were compared (via log normalisation, to allow the inclusion of zero values) to the relative abundances calculated for the meta-genomic (metaG) sequences of the same genes (**Fig. S10B**). This can be taken as an indicative measurement of the relative ratio of transcript versus gene abundances for each meta-gene, i.e. in effect its expression level. These measurements showed a weaker but still significant positive correlation to latitude for cpEnolase surface fractions (R^2^ = 0.10, one-tailed *F*-test, P < 0.05) and for both genes in DCM fractions (cpEnolase R^2^ = 0.28, one-tailed *F*-test P < 0.05, cpPGAM1 R^2^ = 0.29, one-tailed *F*-test P< 0.05 (**Supplemental Dataset S3**, sheet 10). For both genes and in both depth fractions, two individual stations within the Arctic (Station 173, 78.93-78.96°N; Station 188, 78.25°-78.36°N) were observed to have extremely high metaT to metaG ratios ((log_10_(1+metaT)-log_10_(1+metaG)) > 3-5) that disrupted the linear relationship between normalised metaT and latitude and point to specifically high expression of chloroplast glycolysis genes in polar waters. To correct for the impacts of these stations, ranked (Spearman) correlation values were also calculated for normalised chloroplast glycolysis metaT expression levels. Significant positive correlations with latitude were detected in multiple individual size fractions and depths (0.8-5, 3/5-20, 20-180, 180-2000 μm), including for cpPGAM1 metaT normalised against metaG in surface 3/5-20 (one-tailed *F*-test, P < 10^-05^), 20-180 (one-tailed *F*-test; P < 10^-05^) and 180-2000 (one-tailed *F*-test, P < 0.05) μm fractions (**Supplemental Dataset S3**, sheet 10).

The transcriptional preference of diatom cpEnolase and cpPGAM1 for high latitudes contrasted strongly with PGAM2, which showed equivalent relative abundance in stations from the temperate South Pacific and Atlantic as stations from the Arctic and Southern Oceans (**Fig. S11**; **Supplemental Dataset S3**, sheet 10). In certain size fraction and depth combinations (e.g., DCM 0.8-3, and 3/5-20 μm fractions, normalised against metaG abundances; and surface and DCM 180-2000 μm fractions normalised against all diatom metaT abundances) PGAM2 metaT abundances even demonstrated significant negative correlations to latitude (**Supplemental Dataset S3**, sheet 10).

Finally, we tested whether the occurrence of plastidial lower glycolysis may correlate to algal abundance at high latitudes. For this, we screened single-cell and meta-genome assembled genomes (sMAGs) from *Tara* Oceans for potential plastid-targeted Enolase and PGAM enzymes, using similar reciprocal BLAST best hit, PFAM annotation and *in silico* targeting prediction techniques as previously used for cultured algae (**Fig. S12**; **Supplemental Dataset S**3, sheet 11) (Delmont, Gaia et al. 2022). We emphasise these results are preliminary, as many of these genomes are incomplete, and gene non-detection does not formally confirm absence (Delmont, Gaia et al. 2022, Pierella Karlusich, Nef et al. 2023). For each sMAG, we considered presence or absence of possible plastid-targeted Enolase and PGAM sequences; taxonomic assignation of the MAG; and mean mapped vertical coverage of each MAG in each station (i.e., depth and breadth of the coverage of sequences recruited to each genome), as a proxy for abundance, regardless of whether the plastid-targeted glycolysis genes associated with each sMAG were detected (**Fig. S12**).

Across 291 eukaryotic algal sMAGs, 32 were found to possess both plausible cpEnolase and cpPGAM proteins, and a further 84 could be assigned either one or the other (**Fig. S12**). As expected, diatoms were found to possess plastid-targeted glycolysis much more frequently than other groups, with 17/ 49 of the diatom sMAGs found to possess both chloroplast-targeted Enolase and PGAM enzymes, and a further 20 one of the two only (**Fig. S12**). We also detected probable complete plastid glycolysis pathways in 10 further sMAGs belonging to lineages (pelagophytes, dictyochophytes, haptophytes, chrysophytes, and bolidophytes) previously inferred to possess complete plastid-targeted glycolysis pathways amongst cultured species (**Fig. S2, S3**). Surprisingly, given the relative paucity of this pathway in cultured primary green algae, we finally identified fiuve putative chlorophyte sMAGs with both plausible cpEnolase and cpPGAM1A proteins (**Fig. S12**). All five of these sMAGs (TARA_AON_82_MAG_00297, AOS_82_MAG_00181, ARC_108_MAG_00063, ARC_108_MAG_00100, and PSW_86_MAG_00289) are assigned as novel members of the genus *Micromonas* which is abundant at high latitudes (Lovejoy, Vincent et al. 2007, Worden, Lee et al. 2009, Delmont, Gaia et al. 2022). Of note, no cultured *Micromonas* are inferred to possess this pathway **(Supplemental Dataset S1,** sheet 1). We therefore infer, in particular, that the recurrent *Micromonas* sMAG isoform may be a novel plastid glycolysis pathway specific to uncultivated taxa.

Considering the biogeography of each sMAG, we note that diatoms that possess complete lower half plastidial glycolysis pathways shows positive correlations between mean mapped vertical coverage and absolute station latitude, albeit only in DCM fractions (**Fig. S12A**; *r*= 0.517, P< 0.001, while positive correlations to lartitude were observed for diatom sMAGs possessing one of cpEnolase or cpPGAM only atboth depths (**Fig. S12A**; surface *r*= 0.313, two-tailed *t*-test P= 0.003, DCM *r*= 0.614, P< 0.001). This trend was not however observed for diatom sMAGs lacking plastid-targeted copies of both proteins, which showed non-significant and even weakly negative correlations to absolute latitude (**Fig. S12A**; surface *r*= -0.069, DCM *r*= 0.192, P> 0.1). No clear association between plastidial lower half glycolysis and occupancy at high latitudes was observed for other algal groups, with the exception of chlorophytes, in which the presence of both cpEnolase and cpPGAM1A showed a strong association with abundance in high (and particularly) Arctic latitude stations (surface *r* = 0.508, P< 0.001; DCM *r* = 0.386, P = 0.017; **Fig. S12B, S12C**).

### Growth and photo-physiology of Phaeodactylum cpEnolase and cpPGAM1A knockouts across light and temperature conditions

We generated homozygous CRISPR knockout lines for both cpEnolase and cpPGAM1A in the model diatom *P. tricornutum*. cpPGAM1A was selected over other PGAM (cpPGAM1B, cpPGAM2) isoforms because of its transcriptional co-regulation to cpEnolase and occurrence in measurable quantities in plastid proteome data (**Fig. S6**; **Supplemental Dataset S2**) and latitudinal expression correlation in *Tara* Oceans (**Figs. 3**, **S11**).

Multiple CRISPR knockout lines were generated from two regions with unique sequences in the *P. tricornutum* genome for each gene (cpEnolase CRISPR region 1 *n*= 4, CRISPR region 2 *n*= 3; cpPGAM1A CRISPR region 1 *n*= 2, CRISPR region 2 *n*= 3) (**Fig. S13A**). Each CRISPR line was verified by sequencing to be homozygous and to contain a frame-shift mutation sufficient to impede translation of the encoded protein (**Fig. S13A**). Commercial antibodies against enolase and PGAM were found not to specifically label cpEnolase and cpPGAM1A in Western Blots, and so we inferred protein relative expression level by qRT-PCR using recognised *P. tricornutum* housekeeping genes as above (Sachse, Sturm et al. 2013, Zhang, Sampathkumar et al. 2020). The measured knockout mRNA abundance in each line was significantly lower (1.8-39 %) than that identified in empty vector control mRNA (*n* = 4, one-way ANOVA, P < 0.05) 19C LD conditions, **Fig. S13B**). This is consistent with effective knockdown of mutated genes e.g., via non-sense mediated decay (Chang, Imam et al. 2007).

Next, we performed growth curves of cpEnolase and cpPGAM1A knockout lines compared to empty vector controls (**Fig. 4**; **Supplemental Dataset S4**, sheets 3-6). We chose to target changes in light and temperature, given that both show clear associations observed with cpPGAM1A and cpEnolase in *Phaeodactylum* gene expression and *Tara* Oceans data (**Fig. 2**, **3**), using the three conditions (19C CL, 19C LD, and 8C CL) previously tested for qRT-PCR. We note that these conditions are relevant to the environmental conditions in which the type culture of Phaeodactylum (strain CCAP 1055/1) was collected (Irish Sea, 53.5°N) with measured sea temperatures (1960-1999) between 3°C and 17°C ; and day lengths between 7 and 17 hours (Young and Holt 2007, Gachon, Heesch et al. 2013).

**Fig. 4.**
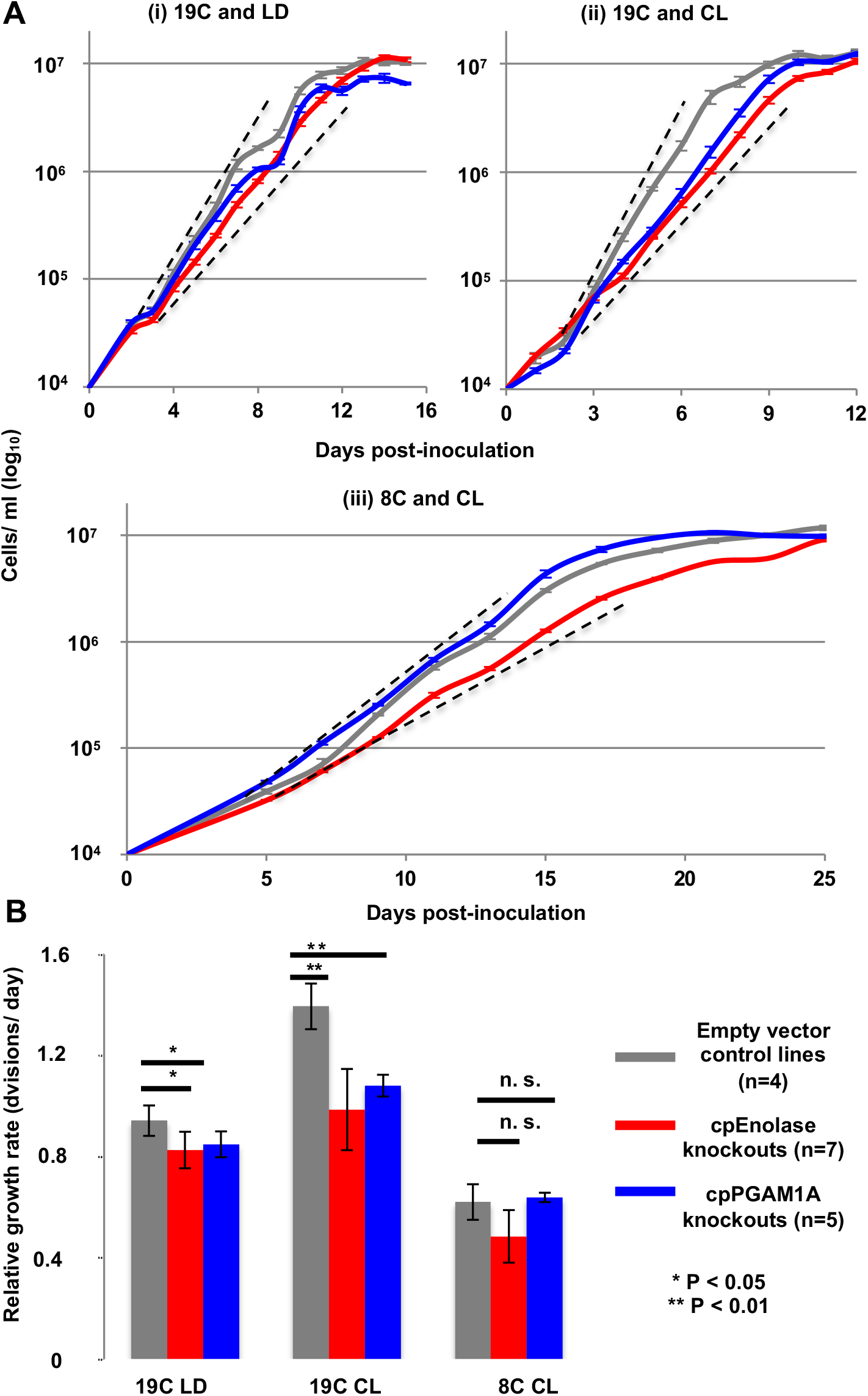
Growth phenotypes of cpEnolase and cpPGAM1A CRISPR-Cas9 knockout mutant and zeocin-resistant empty vector control *P. tricornutum* lines. **A:** exemplar growth curves from single experiments realised for *P. tricornutum* lines in 50 μE m^-2^ s^-1^ illumination, non-shaken cultures and replete ESAW media, under three conditions-(i) 19°C and 12h light: 12h dark Circadian cycles (« 19C LD »); (ii) 19°C and 24h continuous light (« 19C CL »); and (iii) 8°C and 24h continuous light (« 8C CL »). Hashed black lines show the approximative concentrations (between 5 x10^4^ and 4 x 10^6^ cells ml^-1^) over which growth rates were calculated). **B:** mean relative log phase growth rates of each genotype under each condition, measured through a minimum of three biological replicates and two technical repetitions (six measurements per line, minimum 24 measurements per genotype), over five time-points with linear (*r*^2^ > 0.95 relationship between log cell density and time). Asterisks indicate significant differences as inferred by one-way ANOVA. An alternative version of this figure showing absolute growth rates of individual cell lines is provided in **Fig. S14.**

Under 19C LD conditions, plastid glycolysis-gluconeogenesis knockout lines showed an approximately 10-15% reduction in relative growth rate compared to empty vector controls (cpEnolase growth rate 0.83± 0.06 cells day^-1;^ cpPGAM1A growth rate 0.85± 0.07 cells day^-1;^ empty vector growth rate 0.94 ± 0.05 cells day^-1^; **Fig. 4**, **S14; Supplemental Dataset S4**, sheet 3; cpEnolase growth rate 87.7% control and cpPGAM1A growth rate 90.1% control, one-way ANOVA, two-tailed P < 0.05). Under 19C CL, knockout lines showed a 25-30% reduction in relative growth rate compared to controls (cpEnolase growth rate 0.99± 0.16 cells day^-1^; cpPGAM1A growth rate 1.08± 0.04 cells day^-1^; empty vector growth rate 1.39 ± 0.09 cells day^-1^; **Fig. 4**, **S14**; **Supplemental Dataset S4**, sheet 4; cpEnolase growth rate 70.7% control and cpPGAM1A growth rate 77.5% control, one-way ANOVA, two-tailed P < 0.01). Under 8C C, overlapping growth rates were observed for knockout and control lines, albeit with a possible reduction in cpEnolase knockout growth rate (cpEnolase relative growth rate 0.49± 0.10 cells day^-1^, cpPGAM1A growth rate 0.64± 0.02 cells day^-1^, empty vector growth rate 0.62± 0.07 cells day^-1^; **Fig. 4**, **S14; Supplemental Dataset S4**, sheet 5; cpEnolase growth rate 78.1% control and cpPGAM1A growth rate 102.9% control; one-way ANOVA, two-tailed P non-significant).

To test the possibility of off-target effects of the CRISPR constructs, we complemented mutant lines with blasticidin resistance genes linked to either cpEnolase-GFP or cpPGAM1A-GFP modified to remove all CRISPR target sequences (**Supplemental Dataset S4**, sheet 2) (McCarthy, Smith et al. 2017, Buck, Río Bártulos et al. 2018). Despite an overall lower growth rate in all blasticidin-resistant lines compared to primary transformants, and within-line variation, comparative growth curves of 47 complemented versus placebo transformed mutant lines revealed increased growth rates in complemented cpEnolase and cpPGAM1A versus blank transformed knockout lines under 19C CL and 19C LD (**Supplemental Dataset S4**, sheet 7; one-way one-way ANOVA, two-tailed P, < 0.05). By contrast, complemented knockout line growth rates overlapped with empty vector controls either transformed with cpEnolase or blank complementing vectors, indicating effective recovery of mutant phenotypes (**Supplemental Dataset S4**, sheet 7).

Finally, we performed comparative photophysiological measurements of knockout lines in the two conditions (19C LD and 19C CL) where they presented a growth phenotype (see Methods). Our data indicate that the presence/ absence of these enzymes does not significantly impact photosynthetic performance. The light dependencies of either electron transfer rate through photosystem II (PSII) (rETR(II)) or photoprotection (non-photochemical quenching, NPQ) were very similar between control and knock-out lines (**Fig. S15A**; **Supplemental Dataset S4**, sheets 8-11). A slight but significant increase in the functional absorption cross-section of photosystem II (σPSII) was found under 19C CL in both cpEnolase (319.3± 22.5) and cpPGAM1A knockouts (306.6± 11.6) compared to controls (292.3± 8.2; one-way ANOVA, P < 0.05) (Gorbunov, Shirsin et al. 2020). This elevation was suppressed in both complemented lines (**Fig. S15B**; **Supplemental Dataset S4**, sheet 11).

### Gene expression profiling of Phaeodactylum cpEnolase and cpPGAM1A knockouts

Next, we investigated the impacts of disruption of plastid glycolysis on diatom metabolism beyond photosynthesis. First, we performed quantitative RNA-seq analysis using 63 RNA samples drawn from multiple knockout and empty vector lines under all three physiological conditions (19C LD, 19C CL, and 8C CL; **Supplemental Dataset S5**, sheet 1; Materials and Methods). 8C CL was targeted despite the absence of a growth phenotype associated with this line due to the high levels of cpEnolase and cpPGAM1A gene expression inferred from qRT-PCR data (**Fig. 2B**; **Fig. 4**) Complete results are provided in **Supplemental Dataset S5**, sheets 5-11. Both cpEnolase and cpPGAM1A mRNA were found to significantly under-accumulate in the corresponding knockout lines, consistent with qRT-PCR analysis (**Fig. S13B**) and suggesting maintenance of the mutant genotypes throughout RNA sequencing; while cpPGAM1B (Phatr3_J51404) but not cpPGAM2 (Phatr3_J37201) was upregulated in cpPGAM1A knockouts but not cpEnolase knockouts under 19C CL conditions, which may suggest compensatory functions between cpPGAM1A and cpPGAM1B (**Supplemental Dataset S5**, sheet 12).

Genome-scale enrichment analyses of the *in silico* localizations of proteins encoded by differentially expressed genes revealed distinctive changes in glycolysis knockout organelle metabolism. These effects were most evident in 19C CL, in which 90/239 (38%) of the genes differentially upregulated (mean fold-change >2, P-value < 0.05) in both cpEnolase and cpPGAM1A knockout lines compared to controls were predicted to possess chloroplast targeting sequences based on ASAFind (Gruber, Rocap et al. 2015) or HECTAR (Gschloessl, Guermeur et al. 2008). This was significantly greater than the proportion of genes (1,585/11,514, 14%) across the entire genome predicted to encoding chloroplast-targeted proteins that were detected in RNAseq data (one-tailed chi-squared P < 10^-05^; **Fig. 5A**; **Supplemental Dataset S5**, sheet 10). These results were supported by domain enrichment analyses, indicating significant (one-tailed chi-squared P < 0.05) enrichments in light-harvesting complex (GO:0030076), photosynthesis (GO:0009765) and protein-chromophore linkage (GO:0018298) GO terms. A more detailed resolution of gene expression patterns underpinning core organelle metabolism pathways (Ait-Mohamed, Novák Vanclová et al. 2020) suggested concerted upregulation of genes encoding light-harvesting complexes and photosynthesis machinery and plastid fatty acid synthesis machinery, alongside a probable upregulation of mitochondrial respiratory complex I and ATP synthase (**Supplemental Dataset S5**, sheets 10-11). Less dramatic changes were evident in 19C LD and 8C CL, although 13 of the 51 genes (25%) inferred to be downregulated in both cpEnolase and cpPGAM1A knockout lines under 8C CL were inferred to encode chloroplast-targeted proteins by either ASAFind or HECTAR, representing likewise an enrichment compared to all genes identified within the RNAseq data (one-tailed chi-squared P < 0.05; **Fig. 5A**).

**Fig. 5.**
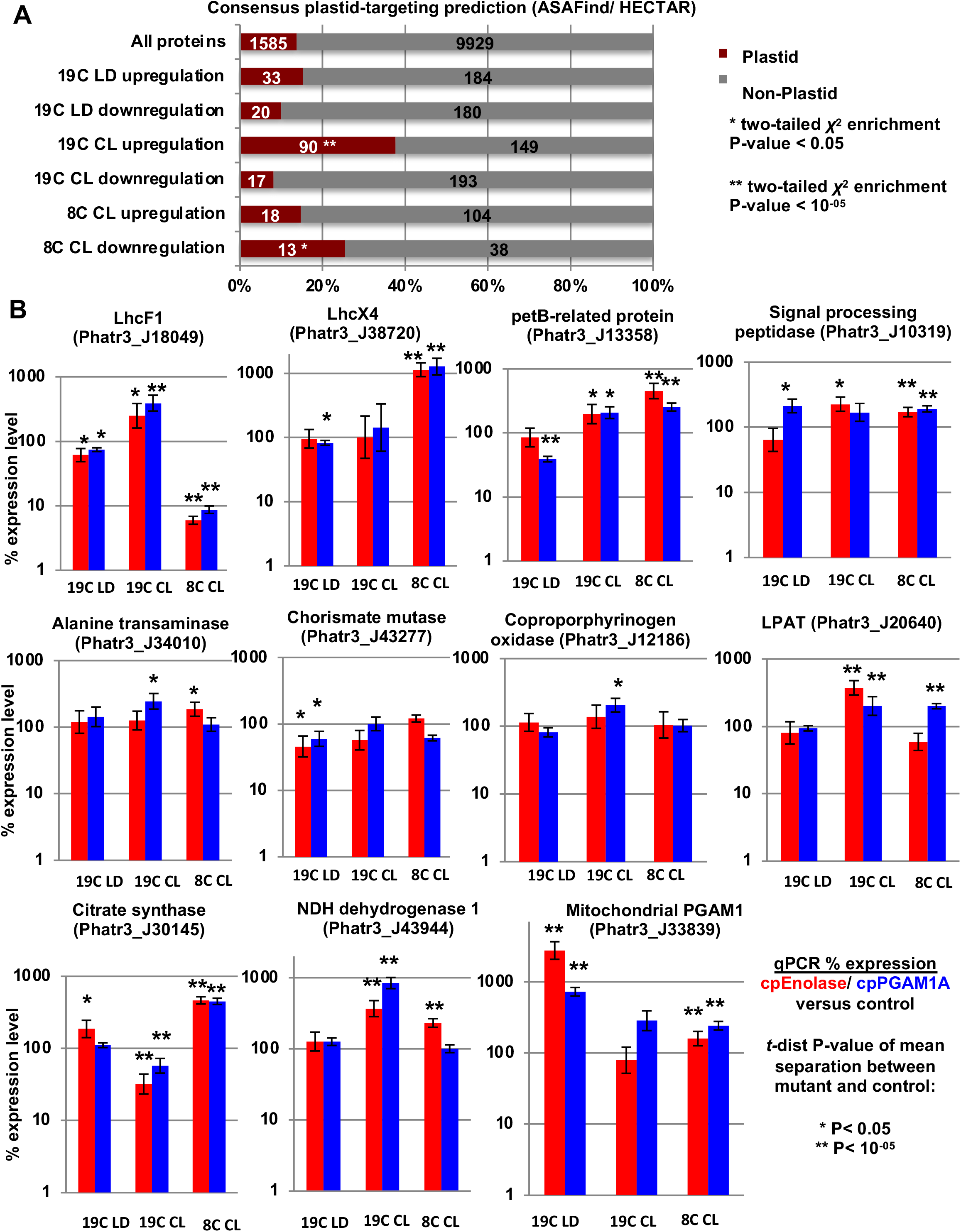
Changes in plastid and mitochondrial metabolic architecture inferred from gene expression analyses. **A:** predicted consensus localizations (either: chloroplast, or non-chloroplast) from ASAFind (Gruber, Rocap et al. 2015) and HECTAR (Gschloessl, Guermeur et al. 2008) of all genes inferred (P < 0.05, fold-change expression >2) to be up- or down-regulated in both cpEnolase and cpPGAM1A knockout compared to control lines under 19C LD, 19C CL and 8C CL. Significantly enriched localizations (two-tailed chi-squared test) are asterisked. **B:** relative mRNA abundances of eleven genes encoding exemplar chloroplast- and mitochondria-targeted proteins, verified by qRT-PCR. Genes differentially expressed (two-tailed *t*-test, P < 0.05) in each condition are asterisked.

To gain a more precise insight into the effects of plastid glycolysis-gluconeogenesis on *P. tricornutum* metabolism, we additionally validated the differential expression of eleven exemplar genes encoding chloroplast- and mitochondria-targeted proteins by qPCR in knockout and empty vector control lines across all three conditions (**Fig. 5B**; **Supplemental Dataset S5**, sheet 12). These genes showed relatively limited differences under 19C LD, limited to a slight depression in the accumulation of *Lhcf1* (Phatr3_J18049) and chorismate mutase (Phatr3_J43277) mRNA in both cpEnolase and cpPGAM1A knockouts compared to control lines (∼50% downregulation, two-tailed *t*-test P < 0.05; **Fig. 5B**). Both knockout lines over-accumulated (>600%; two-tailed *t*-test P < 10^-05^) mRNAs encoding mitochondrial phospho-glycerate mutase (Phatr3_J33839) under 19C LD compared to control lines (**Fig. 5B**).

Under 19C CL, we observed more substantial changes in plastid metabolism, including the significant (two-tailed *t*-test P < 0.05) over-accumulation of mRNAs encoding Lhcf1 (∼150%), a plastid-targeted petB-type protein presumably involved in cytochrome b_6_f metabolism (Phatr3_J13558, ∼90%), and a particularly strong over-accumulation of plastid lysophosphatidyl acyltransferase (LPAT), involved in plastid lipid synthesis (Phatr3_J20640, ∼100%, two-tailed *t*-test P < 10^-05^) in both knockout lines (**Fig. 5B**). Significant over-accumulations were also observed of mRNAs encoding plastid signal processing peptidase (Phatr3_J10319, 60-120%), alanine transaminase (Phatr3_J34010) and coporphyrinogen oxygenase (Phatr3_J12186), in either cpEnolase or cpPGAM1A knockout lines (**Fig. 5B**). Concerning mitochondrial metabolism, a strong increase (>250%, two-tailed *t*-test P < 10^-05^) was observed in mRNA for NDH dehydrogenase subunit 1 (Phatr3_J43944), involved in oxidative phosphorylation, but a corresponding decrease (>40%, two-tailed *t*-test P < 10^-05^) in mRNA for citrate synthase within the TCA cycle (Phatr3_J30145).

Finally, under 8C CL, contrasting and complementary changes were observed: up-regulation (>60%; two-tailed *t*-test P < 10^-05^) of genes encoding both the plastid signal processing peptidase and petB-related protein, and mitochondrial PGAM and citrate synthases in both knockout lines compared to controls (**Fig. 5B**). Both knockout lines were found to under-accumulate *Lhcf1* mRNA (>90%; two-tailed *t*-test P < 10^-05^), while *Lhcx4* (Phatr3_J38720), encoding a dark-expressed gene of unknown direct function but homologous to the Lhcx1 protein implicated in photoprotection (Buck, Sherman et al. 2019), was found to substantially over-accumulate in both cpEnolase and cpPGAM1A knockout lines (**Fig. 5B**).

### Metabolite profiling of Phaeodactylum cpEnolase and cpPGAM1A knockouts

Next, we considered the compound effects of cpEnolase and cpPGAM1A knockout on global metabolite accumulation under each environmental condition via GC-MS profiling of 32 sugars and amino acids (**Fig. 6**; **Fig. S16**), across 139 samples drawn from multiple knockout and control lines under 19C LD, 19C CL and 8C CL. These samples were obtained from cell pellets collected from mid-exponential phase cultures, and thus correspond to the long-term impacts on metabolite accumulation in actively growing plastid glycolysis knockout lines. Complete outputs are tabulated in **Supplemental Dataset S6**, sheets 1-2.

**Fig. 6.**
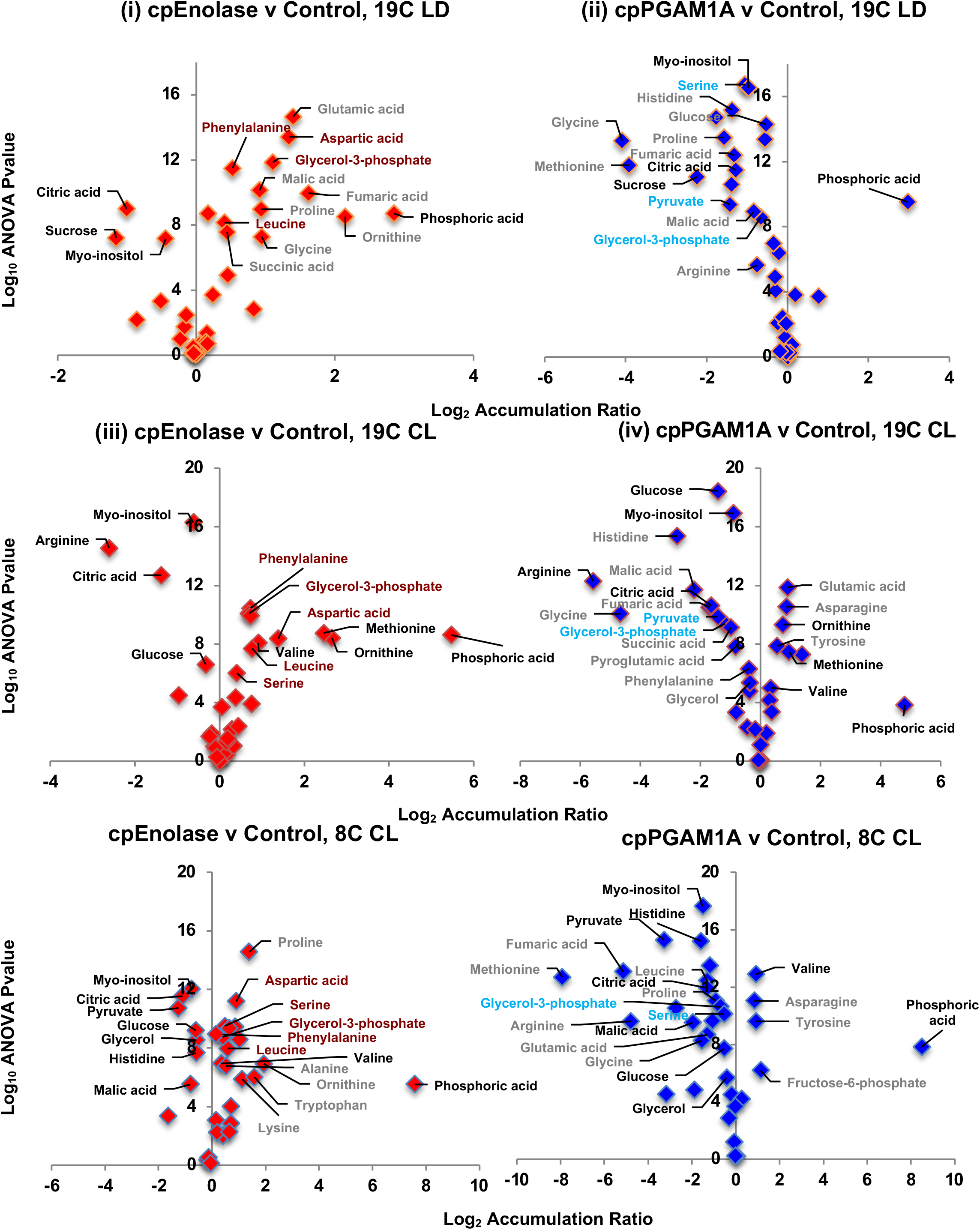
Volcano plots of differentially accumulated metabolites assessed by GC-MS. Scatterplots of the log_2_ accumulation ratios and –log_10_ P-values of difference in the mass, ribitol and quality-control-normalised abundances of 39 sugars and amino acid metabolites in cpEnolase and cpPGAM1A knockout compared to empty vector control lines, measured by GC-MS in all three experimental conditions tested. Metabolites that show a differential accumulation in each plot (P < 10^-05^) are labelled, with metabolites that show a differential accumulation in both knockout lines in each condition shown in black text, and five metabolites that are uniquely over-accumulated in cpEnolase knockout lines under all three conditions shown in dark red text.

We were unable to directly measure the accumulation of any of the products or substrates of either cpPGAM1A or cpEnolase (3-phosphoglycerate, 2-phosphoglycerate, PEP), although we detected significantly diminished (one-way ANOVA two-tailed P-value < 10^-05^) pyruvate accumulation, as a metabolite synthesised from PEP (by pyruvate kinase), in cpPGAM1A knockouts under all three conditions, and in cpEnolase knockouts under 8C CL (**Fig. 6**, **S16**). We similarly could not directly measure the accumulate of glyceraldehyde-3-phosphate (the substrate for PGAM), but could detect an overaccumulation of glycerol-3-phosphate (synthesised from glyceraldehyde-3-phosphate by glycerol-3-phosphate dehydrogenase) in cpEnolase knockout lines under all three conditions (**Fig. 6**).

In all three conditions, significant reductions (one-way ANOVA two-tailed P-value < 0.01 in both cpEnolase and cpPGAM1A knockout lines) were observed in cytoplasmic sugars and sugar derivatives (glucose, sucrose, histidine, *myo*-inositol) in cpEnolase and cpPGAM1A knockouts compared to control lines (**Fig. 6**). cpEnolase and cpPGAM1A knockout lines further under-accumulated citric acid in all three conditions, and malic acid in 8C CL (**Fig. 6**). A probable over-accumulation of phosphoric acid was observed in all knockout lines except cpPGAM1A under 19C CL (**Fig. 6**; **S17**). Significant (one-way ANOVA two-tailed P-value < 10^-05^) over-accumulations were identified for valine in cpEnolase and cpPGAM1A knockouts under 19C CL and 8C CL; for methionine and ornithine in 19C CL only; and an under-accumulation for arginine under 19C CL only (**Fig. 6**).

Finally, specific differences were observed in the metabolite accumulation patterns observed in cpEnolase and cpPGAM1A knockout lines (**Fig. 6**; **S16**). These include a significant (one-way ANOVA two-tailed P-value < 10^-05^) over-accumulation of three amino acids (aspartate, leucine and phenylalanine) and one sugar phosphate (glycerol-3-phosphate) specifically in cpEnolase knockout lines under all three conditions, and in serine under 19C CL and 8C CL only. These differences contrast to cpPGAM1A knockouts in which no significant changes were observed. Surprisingly glycerol-3-phosphate and serine were found to significantly under-accumulate under all three conditions in cpPGAM1A knockouts compared to controls **(Fig. 6; S16**).

### Lipid profiling of Phaeodactylum cpEnolase and cpPGAM1A knockouts

Next, we performed GC-MS (55 samples) and LC-MS (49 samples) of lipid profiles in multiple knockout and control lines under 19C LD, 19C CL and 8C CL. Outputs are tabulated in **Supplemental Dataset S6**, sheets 1, 3-5. While the GC-MS data project significant (one-way ANOVA two-tailed P-value < 0.05) impacts of growth condition on fatty acid profiles (e.g., a decrease of C20:5 side chain lipids balanced by an increase of C16:1 side chain lipids in 19C CL, and an over-accumulation of C16:3 side chain lipids under 19C LD, and of C18:0 side chain lipids under 8C CL), no substantial differences were observed between cpEnolase, cpPGAM1A and control lines under any conditions studied (**Supplemental Dataset S6,** sheet 3).

In contrast to the relatively limited effects on total fatty acid profiles, LC-MS analyses of lipid class distributions revealed substantial changes in lipid class distribution in plastid glycolysis-gluconeogenesis knockout lines (**Fig. 7**; **Supplemental Dataset S6,** sheet 4). Even accounting for within-line variation, both cpEnolase and cpPGAM1A knockouts were found to significantly under-accumulate triacylglycerols (TAG) (cpEnolase 3.98 ± 1.94%, cpPGAM1A 3.60 ± 1.72%, control 12.18 ± 7.26%; one-way ANOVA, two-tailed P separation of means between knockout and control lines < 0.05) and over-accumulate monogalactosyldiacylglycerols (MGDG; cpEnolase 63.83 ± 4.33%, cpPGAM1A 60.89 ± 5.64%, control 49.68 ± 8.88%; one-way ANOVA, two-tailed P< 0.05) under 19C LD (**Fig. 7A**). Further significant (P < 0.05) under-accumulations were detected in knockout lines for diacylglycerols (DAG) and sulfoquinovosyl-diacylcerols (SQDG) under 19C LD. Similar tradeoffs were observed under 19C CL, albeit with an over-accumulation, rather than under-accumulation of DAG, and an additional under-accumulation of digalactosyldiacylglycerols (DGDG), in glycolysis knockouts compared to control lines (**Fig; 6B**).

**Fig. 7.**
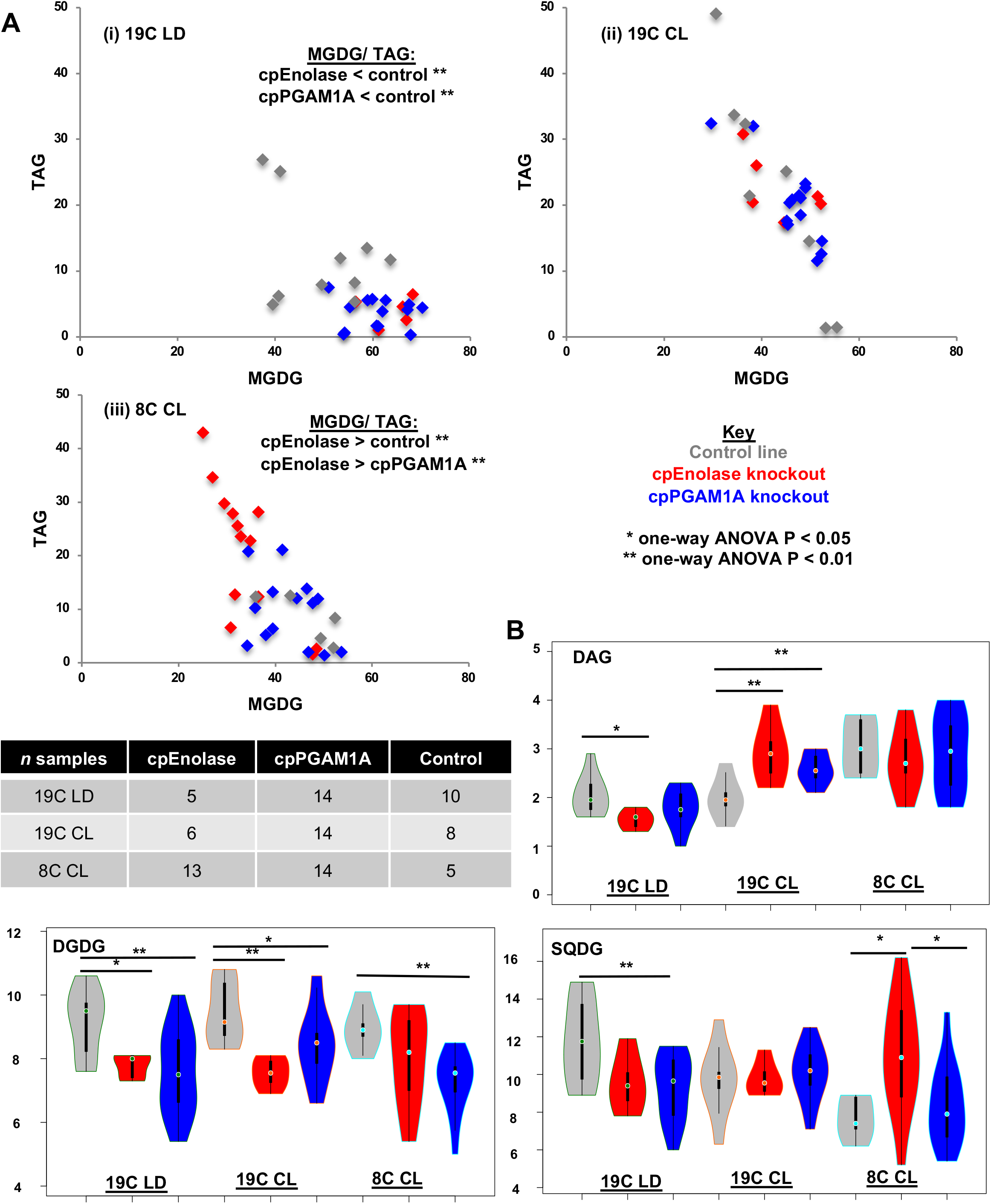
LC-MS lipid distributions in glycolysis-gluconeogenesis mutant lines. **A:** scatterplots of relative proportions of MGDG and TAG in total lipid LC-MS samples in cpEnolase and cpPGAM1A knockout lines and empty vector controls under each growth condition, showing increased MGDG: TAG in glycolysis knockout lines under 19C, and the inverse relationship in cpEnolase knockout lines only under 8C. **B**: violin plots of relative abundances of three further lipid categories inferred to differentially accumulate in glycolysis knockout lines under different growth conditions. Significant differences between knockout and control lines (one-way ANOVA) are asterisked.

Detailed analyses of the individual fatty-acid side-chains associated with different lipid classes in glycolysis knockout lines under 19C indicated increased relative contributions of C16:1 fatty acids to plastid membrane lipid *sn*-1 positions (**Supplemental Dataset S6,** sheet 5). These included conserved (P < 0.01) over-accumulations of DGDG-16-1_16-2 under 19C LD **(Fig. S18)**; and SQDG 16-1_16-0, MGDG-16-1_16-2, MGDG-16-1_16-3 and DGDG-16-1_16-1, in both cpEnolase and cpPGAM1A knockout lines under 19C CL (**Fig. S19**). A significant over-accumulation of 16-1_16-1 side chains and under-accumulation 20-5_18-4 was also observed for diacylglyceryl hydroxymethyltrimethyl-*β*-alanine (DGTA), a betaine lipid known to act as a platform for the biosynthesis of 20:5 fatty acids, in both cpEnolase and cpPGAM1A knockout lines under 19C LD (**Fig. S18**) (Dolch and Maréchal 2015, Popko, Herrfurth et al. 2016).

Under 8C CL, quite different trends were observed in fatty acid accumulation in cpEnolase knockouts compared to cpPGAM1A knockouts and controls. These correlated principally with an over-accumulation of TAG (cpEnolase 20.88 ± 12.21%, cpPGAM1A 9.62 ± 6.31%, control 8.15 ± 3.95%; one-way ANOVA, two-tailed P < 0.05) in lieu of MGDG (cpEnolase 34.20 ± 6.74%, cpPGAM1A 42.94 ± 6.01%, control 46.61.3 ± 6.25%; one-way ANOVA, two-tailed P < 0.5; **Fig. 7A**). An over-accumulation of SQDG was observed in both cpEnolase and cpPGAM1A knockouts compared to controls, albeit with greater severity in cpEnolase knockouts (**Fig. 7B**). Considering side-chain distributions of individual lipid classes, a significant (one-way ANOVA two-tailed P-value < 0.01) over-accumulation of short-chain (C14:0, C16:1) and *sn*-1 and *sn-2* fatty acids was observed in cpEnolase knockouts (**Fig. S20A**). A probable exchange of very long-chain *sn*-2 fatty acids in SQDG pools was further observed in cpEnolase knockouts, with significant (one-way ANOVA two-tailed P-value < 0.01) increases in SQDG 14-0_16-0 and SQDG-14_0-16-1 in lieu of SQDG-16-2_24-0 in cpEnolase knockouts compared to cpPGAM1A and control lines (**Fig. S20B**; **Supplemental Dataset S6**, sheet 5).

### Reaction kinetics of expressed copies of Phaeodactylum cpEnolase and cpPGAM1A

Finally, we assessed the kinetics of cpPGAM and cpEnolase in both glycolytic and gluconeogenic directions. Previous studies (e.g., in animal renal and liver tissue) project reversible reaction kinetics for both enolase and PGAM enzymes. The reaction rates of enolase and PGAM show limited difference in glycolytic versus gluconeogenic directions *in vivo*, with measured enolase rates in rat kidney tissue equivalent to approximately 14,000 µmol g dry weight^-1^ hr^-1^ in the glycolytic direction, and 20,000 µmol g dry weight^-1^ hr^-1^ in the gluconeogenic direction (KREBS 1963, Scrutton and Utter 1968, Reinoso, Telfer et al. 1997). Equally, purified enolase and PGAM typically show greater affinity for 3-PGA than PEP, with a 5-8 fold difference in Km measured in mammalian, yeast and *Trypanosoma brucei* enzymes (Rider and Taylor, 1974; Hannaert et al., 2003).

Using a previously defined assay (Sutherland, Posternak et al. 1949, Zhang, Sampathkumar et al. 2020) with modified versions of each protein (codon-optimised, and lacking signal peptides) expressed in *E. coli,* alongside measured NADH consumption coupled to either lactate dehydrogenase (glycolysis) or glyceraldehyde-3-phosphate dehydrogenase (**Fig. S20**). Both enzymes were inferred to possess reversible reaction kinetics, metabolizing NADH when supplied both with 3-PGA (in the glycolytic direction) and PEP (in the gluconeogenic direction; **Fig. 8**, **Fig. S20**).

**Fig. 8.**
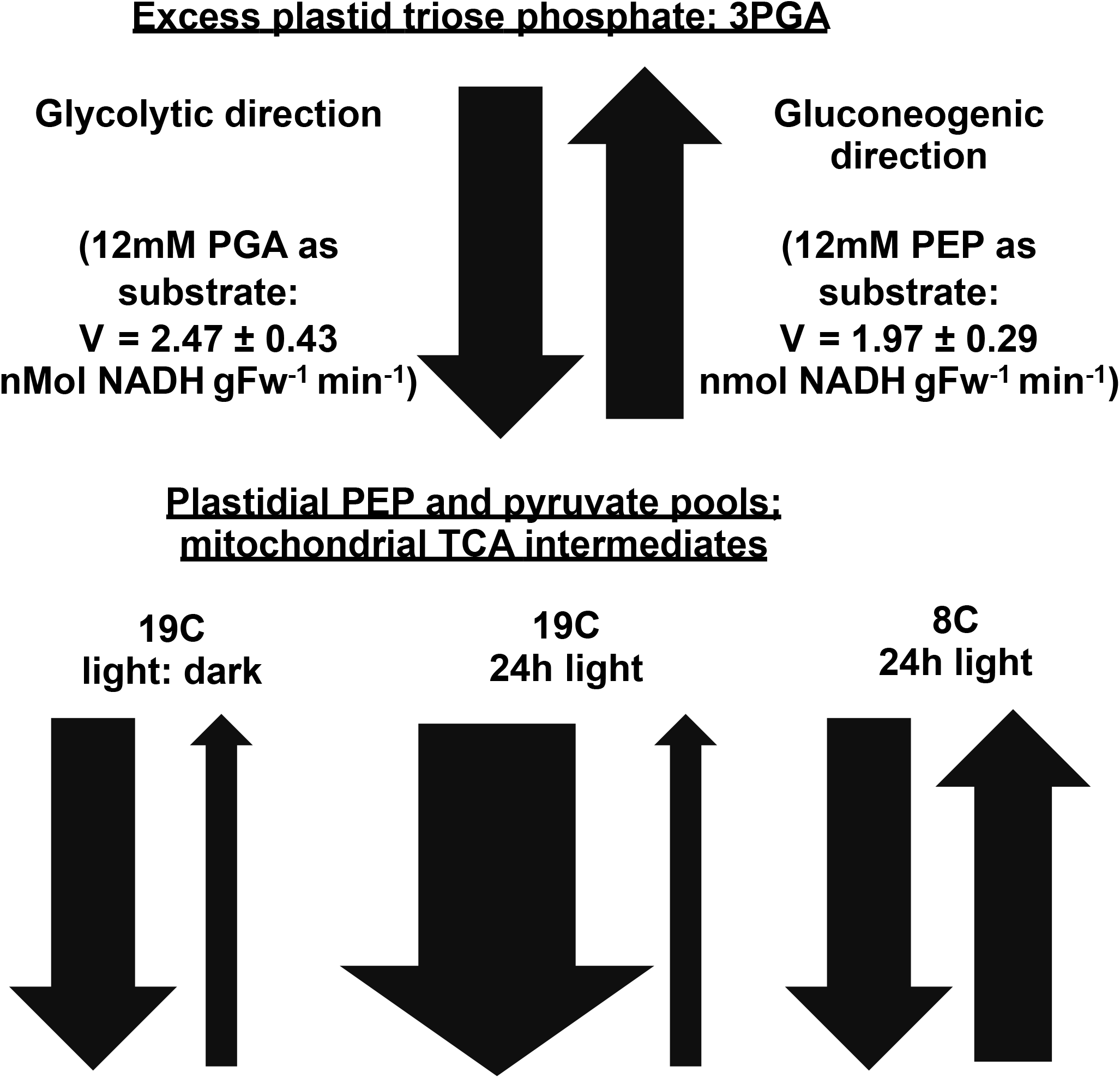
Proposed activities of *P. tricornutum* plastid lower glycolysis-gluconeogenesis. Schematic diagram showing potential inferred roles of lower half diatom plastid glycolysis-gluconeogenesis in each environmental condition tested. The measured V_max_ of purified cpEnolase and cPGAM1A supplemented with 3-PGA (glycolytic direction) or PEP (gluconeogenic direction) are provided for 9mM substrate in each case.

The measured reaction rates were effectively reversible, albeit marginally greater in the glycolytic than gluconeogenic direction (e.g., 2.47 ± 0.43 versus 1.97 ± 0.29 nMol NADH consumption per gram free weight purified enzyme per minute when supplied with12 mM 3PGA or 12 mM PEP; **Fig. 8**).

## Discussion

We characterise a lower glycolytic-gluconeogenic pathway associated with diatom plastids, relating specifically to two plastid-targeted proteins, cpEnolase and cpPGAM1A, and focusing on the model species *P. tricornutum*. Our data position plastid glycolysis-gluconeogenesis as arising in a recent ancestor of diatoms and their closest relatives (e.g., pelagophytes, dictyochophytes) (Nonoyama, Kazamia et al. 2019). The presence of plastid glycolysis in haptophytes may be as a result of endosymbiotic transfers into this group from a pelagophyte/ dictyochophyte-related alga, as suggested in previous studies (Dorrell, Gile et al. 2017, Jiang, Cao et al. 2023). We further show that plastidial lower glycolysis-gluconeogenesis has a limited distribution across the algal tree of life, with no examples in primary red and few in primary green algae (**Fig. S1, S12**). It is possible that the occurrence of organelle-targeted isoforms of these enzymes is underestimated, e.g., due to lower sensitivity of diatom and plant-trained targeting predictors on other algal groups (Fuss, Liegmann et al. 2013, Gruber, Rocap et al. 2015). We propose that diatom plastid glycolysis most likely originated through the duplication and retargeting of mitochondrial respiratory enzymes (**Fig. 1**).

Using meta-genomic data from *Tara* Oceans we demonstrate that diatom plastid glycolysis is likely highly expressed at high latitudes (**Figs. 3**, **S8-S11**), which are subject to extreme photoperiods and low temperature. These data are further supported by collection sites of cultured species, with no occurrences of cultured diatoms lacking plastid-targeted PGAM enzymes beyond 50°N (**Fig. S1B**), and evidence from meta-genome assembled genomes, in which diatoms that possess apparent plastid-targeted Enolase and PGAM enzymes show an associative preference for high latitudes, while those that lack them do not (**Fig. S12**). These enrichments appear to be largely specific to diatoms, with polar circle haptophytes, cryptomonads and other ochrophytes lacking apparent plastidial glycolysis found further than 60°N and 70°S, considering both cultured species and MAG data, although with a potential parallel recruitment of plastidial glycolysis to isolates of the prasinophyte genus *Micromonas* abundant in high latitude *Tara* stations (Lovejoy, Vincent et al. 2007). We thus tentatively propose that lower half plastid glycolysis correlates to diatom occurrence in high latitude *Tara* Oceans stations, with diatoms that lack identifiable copies of these proteins absent from these stations, and other algal groups (except potentially chlorophytes) showing no preference for plastidial glycolysis at high latitudes.

We are hesitant to state that plastid glycolysis is an adaptive feature of diatoms towards high latitudes, given it apparently originated in a common ancestor of diatoms and several other algal groups (i.e., pelagophytes, and dictyochophytes), and is retained in species such as *Phaeodactylum* which is typically associated with intermediate latitudes (Rastogi, Vieira et al. 2020). cpEnolase and cpPGAM cannot therefore have been viewed to have been gained in specific diatom species in response to environmental selection. An open question, particularly given the largely latitude-insensitive distributions to diatom MAGs lacking plastid glycolysis, remains to what extent diatoms that have secondarily lost their plastid glycolytic pathway are abundant in nature (**Fig. S12**). Ultimately, the physiological functions of diatom plastid glycolysis will be best identified through competition assays e.g. between diatom species with different plastid carbon metabolism arrangements, or between *Phaeodactylum* knockout and empty vector control lines under each condition (Siegel, Baker et al. 2020).

Nonetheless, from our analysis of published *Phaeodactylum* transcriptome data and qRT-PCRs, we note that both cpEnolase and cpPGAM1A genes are transcriptionally induced in response to both long day conditions and low temperatures (**Fig. 2B**, **S7**). These biases are particularly interesting given the growth analysis of *P. tricornutum* knockout lines. In particular, we observe more intense growth defects in *Phaeodactylum* lines under continuous illumination than in light: dark cycles (**Figs. 4**, **S9**), which alongside gene expression data suggests increased importance of plastid glycolysis in diatoms subject to long days. In contrast, under low temperatures no difference was observed in the growth rate of glycolysis knockouts showed to control lines (**Fig. 4**). We note that the relative expression of cpEnolase and cpPGAM1A are even greater at 8C than 19C under continuous light conditions, and it is possible that flux occurs through plastid glycolysis at low temperature despite the absence of a clear growth phenotype in knockout versus control lines. We thus tentatively propose that lower glycolysis-gluconeogenesis may have multiple functions in the diatom plastid, with different functions dependent on both light and temperature conditions.

Considering the observed phenotypes of knockout and control lines (**Figs. 4-7**; **S14-S19**) and the reversible kinetics of expressed enzymes, we suggest potential functions contributed by the lower half of plastid glycolysis-gluconeogenesis in diatoms under 19C LD, 19C CL and 8C CL conditions (**Fig. 8**). Overall, our suggested roles for cpEnolase and cpPGAM1A are predominantly in favour of metabolic flux in the glycolytic direction, reflecting the underaccumulation of pyruvate in cpPGAM1A knockouts, and overaccumulation of 3-phospho-glycerate in cpEnolase knockouts (**Fig. 6**). We also present this hypothesis based on the innate metabolic activity of the Calvin cycle, which is likely to yield a high relative abundance of triose phosphate in the plastid under illuminated and photosynthetically active conditions; although we note that studied diatom triose phosphate transporters show higher transport affinity for PEP than DHAP, which may facilitate substrate supply for gluconeogenic activity (Moog, Nozawa et al. 2020) (**Fig. 8**). These results are nonetheless inferential based on the long-term accumulation patterns of stable metabolites and the expression of implicated metabolic genes. Whilst these would be more effectively validated via direct flux measurements, e.g., comparative ^13^C-glycerol or -glucose labelling of glycolysis knockout and control lines (Zheng, Quinn et al. 2013, Huang, Liu et al. 2015), this was beyond the scope of the current study.

Under 19C LD, we observe limited gene expression changes in cpPGAM1A and cpEnolase knockout lines, except (as inferred from qPCR) a downregulation in plastid chorismate mutase, which forms part of the plastid shikimate pathway, that typically consumes PEP (Bromke 2013) and may form a primary acceptor of glycolytic products (**Fig. 5B**). One of the products of chorismate mutase activity, phenylalanine, does seem to overaccumulate in metabolite pools of cpEnolase mutants only under these conditions, pointing to potentially more complex fluxes (discussed below). We also note an upregulation of mitochondrial PGAM in both lines (**Fig. 5B**), which might relate to a greater level of mitochondrial glycolysis, e.g. of exported plastid glyceraldehyde-3-phosphate in the knockout lines. Strikingly, both mutant lines underaccumulate TCA cycle intermediates (citric acid in both lines, and fumaric and malic acid in cpPGAM1A only) which may suggest less retention of metabolised sugar in the mitochondrion. Finally, both mutants also underaccumulate sugars typically synthesised in the cytosol (sucrose, myo-inositol) which we interpret to imply less excess fixed carbon in the knockout compared to control lines (**Fig. 6**). Overall these data seem to point to less efficient carbon usage, and an overall redirection of plastidial triose phosphate from plastid or cytoplasmic anabolic reactions to mitochondrial catabolism in knockout lines.

We also note some evidence for lipid remodelling in glycolysis mutant lines. These include a relative over-accumulation of galactolipids in lieu of TAGs, and short-chain fatty acids in lieu of longer equivalents (**Figs. 7**, **S18**). Previous studies have noted the importance of lipid metabolism in diatom stress responses (Zulu, Zienkiewicz et al. 2018), and that most or all diatom lipid synthesis occurs directly in the plastid (Huang, Pan et al. 2024). Many of the metabolic reactions required for lipid synthesis, including acyl-coA synthesis from pyruvate (Maréchal and Lupette 2020), glycerol-3-phosphate from glyceraldehyde-3-phosphate (Kroth, Chiovitti et al. 2008), and glucosyl-1-phosphate from cytoplasmic glucosyl-1-phosphate (Zhu, Shi et al. 2016), are likely to be Impacted by plastid carbon metabolism. Specifically, an underaccumulation of TAGs and fatty acids in lieu of galactolipids would also suggest a lower ratio of pyruvate (for acyl-coA synthesis) to glyceraldehyde-3-phosphate (for galactosyl-phosphate synthesis) in glycolysis knockout lines (Demé, Cataye et al. 2014), similarly to the underaccumulation of pyruvate observed in our metabolomic data (**Fig. 6**). We infer that these changes are probably driven by substrate limitation, as we observe no changes in the transcription of genes involved in fatty acid synthesis in glycolysis knockout lines; nor do we increased expression of cpEnolase or cpPGAM1A in cellular conditions (N and P limitation) known to induce diatom lipid accumulation (**Fig. 5B**; **Fig. S7**) (Abida, Dolch et al. 2015)

Under 19C CL, we observed much more dramatic remodelling of cellular transcription in knockout lines compared to controls (**Fig. 5A**). These are notably concordant with greater expression of cpEnolase and cpPGAM1A in wild-type cells (**Fig. 2B**), and an enhanced growth defect in knockout lines, together suggesting greater potential flux through this pathway than under 19C LD (**Fig. 4B**). The transcriptional changes include greater overall photosynthesis gene expression, e.g., *Lhcf1* (**Fig. 5B**), which was corroborated in photo-physiological analyses by larger PSII antenna size, i.e., a larger functional cross-section (σPSII) (**Fig. S15**). It should be noted that the increase in PSII antenna size does not necessarily change the quantum yield of individual PSII reaction centres, and therefore the increased σPSII is independent of the Fv/ Fm measured, which remains equivalent between knockout and control lines (**Fig. S15**). We did not observe consistent differences in the expression of nitrogen or phosphorus stress metabolism, or in the expression of the *P. tricornutum* biophysical carbon concentration mechanisms of knockout lines, suggesting that these differences were not caused by N, P or CO_2_ limitation in the control lines (**Supplemental Dataset S5,** sheets 4-5) (McCarthy, Smith et al. 2017, Nawaly, Matsui et al. 2023). We further did not measure differences in photosynthetic performance (electron transport), or an upregulation of genes encoding proteins involved in photoprotection, e.g., LhcX family or xanthophyll cycle enzymes in knockout lines under 19C CL (**Fig. 5B**; **Fig. S15**; **Supplemental Dataset S4,** sheet 12) (Buck, Sherman et al. 2019, Bai, Cao et al. 2022), suggesting that the differential expression of photosynthesis genes in the knockout lines does not directly influence photosynthesis.

In contrast, from RNAseq and qRPCR we observed an upregulation of multiple mitochondrial NDH dehydrogenase and ATP synthase subunits; and downregulation of TCA cycle enzymes in glycolysis knockout lines (**Fig. 5B**; **Supplemental Dataset S5,** sheet 12). Our metabolomic data further show an underaccumulation of citric acid, as per in 19LD conditions; but also arginine, synthesised in the diatom mitochondria from glutamate and aspartate in the urea cycle (Allen, Dupont et al. 2011, Bromke 2013). We globally interpret these phenotypes to mean an increase in mitochondrial respiratory electron transport in glycolysis knockout lines without necessarily an increase in mitochondrial primary metabolic activity. Previous studies have noted the important role of diatom mitochondria in dissipating excess plastid reducing potential (Bromke 2013, Bailleul, Berne et al. 2015, Broddrick, Du et al. 2019), and we wonder if these phenotypes observed in knockout lines under 19C CL conditions relate to the respiratory dissipation of plastidial NADPH.

It remains to be determined what routes beyond plastidial glycolysis contribute substrates (e.g. PEP, pyruvate) to the *P. tricornutum* pyruvate hub. Previous studies have noted that diatom plastid triose phosphate transporters may be able to transport PEP directly from the cytoplasm, and one of these (Phatr3_J54017) is indeed upregulated in both cpEnolase and cpPGAM1A knockout lines under 19C CL (**Supplemental Dataset S5**, sheet 3) (Moog, Nozawa et al. 2020, Liu, Storti et al. 2022). Elsewhere our data suggest that amino acids may modify the concentrations of *P. tricornutum* plastid PEP and/or pyruvate. These data include the overexpression of plastid alanine transaminase in cpPGAM1A knockouts (**Fig. 5B**); and the overaccumulation in both knockout lines of amino acids synthesised either from pyruvate (valine), PEP (aspartate via PEP carboxylase, and methionine from aspartate), or more broadly involved in plastid amino acid recycling (ornithine and glutamate, in the diatom plastid ornithine cycle) (Levering, Broddrick et al. 2016, Smith, Dupont et al. 2019, Yu, Nakajima et al. 2022) (**Fig. 6**). The direction (import or export) and significance of these amino acid fluxes will be best determined e.g. with an inducible knockout mutant compromised for both plastidial glycolysis and amino acid incorporation (e.g., via nitrate reductase) to allow direct metabolic quantification of *de novo* synthesised amino acids (McCarthy, Smith et al. 2017, Yin and Hu 2023).

Under 8C CL, we identify an over-accumulation of mRNAs encoding plastid biogenesis and mitochondrial glycolytic proteins, an over-accumulation of short-chain amino acids (valine) and an under-accumulation of cytoplasmic sugars and amino acids (glucose, histidine) in cpEnolase and cpPGAM1A knockouts relative to controls (**Figs. 5B, 5**). We further note under-accumulations of pyruvate in both knockout lines (**Fig. 6**). Knockout lines under 8C conditions, however, have additional phenotypes not observed at 19C. These include an overall enrichment in down-regulated genes encoding plastid-targeted proteins (**Fig. 5A**); and a specific over-accumulation of TCA cycle (citrate synthase) and a possible non-photochemical quenching-associated mRNA (*LhcX4*) (**Fig. 5B**) (Bailleul, Berne et al. 2015, Murik, Tirichine et al. 2019). Finally, specific differences are observed between cpEnolase and cpPGAM1A knockout lines at 8C. These include an overaccumulation of TAGs and SGDQs over glucosyl-lipids, and the over-accumulation of aspartate and phenylalanine in cpEnolase knockouts only (**Figs. 6**, **7; S17**).

The more complex phenotypes observed in our knockout lines at 8C, and particularly the differences between cpPGAM1A and cpEnolase knockouts, may be due to several reasons. PGAM is typically viewed to have a greater catalytic activity than Enolase from biochemical studies, although this may be compensated by the greater relative abundance of cpEnolase than cpPGAM1A in *Phaeodactylum* plastids (**Fig. S6A**) (Scrutton and Utter 1968, Huang, Pan et al. 2024). It is true that cpPGAM1A may be compensated by functionally redundant proteins (cpPGAM1B, cpPGAM2) in the *Phaeodactylum* plastid in knockout lines, whereas cpEnolase is functionally non-redundant, but none of these genes are specifically induced in RNAseq data of cpPGAM1A knockouts under 8C CL (**Supplemental Dataset S5**, sheet 3) suggesting an absent of specific compensation in the cpPGAM1A mutant line. We note that several of the phenotypes associated with 8C CL, and with cpEnolase knockouts specifically, relate to accumulation either of acyl-coA (TAGs) or PEP (aspartate, phenylalanine), which may suggest impediment of the gluconeogenic rather than glycolytic activity of cpEnolase and cpPGAM. The reversibility of the cpPGAM1A and cpEnolase reaction is confirmed by enzymatic data (**Fig. 8**), and it remains to be determined to what extent these enzymes function bidirectionally *in vivo*. It also remains to be determined how these functions impact on growth kinetics and viability of diatoms in the wild, given the limited differences in growth rate observed in the lab between knockout and control lines (**Fig. 4**).

The complex phenotypes for diatom plastid glycolysis inferred from environmental and experimental data contrast with those for plant plastid glycolysis, with (for example) *A. thaliana* cpEnolase and cpPGAM mutants presenting relatively limited phenotypes (Prabhakar, Löttgert et al. 2009, Andriotis, Kruger et al. 2010). We note that the cytoplasmic and respiratory plant Enolase and PGAM1 isoforms, alongside having predominant impacts on plant carbon flux, also have important moonlighting roles in plant development, immune responses and even in the structural coordination of plastids and mitochondria (Zhao and Assmann 2011, Zhang, Sampathkumar et al. 2020, Yang, Wang et al. 2022). We similarly anticipate that further surprises will be identified for the functions of diatom plastid glycolysis, and for this still poorly understood pathway in the photosynthetic tree of life.

## Materials and Methods

### Culture conditions

*Phaeodactylum tricornutum* strain Pt1.86 was grown in enhanced seawater (ESAW) medium supplemented with vitamins, but without silicon or added antibiotics, in 50 μE m^-2^ s^-1^ white light. Light profiles were measured with a SpectraPen photofluorometer (Photon Systems Instruments, Czech Republic); and are provided in **Supplemental Dataset S4**, sheet 13. Cultures were grown under one of four light, temperature and shaking regimes. For general molecular work and transformation, cultures were grown under 19 °C with 12h light: 12 dark cycling, shaken at 100 rpm (for general molecular work and transformation), following the established methodology of Falciatore et al. (Falciatore, Casotti et al. 1999). For comparative physiology work, we were unable to replicate shaking conditions at low temperatures, and therefore chose to use conditions without shaking: 19 °C with 12 h light: 12 h dark cycling (« LD » growth conditions and physiological analysis); 19 °C with 24 h continuous light and without shaking (« CL » growth conditions and physiological analysis); or 8°C with 24h continuous light and without shaking (« 8C » growth conditions and physiological analysis). All cultures achieved measured mid exponential Fv/Fm values of > 0.6, suggesting that the absence of shaking did not impact on photosynthetic efficiency (**Supplemental Dataset S5**, sheet 8).

Batch culturing of *P. tricornutum* for genetic manipulation was performed under fluorescent lamps. Physiological experiments were principally performed at 19°C in an AlgaeTron AG230 (Photon Systems Instruments) with cool white LED (WIR) illumination, and technical specifications described in https://growth-chambers.com/data/algaetron-ag-230/download/AlgaeTron_AG_230_Manual2021-finalweb.pdf. Growth experiments were performed at 8°C using a low-temperature adapted cool white LED (WIR, ECCLIM). Details of all three spectra used, as measured with a SpectraPen (PSI), are provided in **Table S4**, sheet 13.

Mutant *P. tricornutum* lines were maintained on ½ ESAW 1% agarose plates, supplemented by 100 μg ml^-1^ each ampicillin and streptomycin, and 30 μg ml^-1^ chloramphenicol, and either 100 μg ml^-1^ zeocin (single transformants), or 100 μg ml^-1^ zeocin and 4 μg ml^-1^ blasticidin (complementation lines), as previously described (Falciatore, Casotti et al. 1999, Buck, Río Bártulos et al. 2018). All functional analyses of transformant lines were performed on transformant lines grown in the absence of antibiotic selection, to avoid secondary effects on growth or physiology.

### Phylogenetic identification of plastid lower half glycolysis-gluconeogenesis enzymes

Plastid-targeted glycolysis lower half enzymes were searched across 1,673 plant and algal genomes and transcriptomes (**Dataset S1**, sheet 1). Briefly, this involved searching all annotated *P. tricornutum* PGAM (Phatr3_J17086, Phatr3_J51404, Phatr3_J5605, Phatr3_J5629, Phatr3_J8982, Phatr3_J37201, Phatr3_J47096) and enolase (Phatr3_draftJ1192, Phatr3_draftJ1572, Phatr3_J41515) peptide sequences with BLASTp and a threshold e-value of 10^-05^, and a reciprocal BLASTp with criteria - max_target_seqs 1 to retrieve the best homologues against the entire *P. tricornutum* genome. For PGAM, where *P. tricornutum* queries failed to retrieve homologues in >50% searched libraries, a second BLASTp was performed with query peptide sequences from *A. thaliana* (AT2G17280, AT1G09780, AT3G05170, AT3G08590, AT3G50520, AT5G04120, AT5G64460), and a reciprocal BLASTp was performed with the *P. tricornutum* genome supplemented with these sequences. Similar approaches were subsequently used to identify equivalent plastid glycolysis proteins from *Tara* Oceans meta-genome assembled genomes (MAGs), as assembled in (Delmont, Gaia et al. 2022).

The domain content of each potential homologue was identified using hmmscan and the version 33.1 Pfam database (Mistry, Chuguransky et al. 2020). Only Enolase sequences that contained >90% predicted domain coverage to both Enolase_N and Enolase_C domains; and PGAM sequences that contained >50% domain coverage to the His_Phos domain (based on the corresponding coverage observed in *P. tricornutum* sequences) were viewed as being complete. Sequences for which the N-terminus of the region homologous to the PFAM domain was located within the first 20 aa of the predicted sequence (i.e., less than the length of a typical plastid-targeting sequence) (Emanuelsson, Brunak et al. 2007) were viewed as lacking credible targeting sequences. All remaining proteins were scanned, considering both complete proteins and sequences trimmed to the first encoded N-terminal methionine, using targetp (using a plant scoring matrix) (Emanuelsson, Brunak et al. 2007), PredAlgo (Tardif, Atteia et al. 2012), HECTAR (Gschloessl, Guermeur et al. 2008) and ASAFind (with SignalP 5.0) (Gruber, Rocap et al. 2015, Almagro Armenteros, Tsirigos et al. 2019). Sequences from primary plastid-containing organisms (plants, green and red algae, glaucophytes) that were inferred to possess a plastid-targeting sequence either with TargetP or PredAlgo, and sequences from secondary plastid-containing organisms that were inferred to possess a plastid-targeting sequence with either HECTAR or ASAFind, considering both complete and N-trimmed sequence models, were annotated as putatively plastid-targeted.

A more detailed phylogenetic analysis was performed using Enolase and PGAM homologues obtained from a subset of 289 complete cryptomonad, haptophyte and stramenopile genomes and transcriptomes in the above library, alongside homologues identified from a further 85 prokaryotic and eukaryotic genomes sampled with taxonomic balance from across the remainder of the tree of life (Liu, Storti et al. 2022). Sequences were also screened for mitochondrial presequences using HECTAR (Gschloessl, Guermeur et al. 2008), and MitoFates, run with threshold value 0.35 (Fukasawa, Tsuji et al. 2015).

The pooled set of sequences were aligned first with MAFFT v 7.0 under the --auto setting, followed by the in-built alignment programme in GeneIOUS v 10.0.9, under default settings (Kearse, Moir et al. 2012, Katoh, Rozewicki et al. 2017). Incomplete and poorly aligned sequences, alongside taxonomically uninformative N- and C-terminal regions were removed from the alignment manually, followed by trimal with setting –gt 0.5 (Capella-Gutiérrez, Silla-Martínez et al. 2009). Phylogenetic analyses were performed with MrBayes v 3.2 and rAxML v 8, integrated into the CIPRES webserver (Stamatakis 2014, Miller, Schwartz et al. 2015). MrBayes trees were run for 10,000,000 generations with the GTR, Jones and WAG substitution matrices, 4 starting chains and sumt and sump burnin fractions set to -0.5; all cold chains were confirmed to have reached a pvalue plateau below 0.1 prior to the start of the consensus building. rAxML trees were run with GTR, JTT and WAG substitution matrices, 350-400 ML generations, and automatic bootstopping. Phylogenies were either rooted between bacterial and eukaryotic sequences (Enolase); or on the mid-point (PGAM1, PGAM2) due to the absence of a single monophyletic bacterial outgroup. A summary of these analyses is provided in **Supplemental Dataset S1**.

### Analysis of previously published *Phaeodactylum* data

First, the mean relative abundance of peptides corresponding to Enolase and PGAM sequences was retrieved from published mass spectrometry data of *Phaeodactylum* plastid-enriched fractions and total cell pellets, following (Huang, Pan et al. 2024).

The two datasets were found to show a positive correlation with one another (*r*= 0.891, n = 901, P < 10^-05^) considering all proteins recovered in the plastid-associated fraction with a previously suspected plastid localisation (plastid-encoded, or plastid-targeted nucleus-encoded proteins inferred using combined ASAFind and HECTAR predictions). These data are provided in **Supplemental Dataset S2**, sheet 1.

Next, the expression trends of *Phaeodactylum* plastid glycolysis proteins (cpEnolase, cpPGAM1A, PGAM2) were assessed in combat normalised RNAseq data (Ait-Mohamed, Novák Vanclová et al. 2020) assembled from three prior studies, relating to induced nitrate limitation (nitrate reductase knockout), phosphate limitation and resupply, and iron limitation over a Circadian cycle (Cruz de Carvalho, Sun et al. 2016, Smith, Gillard et al. 2016, McCarthy, Smith et al. 2017). A second set of comparisons were performed using normalised *Phaeodactylum* microarray data (summarised in (Ashworth, Turkarslan et al. 2016), particularly relating to changes in light quality, wavelength and Circadian time point. To enable global analyses of coregulation to cpEnolase and cpPGAM enzymes, these two datasets were converted into ranked values (i.e., for Spearman correlation) and merged, following (Liu, Storti et al. 2022). These data are summarised in **Supplemental Dataset S2**, sheets 2-4.

### *Tara* Oceans Analysis

The complete *Tara* Oceans and *Tara* Oceans Polar Circle libraries of meta-genome and meta-transcriptome diversity (Carradec, Pelletier et al. 2018, Royo-Llonch, Sánchez et al. 2020) were searched for orthologues of diatom cpEnolase, cpPGAM1A and PGAM2 sequences via a phylogenetic reconciliation approach benchmarked in previous studies (Kazamia, Sutak et al. 2018, Liu, Storti et al. 2022). This approach uses the combined outputs of hmmer, BLAST best-hit, and single-gene tree topologies to only retain *Tara* Oceans meta-genes that reconcile as monophyletic with a defined query set, in these case plastid-targeted diatom isoforms of each enzyme. Exemplar tree topologies are shown in **Fig. S8.**

First, a HMM (hidden Markov model) was constructed for all diatom plastid-targeted sequences in the untrimmed alignments for each phylogeny, as detailed above, and searched into the complete *Tara* Oceans catalog by hmmer (http://hmmer.org) with evalue 10^-10^ to identify putative meta-gene homologues of each protein. Matching sequences were extracted, and searched by BLASTp against the complete copy of the *P. tricornutum* genome (Rastogi, Maheswari et al. 2018). Only sequences that retrieved a best hit against an Enolase or PGAM sequence (and therefore likely correspond to homologues of each protein) were retained. Next, the retained sequences were similarly searched by BLASTp against the complete untrimmed alignment of cultured Enolase and PGAM sequences. Only sequences that retrieved a diatom plastid-targeted isoform were retained, allowing the elimination of non-diatom and homologues of diatom non-plastid sequences. Finally, sequences were combined with the untrimmed alignment of cultured sequences from each gene and realigned using the same MAFFT, GeneIOUS and trimal pipeline as defined above. Curated alignments were screened by rAxML tree with the JTT substitution matrix, as above. Only *Tara* Oceans sequences that resolved within a monophyletic clade with diatom plastid-targeted proteins, defined as all sequences that position closer on a midpoint rooting of the tree to diatom plastid-targeted proteins than to any non-diatom or non-plastid targeted sequences, were extracted for further analyses.

Relative abundances were calculated for the total occurrence of all phylogenetically verified diatom plastid-targeted proteins in both meta-transcriptome and meta-genome data. Relative expression levels of each gene were estimated by reconciling the calculated meta-transcriptome abundances either to total diatom meta-transcriptome sequences using the formula 10^^6^(Σ_metaT_/ Σ_DiatomT_), i.e., expressed per million reconciled diatom reads, or to calculated meta-genome abundances, using the formula and log_10_ (1+ Σ_metaT_) -log_10_ (1+ Σ_metaG_), to allow inclusion of zero metaG values. Pearson and Spearman correlations were calculated between relative abundances and all quantitative measured environmental variables associated with *Tara* Oceans samples as stored within the PANGAEA repository (Pesant, Not et al. 2015). All calculations were repeated independently for each depth (surface, or deep chlorophyll maximum/ DCM) and size fraction (0.8-2000 μm, 0.8-5 μm, 3/5-20 μm, 20-180 μm, and 180-2000 μm), with 3 and 5 μm filters viewed as equivalent to allow reconciliation of Arctic and non-Arctic data, respectively. All *Tara* Oceans meta-gene assignations, alongside individual and total abundance calculations are provided in **Supplemental Dataset S3**, sheets 1-10.

*Tara* Oceans MAGs were partitioned into those that contained credible chloroplast-targeted copies of Enolase and/ or PGAM sequences, using a similar reciprocal BLAST best-hit, PFAM analysis and *in silico* targeting prediction pipeline as used for cultured species data (**Supplemental Dataset S3**, sheet 11). MAGs were partitioned into those containing both identifiable chloroplast-targeted Enolase and PGAM enzymes; one only; and neither. The mean mapped vertical depth (analogous to abundance) was calculated for each MAG from 0.8-2000 μm data at DCM and surface depths, following data from (Delmont, Gaia et al. 2022), and was compared to absolute station latitude by Pearson correlation and two-tailed *t*-test.

### Nucleic acid isolation

For DNA isolation, 150 ml early stationary phase *P. tricornutum* culture, grown under 19°C with 12h light: 12h dark cycling, and shaken at 100 rpm as described above, was centrifuged at 4000 rpm for 10 minutes. The resulting cell pellet was washed in sterile growth medium three times, and incubated for four hours in 5 ml TEN buffer (0.1 M NaCl, 0.01 M Tris pH8, 0.001 M EDTA) supplemented with 2% volume: volume SDS, and 1U μl^-1^ proteinase K (Fisher Scientific). Lysed cell fractions were used for phenol: chloroform precipitation of cellular DNA, as previously described (Nash, Barbrook et al. 2007), prior to dissolution in 50 μl nuclease-free water, and quantification with a nanodrop photospectrometer.

For RNA isolations, 10^5^ stationary phase *P. tricornutum* cells, as calculated with cell densities counted from a Malassez haemocytometer were inoculated in a 250 ml conical Erlenmeyer containing 80 ml ESAW without antibiotics. Cell cultures were harvested in mid-exponential phase, at counted densities of between 1 and 2 x 10^6^ cells ml^-1^. 19C CL cultures were typically harvested eight days post-inoculation, 19C LD cultures nine days post-inoculation, and 8C CL cultures seventeen days post-inoculation, in agreement with growth curve dynamics. Cells were harvested at the mid-point of the light-induction phase of the LD growth condition (15:00 CET), per previous gene expression studies in *P. tricornutum* (Cruz de Carvalho, Sun et al. 2016).

RNA was isolated from 10^8^ cells from each culture, pelleted and washed as before, and snap-frozen in liquid nitrogen. Frozen cell suspensions were re-equilibrated with 1 ml Trizol reagent (Invivogen) and 200 μl chloroform (Honeywell), prior to phenol: chloroform extraction. An additional separation step was performed in 500 μl pure chloroform to remove any residual phenol traces from the aqueous phase, and purified nucleic acids were precipitated overnight in 500 μl isopropanol at -20°C. RNA was collected by centrifugation at 10,000 rpm for 30 minutes, washed with 900 μl 100% ethanol, and resupended in 50 μl RNAse-free water (Qiagen).

2 μg RNA, as quantified with a nanodrop photospectrometer, was subsequently treated with 2U RNAse-free DNAse (Promega) for 30 minutes at 37 °C, and the reaction was stopped with 1 μl 0.5 M EDTA. The digested RNA sample was reprecipitated in isopropanol for one hour at -20 °C, washed in ethanol, and resuspended in 20 μl RNAse-free water. Purified RNA sample concentrations were quantified with a nanodrop spectrometer, and a 3 μl aliquot was migrated on an RNAse-free 1% agarose gel stained with 0.2 μg ml^-1^ ethidium bromide to confirm RNA integrity prior to all downstream applications.

### GFP localization

Full length mRNA sequences of cpEnolase, cpPGAM1A and cpPGAM2 were amplified from *P. tricornutum* RNA libraries grown under 19 °C, light: dark cycling and replete nutrient conditions as described above, by reverse transcription with RT Maxima First Strand synthesis kit (Thermo Fisher) from 200 ng template RNA, following the manufacturer’s instructions; and gene-specific primers as shown in **Supplemental Dataset S2**, sheet 4. PCRs were performed using Pfu high-fidelity DNA polymerase, in 50 μl total reaction volume, including 1 μl cDNA template and 2 μl each specific primer, following the manufacturer’s protocol. Typical PCR conditions were: 10 minutes at 95 °C; followed by 35 cycles of 45 seconds at 95 °C, 45 seconds at 55 °C, and 2 minutes at 72 °C; followed by a terminal elongation phase of 5 minutes at 72 °C. Amplified products were migrated on a 1% agarose gel stained with ethidium bromide, cut out, and purified using a MinElute PCR purification kit (Qiagen).

Purified products were cloned into linearised versions of pPhat vectors containing eGFP and a zeocin resistance gene (SHBLE). These products were amplified using an analogous PCR protocol as above, with 1 ng purified plasmid DNA, and outward-directed PCR primers extending from the start of the fluorescence protein gene sequence to the end of the FcpA promoter region (**Supplemental Dataset S2**, sheet 4); cut, purified, and treated with 1U FastDigest DpnI (Thermo Fisher) to remove any residual plasmid DNA. The gene-specific primers for each cpEnolase and cpPGAM construct were modified with 15 5’ nucleotides complementary to the terminal regions of the FcpA and GFP sequences, allowing cloning of complete vector sequences using a HiFi DNA assembly kit (NEB), following the manufacturer’s instructions.

Assembled vectors were transformed into chemically competent Top10 *E. coli*, and positive clones (as identified by Sanger sequencing of positive colony PCR products) were used to generate purified plasmid DNA with a Plasmid Midi Kit (Qiagen).

Subcellular localization constructs were transformed into wild type *P. tricornutum* Pt186 by biolistic transformation, as previously described (Falciatore, Casotti et al. 1999). 5 x 10^7^ mid-exponential phase cells were plated on a ½ ESAW-1% agarose plate, and left to recover for two days, prior to bombardment with 10 mg 1 μm tungsten beads treated with 5 μg purified plasmid DNA in a Helios gene gun (BioRad) at 1,550 PSI. Cells were left to recover for two days, prior to replating on ½ ESAW-1% agarose plates supplemented with 100 μg ml^-1^ ampicillin, 100 μg ml^-1^ streptomycin, 30 μg ml^-1^ chloramphenicol and 100 μg ml^-1^ zeocin. Plates post-bombardment and for the first two days post-selection were maintained in a low light environment (10 μE m^-2^ s^-1^) prior to return to standard growth conditions.

Positive transformant colonies, as verified by Western Blot with a mouse anti-GFP antibody (Thermo Fisher), were visualised using a SP8 inverted spinning disc confocal microscopy (Leica) under 400 x magnification, with excitation wavelength 485 nm and emission wavelength filters 500-550 nm. GFP-negative colonies were used to confirm detection specificity, and empty-vector GFP (with cytoplasmic localizations) were used as fluorescence positive controls. A minimum of 12 GFP expressing clones were visualised for each construct with consistent localization.

### CRISPR mutagenesis

CRISPR target sequences for cpEnolase and cpPGAM1A genes were identified using PhytoCRISP-Ex (Rastogi, Murik et al. 2016), privileging positions in the N-terminal region of the conserved domain to minimize the probability of enzyme functionality in knockout lines, and uniqueness across the entire *P. tricornutum* genome within the final 11 bp of the target sequence to minimize off-target effects. Primers were designed for each target sequence, and introduced into a pu6:SG1 CRISPR target sequence plasmid by PCR, as previously described (Nymark, Sharma et al. 2016). 2 μg insertion-positive pu6:SG1 plasmids, as confirmed by Sanger sequencing were co-introduced into wild type *P. tricornutum* Pt186 cells by bombardment along with 2 μg HA-tagged Cas9 and pPhat vectors, as described above. Mutant colonies were genotyped using a DNA lysis buffer containing 0.14 M NaCl, 5 mM KCl, 10 mM Tris-HCl pH 7.5, 1% v/v NP40 to generate crude DNA extracts, followed by PCR amplification across the CRISPR target sequences with DreamTaq PCR reagent (Promega) and Sanger sequencing (Eurofins genomics). Mixed mutant: wild-type colonies were segregated via repeated dilution on ESAW: zeocin plates until only homozygous mutant genotypes were detected (Nymark, Sharma et al. 2016, McCarthy, Smith et al. 2017). Empty vector control lines were generated using the same protocol, except with only HA-Cas9 and pPhat plasmids, cotransformed without a CRISPR target sequence.

Tabulated cleaned knockout mutants, their associated genotypes and the expression levels of mutated gene copies are shown in **Supplemental Dataset S4**, sheets 1-2. Mutant colony genotypes were periodically confirmed (approx. once per month) by PCR and Sanger sequencing throughout the duration of all subsequent experiments, and the CRISPR-induced gene modifications were found to remain stable. *P. tricornutum* Enolase proteins were determined by Western blot to be crossreactive to an anti-*Arabidopsis thaliana* Enolase-2 antibody (Agrisera), and thus knockout line protein expression was confirmed by qRT-PCR, as described below.

### Complementation of knockout lines

Knockout lines were complemented with pPhat:GFP vectors carrying overexpressing copies (under an FcpA promoter) of cpEnolase and cpPGAM1A synthetic constructs, with all CRISPR target sequences replaced with silent mutations (Eurofins). Genes were fused to C-terminal GFP, allowing the verification of protein expression and localization. Vectors were identical to those previously used for localization, but with a blasticidin S-deaminase gene in lieu of SHBLE (Buck, Río Bártulos et al. 2018) introduced by NEB Hi-Fi kit as before. Complementation constructs were transformed via bombardment, and cotransformed colonies were selected on ½ ESAW-1% agarose plates supplemented with 100 μg ml^-1^ ampicillin, 100 μg ml^-1^ streptomycin, 30 μg ml^-1^ chloramphenicol, 100 μg ml^-1^ zeocin, 4 μg ml^-1^ blasticidin.

For each complementation, three cpEnolase and cpPGAM1A knockout lines (including at least one for each CRISPR target sequence) were complemented both with the conjugate construct, and an empty blasticidin resistance vector as a placebo; and two empty vector lines were further complemented with both cpEnolase and cpPGAM1A overexpressor constructs, plus an empty blasticidin resistance vector, to exclude possible effects from ectopic overexpression of each gene on cell physiology. A total of 47 colonies, with a minimum of 6 colonies for each knockout: complementation combination, including lines complemented from at least two distinct primary knockout mutant genotypes, were selected for subsequent study (**Supplemental Dataset S4**, sheet 7). The retention of the primary knockout mutant genotype in each complemented line was verified by colony PCR and sequencing as above, and the overexpression and correct localization of the complementing protein sequence (i.e., to the chloroplast for cpEnolase:GFP and cpPGAM1:GFP, or the cytoplasm for ectopic GFP) was verified by western blot with an anti-GFP antibody (Thermo Fisher) (Erdene-Ochir, Shin et al. 2019) and confocal microscopy.

### Growth rate measurements

A starting density of 10^4^ ml^-1^ stationary phase *P. tricornutum* cells of a given culture line, as verified with a Malassez haemocytometer, were inoculated into a 15 ml volume antibiotic-free ESAW within a sterile, ventilated cap plastic culture flask (Celltreat) and grown under LD, CL, or 8C culture conditions as described. Cell densities were recorded: every day from one day post-inoculation (CL); every day from two days post-inoculation (LD); or every two days from five days post-inoculation (8C) at the mid-point of the LD light induction phase using a counting CyFlow Cube 8 cytometer (ParTec).

Typically, 15 μl cell culture, passed at 0.5 μl s^-1^, were used for each measurement, with three technical replicates performed for each culture of which the first (enriched in non-cellular particles) was excluded from downstream calculations. Cytometer particle counts were correlated to actual cell densities using a calibration curve realised from haemocytometer counted densities of wild-type cell culture, and cultures with observed densities > 2 x 10^6^ cells ml^-1^ were diluted ten-fold in blank growth media to avoid saturation of the cytometer.

Cell densities were measured until cell lines were confirmed to have exited log phase (i.e., reached a stationary phase plateau). Primary knockout mutant growth curves were repeated a minimum of six times (three biological replicates per-inoculation, with two independent repetitions) for each mutant line. Growth curves were tested for seven cpEnolase knockout, five cpPGAM1A knockout and four empty vector control lines, providing a minimum of 24 measurements (i.e., four distinct mutant lines) per each genotype studied (cpEnolase knockout, cpPGAM1A knockout and empty vector control lines).

To avoid artifacts based on the proximity of the seed cell culture to exponential phase at the time of inoculation (which may impact on lag phase length) or the relative diameter of each cell in culture (which may impact on carrying capacity), cell growth rates were measured exclusively from the log-phase relative division rate. This was calculated via considering Δlog_2_ (cell density) / Δlog_2_ (time) for a time-period corresponding to 5 x 10^4^ to 4 x 10^6^ cells/ ml, covering in most cases six successive measurements of each individual growth curve. To confirm that the cells were measured in exponential phase and were neither influenced by particle contamination of the cytometer or cell exhaustion of the growth medium, the linear correlation was calculated between the log value, with most calculated correlations (129/ 132) showing linearity (*r*> 0.95). Three exemplar growth curve outputs are provided in **Supplemental Dataset S4**, sheets 3-5; and an overview of relative growth rates expressed as a function of mean empty vector control growth rates are provided in **Supplemental Dataset S4**, sheet 6.

Complementation growth curves were repeated with at least two independent repetitions for each cell line, with five timepoints taken to project growth rates, and therefore a minimum of sixty independent measurements for each mutant: complementation genotype under each growth condition, with the average of the two fastest growth rates of each culture calculated as estimates for the growth rate. A heatmap of all estimated complementation line growth rates is provided in **Supplemental Dataset S4**, sheet 7.

### Photophysiology

Cultures for photophysiological analysis were grown in 10ml ventilated plastic flasks, without shaking, under 19C CL and 19C LD as described above. Cultures were grown from a starting inoculum of 10^5^ cells ml^-1^ as measured with a Malassez haemocytometer. Cell cultures that had reached a measured density of 10^6^ cells ml^-1^ were then refreshed into fresh media at the initial starting concentration of 10^5^ cells ml^-1^ to allow a prolonged adaptation to each photophysiological condition under a continuous exponential phase. Cells from refreshed culture lines were harvested in exponential phase (between 1 and 3 × 10^6^ cells ml^-1^, and good physiology was verified by Fv/Fm measurements > 0.6 across all measured lines (**Supplemental Dataset S4**, sheet 8).

Steady-state light curves (SSLC) were conducted with a fluorescence CCD camera recorder (SpeedZen, jBeamBio, France) in a selected set of control lines (*n*=2), cpPGAM (*n*=3) and cpEnolase knockouts (*n*=6), as well in complemented cpEnolase (*n*=2) and cpPGAM1A (*n*=3) knockout lines in which we observed a suppression of the knockout growth defect compared to complemented control lines. Measurements were repeated a minimum of two and in most cases four times per line and treatment condition, with a minimum of six unique measurements performed for each genotype and treatment. Curves were measured on cell cultures concentrated to between 2 and 5 × 10^7^ cells ml^-1^. Samples were exposed to an initial 5 min illumination of 35 µmol photons m^-2^ s^-1^ green actinic light (532 nm), followed by a 6 steps of 3 min each of increasing intensity to 750 µmol photons m^-2^ s^-1^.

Minimum (*F_0_*) and maximum (*F_M_*) fluorescence were measured in dark-adapted (at least 1 min) samples, before and at the end of a 250 ms saturating (multiple turnover) pulse of light (532 nm, 5000 µmol photons m^-2^ s^-1^) and the maximum quantum yield of PSII in the dark was calculated as *F_V_*/*F_M_*= (*F_M_* -*F_0_*)/*F_M_.* Every minute of light exposure, steady-state fluorescence (*F_S_*) and maximum fluorescence under Light (*F_M_^’^*) were measured. PSII quantum yield (φPSII) and nonphotochemical quenching (NPQ) were calculated on the last time point of each light step as φPSII = (*F_M_*’-*F_s_*)/*F_M_*’ and NPQ = *F_M_*/*F_M_’*-1, and rETR at PSII as rETR = φPSII.*E*.

The whole rETR vs *E* curve was fitted as rETR = rETR_M_.(1-exp(-α.*E*/rETR_M_)) where rETR_M_ is the maximum rETR and α is the light-limited slope of rETR vs E (Jassby and Platt 1976). Only rETR values from 0 to 450 µmol photons m^-2^ were used for the fit because values from 600 and 750 µmol photons m^-2^ were too noisy. The light saturation parameter *E_K_* was calculated as rETR_M_/α and the fitted values of the parameters were used to estimate φPSII under the growth light intensity of 50 µmol photons m^-2^ s^-1^ as φPSII_50µE_= (rETR_M_.(1-exp(-α.50/rETR_M_)))/ 50. The NPQ vs E curve was fitted as NPQ = NPQ_M_× *E^n^*/ (E_50_NPQ*^n^*+*E^n^*), where NPQ_M_ is the maximal NPQ, E_50_NPQ is the half-saturation intensity for NPQ and *n* is the sigmoidicity coefficient (Serôdio and Lavaud 2011).

The PSII functional absorption cross-section, σPSII, was calculated from the fluorescence induction upon a single turnover flash of blue light (100 µs, 455 nm, 60 nm bandwidth) on non-concentrated cell culture. The induction curve was measured on 20 min dark-acclimated samples before centrifugation (average of 2-4 independent replicates) with a Fluorescence Induction and Relaxation (miniFIRe) fluorometer (Gorbunov, Shirsin et al. 2020), which also measures single turnover *F_V_*/*F_M_* and PSII connectivity. Parameters measured with the miniFIRe fluorometer (as defined below) were also quantified for cultures grown under 8C CL, as the measurements were sufficiently rapid to allow the culture to be maintained at growth temperatures (Gorbunov, Shirsin et al. 2020). Measured photophysiological values are tabulated in **Supplemental Dataset S4**, sheet 8. Violin plots of photophysiological parameters were generated with BoxPlotR (Spitzer, Wildenhain et al. 2014).

### Gene expression analysis

Libraries were prepared from 200 ng DNAse-treated RNA for each mutant line and treatment condition, with at least three replicates per sample. Sequencing was performed by Fasteris (Plan-les-Ouates, Switzerland). After initial quality control checks, stranded Illumina mRNA libraries were prepared with a Novaseq V1.5 kit and sequenced with an SP-flow cell with 2x 100 bp over 200 cycles, yielding circa 130-160 Gb sequence data per sample with ≥85% of bases higher than Q30.

FastQ files were mapped using Nextflow’s RNA sequencing assembly pipeline https://nf-co.re/rnaseq/usage, to gff3 annotations of the *P. tricornutum* version 3 genome (Rastogi, Maheswari et al. 2018, Lataretu and Hölzer 2020). Total mapped read counts were then compared between all biological and technical replicates for (i) each mutant line sequenced, (ii) each genotype (cpEnolase knockout, cpPGAM1A knockout, control), and (iii) each treatment condition performed (LD, CL, 8C) by principal component analysis (PCA) using the R package factoextra, with highly variant libraries removed (Kassambara and Mundt 2017). The final dataset included 63 RNAseq libraries, including five cpEnolase and five cpPGAM1A knockout lines and four empty vector controls, and a minimum of four RNA libraries prepared from at least two genetically distinct mutant constructs for each genotype (cpEnolase, cpPGAM1A and control) considered (**Supplemental Dataset S5**, sheets 1-2)., Differentially expressed genes (DEGs) were then calculated between each genotype for each treatment condition using DESeq2 with cutoff fold-change 2 and P-value 0.05 (Liu, Wang et al. 2021) (**Supplemental Dataset S5**, sheets 2-3).

The mean transcript abundances of DEGs in knockout compared to control lines were first assessed in RNAseq data of N and P-limited *P. tricornutum* cell lines under two and nine time-points respectively (**Supplemental Dataset S5**, sheet 4) (Cruz de Carvalho, Sun et al. 2016, McCarthy, Smith et al. 2017). No significant differences were found between DEGs and other genes in the *P. tricornutum* genome (one-way ANOVA, P > 0.05; **Supplemental Dataset S5**, sheet 5), confirming that the RNAseq samples were not generated from N- or P-limited cultures. Next, functional enrichments in DEGs from previously tabulated values for the entire *P. tricornutum* genome (**Supplemental Dataset S5**, sheets 6-10) (Rastogi, Maheswari et al. 2018, Ait-Mohamed, Novák Vanclová et al. 2020). Functional enrichments were tested by two-tailed chi-square (P < 0.05) of a differentially expressed gene occurring in either one (cpEnolase v control; cpPGAM1A v control) knockout-versus-control line tests, or in both tests realised under each physiological condition. Finally, the distribution of DEGs across *P. tricornutum* core plastid and mitochondrial metabolism pathways were mapped onto a previously defined proteomic model of each organelle (Ait-Mohamed, Novák Vanclová et al. 2020); with the strongest DEG enrichment taken in the case of enzymes with multiple homologues (**Supplemental Dataset S5**, sheet 11).

Quantitative RT-PCR (qRT-PCR) validations were performed using cDNA synthesised from 5 ng dNase-treated RNA (per qRT-PCR reaction) and a RT Maxima First Strand synthesis kit (Thermo Fisher), following the manufacturer’s instruction; using a 384-well Lightcycler (Roche) and Takyon™ No ROX SYBR 2X MasterMix (Eurogentec), following the manufacturers’ protocols. Typical amplification conditions were: 10 minutes at 95°C, followed by 40 cycles of 30 seconds at 95°C, 30 seconds at 55°C, and 30 seconds at 72°C. Primer pairs for qRT-PCR amplifications were designed using NCBI Primer-BLAST (Ye, Coulouris et al. 2012), privileging unique amplification targets within the genomic sequence, an amplicon size of 100 to 150 base pairs, primer positions at different regions of the gene studied, and a 3’ terminal G or C on each primer. Primer efficiencies were tested by qPCR with serial dilutions of *P. tricornutum* gDNA, with only primer pairs that yielded a Cp increment of between 0.9 and 1.1 per half dilution of DNA retained for qRT-PCR analysis. qRT-PCRs were at least three times for each amplicon: sample pair. RT-equivalents were performed to subtract residual genomic DNA from each Cp value obtained, and two housekeeping genes (Ribosomal protein S1, RPS; and TATA binding protein, TBP) previously shown to have conditionally invariant expression patterns in *P. tricornutum* were used as quantification references (Sachse, Sturm et al. 2013). Tabulated qRT-PCR outputs are shown in **Supplemental Dataset S5**, sheet 13; and sample information and reaction conditions per MIQE guidelines (Bustin, Benes et al. 2009) are tabulated in **Supplemental Dataset S5**, sheet 14.

### Metabolite analysis

Cell pellets were taken from exponential-phase *P. tricornutum* culture (counted density 1-2 x 10^6^ cells ml^-1^, 1.5 x 10^8^ cells per sample) for metabolite and lipid analysis. Cell pellets were collected without washing to minimise impacts on metabolite turnover, then transferred to a pre-weighed, double-pierced and double-autoclaved 1.5 ml Eppendorf tube for lyophilization. Cell pellet masses were recorded, and samples were immediately snap-frozen in liquid nitrogen and stored at -80 °C for subsequent analysis.

Metabolite profiling was carried out by gas chromatography–mass spectrometry (ChromaTOF software, Pegasus driver 1.61; LECO) as described previously (Lisec, Schauer et al. 2006). The chromatograms and mass spectra were evaluated using TagFinder software (Luedemann, von Malotky et al. 2012). Metabolite identification was manually checked by the mass spectral and retention index collection of the Golm Metabolome Database (Kopka, Schauer et al. 2005). Peak heights of the mass fragments were normalized successively on the basis of the fresh weight of the sample, the added amount of an internal standard (ribitol) and values obtained for loading column controls obtained from the same experiment.

### Glycerolipid analysis

Glycerolipids were extracted by suspending cell pellets in 4 mL of boiling ethanol for 5 minutes to prevent lipid degradation. Lipids were extracted by addition of 2 mL methanol and 8 mL chloroform at room temperature (Folch, Lees et al. 1957). The mixture was then saturated with argon and stirred for 1 hour at room temperature.

After filtration through glass wool, cell remains were rinsed with 3 mL chloroform/methanol 2:1, v/v and 5 mL of NaCl 1% was added to the filtrate to initiate biphase formation. The chloroform phase was dried under argon and stored at -20°C. The lipid extract was resuspended in pure chloroform when needed.

Total glycerolipids were quantified from their fatty acids: in an aliquot fraction, 5 µg of 15:0 was added and the fatty acids present were converted to methyl esters (FAME) by a 1-hour incubation in 3 mL 2.5% H_2_SO_4_ in pure methanol at 100 °C (Jouhet, Maréchal et al. 2003). The reaction was stopped by addition of 3 mL water and 3 mL hexane. The hexane phase was analyzed by a gas chromatography-flame ionization detector (GC-FID) (Perkin Elmer) on a BPX70 (SGE) column. FAMEs were identified by comparison of their retention times with those of standards (Sigma) and quantified using 15:0 for calibration.

Glycerolipids were further analyzed by high pressure liquid chromatography-tandem mass spectrometry (HPLC-MS/MS), based on a previously described procedure (Rainteau, Humbert et al. 2012). The lipid extracts corresponding to 25 nmol of total fatty acids were dissolved in 100 µL of chloroform/methanol [2/1, (v/v)] containing 125 pmol of each internal standard. Internal standards used were phosphatidylethanolamine (PE) 18:0-18:0 and diacylglycerol (DAG) 18:0-22:6 from Avanti Polar Lipid, and sulfoquinovosyldiacylglycerol (SQDG) 16:0-18:0 extracted from spinach thylakoids (Demé, Cataye et al. 2014) and hydrogenated (Buseman, Tamura et al. 2006). Lipid classes were separated using an Agilent 1200 HPLC system using a 150 mm × 3 mm (length × internal diameter) 5 µm diol column (Macherey-Nagel), at 40 °C. The mobile phases consisted of hexane/ isopropanol/ water/ 1 M ammonium acetate, pH 5.3 [625/350/24/1, (v/v/v/v)] (A) and isopropanol/ water/ 1 M ammonium acetate, pH 5.3 [850/149/1, (v/v/v)] (B). The injection volume was 20 µL. After 5 min, the percentage of B was increased linearly from 0% to 100% in 30 min and kept at 100% for 15 min. This elution sequence was followed by a return to 100% A in 5 min and an equilibration for 20 min with 100% A before the next injection, leading to a total runtime of 70 min. The flow rate of the mobile phase was 200 µL min^-1^. The distinct glycerophospholipid classes were eluted successively as a function of the polar head group. Mass spectrometric analysis was performed on a 6460 triple quadrupole mass spectrometer (Agilent) equipped with a Jet stream electrospray ion source under following settings: drying gas heater at 260 °C, drying gas flow at 13 L·min^-1^, sheath gas heater at 300 °C, sheath gas flow at 11 L·min^-1^, nebulizer pressure at 25 psi, capillary voltage at ± 5000 V and nozzle voltage at ± 1,000 V. Nitrogen was used as the collision gas. The quadrupoles Q1 and Q3 were operated at widest and unit resolution, respectively.

Phosphatidylcholine (PC) and diacylglyceryl hydroxymethyltrimethyl-*β*-alanine (DGTA) analyses were carried out in positive ion modes by scanning for precursors of m/z 184 and 236 respectively at a collision energy (CE) of 34 and 52 eV. SQDG analysis was carried out in negative ion mode by scanning for precursors of m/z -225 at a CE of -56eV. PE, phosphatidylinositol (PI), phosphatidylglycerol (PG), monogalactosyldiacylglycerol (MGDG) and digalactosyldiacylglycerol (DGDG) measurements were performed in positive ion modes by scanning for neutral losses of 141 Da, 277 Da, 189 Da, 179 Da and 341 Da at cEs of 20 eV, 12 eV, 16 eV, 8 eV and 8 eV, respectively. DAG and triacylglycerol (TAG) species were identified and quantified by multiple reaction monitoring (MRM) as singly charged ions [M+NH4]^+^ at a CE of 16 and 22 eV respectively. Quantification was done for each lipid species by MRM with 50 ms dwell time with the various transitions previously recorded (Abida, Dolch et al. 2015). Mass spectra were processed using the MassHunter Workstation software (Agilent) for lipid identification and quantification. Lipid amounts (pmol) were corrected for response differences between internal standards and endogenous lipids as described previously (Jouhet, Lupette et al. 2017).

Normalised metabolite and lipid abundances were screened by PCA, as per the RNAseq analysis above, and outliers and biologically non-representative samples were removed. The final datasets consist of 139 libraries (metabolite GC-MS), 55 libraries (lipid GC-MS) and 49 libraries (lipid LC-MS), with a minimum of three libraries prepared from at least two genetically distinct mutant constructs for each genotype considered (**Supplemental Dataset S6**, sheet 1). Violin plots of differentially accumulated lipids were generated with BoxPlotR (Spitzer, Wildenhain et al. 2014).

### Expressed enzyme reaction kinetics

Measurements of cpEnolase and cpPGAM1A reaction rates were performed following a previously defined protocol (Zhang, Sampathkumar et al. 2020) (**Fig. S20**). First, codon-optimised constructs for *E. coli* expression were synthesized (Eurofins) using full-length cpEnolase and cpPGAM1A mRNA sequences as template. Constructs were cloned into a Gateway pDest-CTD-His vector via a pDONR intermediate and BP /LR clonase (all reagents Thermo Fisher) following the manufacturer’s instructions (Hartley, Temple et al. 2000). To enable optimal expression in *E. coli*, multiple N-terminal length variants were synthesized from each gene, with those corresponding to the full gene length minus the predicted N-terminal signal peptide domain as inferred with SignalP (Almagro Armenteros, Tsirigos et al. 2019). Complete constructs and primers tested are provided in **Supplemental Dataset S6**, sheet 7.

cpEnolase and cpPGAM1A –CTDHis vectors were cloned into Rosetta (DE3) strain *E. coli* (Novagen) and coselected on ampicillin (100 µg /ml) and chloramphenicol (34 µg /ml). Proteins were induced in overnight cultures at 28°C, purified on a His-Trap column (GE Healthcare) following the manufacturers’ instructions, and eluted in a buffer consisting of 125 mM NaCl, 250 mM Imidiazol (Sigma) and protease inhibitors. Eluted proteins were desalted using a Q10/ PD10 column (GE Healthcare) and quantified using a Bradford. Protein integrity and quantity were assessed routinely throughout the purification using SDS-PAGE.

Reaction rates were measured on purified 100 µg cpPGAM1A and 100 µg cpEnolase, as quantified with a nanodrop spectrometer. Rates were measured separately for glycolytic and gluconeogenic activity. Briefly, to measure glycolytic reaction rates, both enzymes were combined with 10 units pyruvate kinase and 10 units lactate dehydrogenase (both Sigma-Aldrich) at 25°C, alongside varying concentrations 9 mM D(-)3-Phosphoglyceric Acid, 25 mM Adenosine ’’-Diphosphate, and 25 mM reduced ß-Nicotinamide Adenine Dinucleotide (NADH). Enzymatic activity was measured by considering 340 nm colorimetry as a proxy for NADH consumption following a previously defined protocol (Sigma protocols EC 5.4.2.1) (Sutherland, Posternak et al. 1949). To measure gluconeogenic reaction rates, a similar reaction was performed with both enzymes combined with 10 units phosphoglycerate kinase and 10 units glyceraldehyde-3-phosphate dehydrogenase (both Sigma-Aldrich), alongside 9 mM phospho-enol-pyruvate, 25 mM Adenosine 5’-Diphosphate, and 25 mM reduced ß-Nicotinamide Adenine Dinucleotide (NADH).

Enzymatic activity was similarly measured by 340 nm colorimetry. A schematic of the measured reactions is provided in **Fig. S20**. Complete measured reaction rates over all technical replicates are provided in **Supplemental Dataset S6**, sheet 8.

### Accession Numbers

RNAseq data associated with this project is deposited with NCBI BioProject with project number PRJNA788211.

### Materials Distribution Statement

The author(s) responsible for distribution of materials integral to the findings presented in this article in accordance with the policy described in the Instructions for Authors (https://academic.oup.com/plcell/pages/General-Instructions) are: Richard G. Dorrell and Chris Bowler.

## List of Supporting Files

**Supplemental Dataset S1. Phylogenetic diversity of Enolase and PGAM sequences from across the tree of life.**

**Supplemental Dataset S2. Transcriptional and localization patterns of cpPGAM and cpEnolase genes in *Phaeodactylum tricornutum*.**

**Supplemental Dataset S3. *Tara* Oceans analysis of diatom plastid glycolysis.**

**Supplemental Dataset S4. Genotyping, growth dynamics and photophysiology in *P. tricornutum* plastid glycolysis mutant lines.**

**Supplemental Dataset S5. Differentially and conditionally expressed genes in *P. tricornutum* plastid glycolysis mutants.**

**Supplemental Dataset S6. Lipid and metabolite profiles of *P. tricornutum* plastid glycolysis mutant lines revealed by GC- and LC-mass spectrometry, and measured reaction kinetics of expressed enzymes.**

All remaining supporting data not provided directly in paper supporting tables are provided in the linked Open Science Foundation Supporting database https://osf.io/89vm3/ (Dorrell, Novak Vanclova et al. 2022). Project contents are ordered hierarchically by theme, with an overview of all contents provided on the site wiki page. A dedicated README file in each project folder explains the data presented and provides detailed methodology for each analysis.

## Author Contributions

RGD designed the research, with critical input from YZ, YZ, DC, BB, ARF, JJ, EM and CB. RGD, YZ, NG, TN, DC, MP, and VG performed the research. RGD, YZ, YL, DC and MP analysed the data. SA provided new analytical tools for cell growth measurements, and JJPK and NZ provided new computational tools for meta-genomic and RNA-seq analysis. RGD wrote the paper, with critical input from YL, TN, MP, SA, BB, JJPK, JJ, EM and CB. All coauthors read and approved the manuscript draft prior to final submission.

## Acknowledgments

RGD acknowledges a CNRS Momentum Fellowship, awarded from 2019-2021, an ANR JCJC Grant (« PanArctica » ANR-21-CE02-0014-01) awarded 2021-2022 and an ERC Starting Grant Grant (ChloroMosaic, 101039760), awarded from 2023-2027. CB acknowledges support from FFEM - French Facility for Global Environment, French Government ‘Investissements d’Avenir’ programs OCEANOMICS (ANR-11-BTBR-0008), FRANCE GENOMIQUE (ANR-10-INBS-09-08), MEMO LIFE (ANR-10-LABX-54), and PSL Research University (ANR-11-IDEX-0001-02), the European Research Council (ERC) under the European Union’s Horizon 2020 research and innovation program (Diatomic; grant agreement No. 835067), and from the ANR ‘BrownCut’ project (ANR-19-CE20-0020). CB, EM and JJ were supported by ANR ‘DIM’ (ANR-21-CE02-0021) and PEPR AlgAdvance (22-PEBB-0002). The lipid analyses were performed at the LIPANG (Lipid analysis in Grenoble) platform hosted by the LPCV (UMR 5168 CNRS-CEA-INRAE-UGA) and supported by the Rhône-Alpes Region, the European Regional Development Fund (ERDF), Institut Carnot 3BCAR and Labex GRAL (10-LABX-0049), financed within the University Grenoble Alpes graduate school (Ecoles Universitaires de Recherche) CBH-EUR-GS (ANR-17-EURE-0003). YZ and ARF acknowledge funding from the European Union’s Horizon 2020 research and innovation program, project PlantaSYST (SGA-CSA No. 739582 under FPA No. 66462; and the BG05M2OP001-1.003-001-C01 project, financed by the European Regional Development Fund through the Bulgarian ‘Science and Education for Smart Growth’ Operational Programme. D.C and B.B acknowledge the support of the European Research Council (ERC) under the European Union’s Horizon 2020 research and innovation program (PhotoPHYTOMIX project, grant agreement No. 715579). The authors acknowledge Giselle McCallum, Elena Kazamia and Xia Gao (IBENS) for assistance with the optimization of cell cytometry protocols; Amandine Baylet (Lycée ENCPB-Pierre Gilles de Gennes), Quentin Caris (Lycée de la Vallée de Chevreuse) and Yonna Lauruol (Ecole Sup’BioTech) for assistance with growth measurements, Frédy Barneche and Clara Richet-Bourbousse (IBENS) for the use of culture facilities; Nathalie Joli (IBENS) for aid with RNAseq library production; Catherine Cantrel and Priscillia Pierre-Elies (IBENS), Pauline Clémente (Lycée ENCPB-Pierre Gilles de Gennes) and Shun-Min Yang (University of South Bohemia) for the preparation of media substrates and aid with biolistic transformations; Ansgar Gruber (University of South Bohemia) for advice concerning the experimental localisation of chloroplast glycolysis proteins; Max Gorbunov (Rutgers University, NJ, USA) for the provision of the miniFIRE fluorometer used for photophysiological assays; and Mattia Storti (CEA Grenoble) for cell outlines used in the production of **Fig. 1A**. This paper is contribution XXX of *Tara* Oceans.

---

**Fig. S1. Distribution of lower plastid glycolysis-glucneogenesis across photosynthetic eukaryotes. A:** Occurrence of plastid-targeted enolase and PGAM enzymes across 1,673 plant and algal genomes and transcriptomes, inferred using reciprocal BLAST best hit of *P. tricornutum* query enzymes as per Fig. 1B and 1C, PFAM domain annotations, and in silico targeting predictions with TargetP and PredAlgo (primary chloroplast bearing lineages) and HECTAR and ASAFind (secondary lineages). **B:** scatterplots of collection site latitude for (**i)** diatoms, (ii) other stramenopiles, **(iii)** cryptomonads and haptophytes and **(iv)** green algae with detectable enolase and PGAM enzymes, divided by presence of inferred plastid-targeted isoforms. Notably, diatoms lacking both plastid-targeted glycolysis enzymes do not occur outside of low and intermediate latitudes (50°N in the northern and 60°S in the southern hemisphere) compared to other groups, which show no significant association between plastid glycolysis and latitude. The data in this figure were subselected for the phylogenies shown in Fig. 1 and support the latitudinal correlations revealed by Tara analysis of Fig. 3.

**Fig. S2. Consensus topology of a 380 taxa x 413 aa alignment of Enolase sequences.** Sequences represent a sample of all organelle-targeted isoforms from cryptomonads, haptophytes and stramenopiles and representatives from a densely-sampled dataset of 151 taxonomic groups across the tree of life (Dorrell et al., 2021). The tree topology shown is the consensus of the best-scoring rAxML trees identified using three substitution matrices: GTR, JTT, and WAG. Branch thickness corresponds to frequency of consensus tree topology recovery in individual trees; branches are coloured by taxonomic affiliation; and tips (cryptomonads, haptophytes and stramenopiles only) by predicted *in silico* localization. Individual clades (considering both taxonomic origins and inferred localization) of organelle-targeted enolase isoform are labelled with coloured brackets. This figure extends on the topology shown in Fig. 1B.

**Fig. S3. Consensus topology of a 220 aa x 560 taxa alignment of PGAM isoform 1 sequences.** Data are shown as per **Fig. S1**, extending on the topology of Fig. 1C.

**Fig. S4. Consensus phylogeny of a 235 aa x 66 taxa alignment of PGAM isoform 2 sequences.** Data are shown as per **Fig. S1,** forming a complement to the PGAM1 topologies shown in Fig. 1C and **S3.**

**Fig. S5. PGAM2 and control confocal microscopy images for *P. tricornutum* plastid glycolysis proteins.** Images complement those shown in Fig. 2A.

**Fig. S6. Relative abundances and transcriptional coordination of plastidial enolase, PGAM1 and PGAM2. A:** relative % accumulation of different plastid- and mitochondrial-associated carbon metabolism proteins (large points) and all known plastid-encoded or inferred plastid-targeted proteins (small points, with power-law trendline) for experimental proteomics of *Phaeodactylum tricornutum* total cell proteomes (horizontal axis) or plastid-enriched protein fractions (vertical axis), following (Huang, Pan et al. 2024). Only proteins detected in at least two replicates of plastid-enriched and total cell fractions are included, and abundances are renormalized based on the total abundances of these proteins. Both cpEnolase (Phatr3_J41515) and cpPGAM1A (Phatr3_J17086) are detected in the plastid-associated proteome fraction with similar frequencies to other plastid carbon metabolism proteins, whereas other predicted plastid-targeted PGAMs (e.g. cpPGAM1B, Phatr3_J51404; cPGAM2, Phatr3_J37201) are not. B: Spearman correlation coefficients of all genes across the version 3 annotation of the *Phaeodactylum tricornutum* genome against cpEnolase and cpPGAM1A; calculated from ranked transcript data (Liu, Storti et al. 2022) from two merged datasets, PhaeoNet (Ait-Mohamed, Novák Vanclová et al. 2020) and DiatomPortal (Ashworth, Turkarslan et al. 2016). Genes are coloured by inferred *in silico* localisation; and other annotated enolase and PGAM enzymes in the *Phaeodactylum* genome are shown as large, labelled points. Notably, while cpEnolase and cpPGAM1A show a strong positive *(r* > 0.8) coregulation to one another, other PGAM isoforms (e.g. cpPGAM1B, cpPGAM2) show less apparent coregulation to either of these two genes. These data support the experimental localisations shown in Fig. 2A, and identify cPGAM1A as the PGAM most likely to work cooperatively with cpEnolase from the homologues shown in Fig. 1C.

**Fig. S7. Relative transcriptional regulation of cpEnolase and cpPGAM under nutrient, light and temperature stress conditions. A-C:** the relative ratio of plastid to mitochondria-targeted gene isoform expression for enolase (vertical axis), PGAM1 (horizontal axis), and PGAM2 (bubble size) in published *Phaeodactylum* RNAseq data subject to nitrate limitation (McCarthy, Smith et al. 2017), phosphate starvation and addition (Cruz de Carvalho, Sun et al. 2016) and Circadian Fe enrichment and limitation (Smith, Gillard et al. 2016). While limited transcriptional responses are identifiable in response to changing N, P or Fe availability, the relative transcription of plastidial to mitochondrial-targeted enolase is substantially greater in RNA sampled in long day (> 12h post-illumination) conditions in **C**. **D**: relative fold changes in plastidial enolase and PGAM1 expression in published microarray data assembled in (Ashworth, Turkarslan et al. 2016), under different illumination conditions. Both Enolase and PGAM1 show substantial downregulation in dark-incubated (48h) and short post-illumination-incubated (< 0.5h) cultures, and show the greatest positive fold expression changes respectively > 16h and > 12h post-illumination. Specific data points showing either very high or low cpEnolase and cpPGAM1A expression are labelled.These data support the qRT-PCR and *Tara* Oceans observations of correlation of diatom cpEnolase and cpPGAM expression in Fig. 2 and Fig. 3.

**Fig. S8. Identification of *Tara* Oceans homologs of diatom plastid-targeted enolase and PGAM enzymes. A:** Consensus rAxML JTT topologies of the phylogenetically verified *Tara* Oceans homologs of diatom plastidial enolase and PGAM enzymes and cultured species sequences, demonstrating reconciliation of retained homologs within monophyletic clades containing exclusively diatom plastidial isoforms amongst cultured species. **B:** *in silico* targeting predictions of all retrieved homologs inferred by BLAST alignment to be probably N-terminally complete, showing a strong enrichment in homologs with predicted plastid-targeting sequences. Sequences shown in this figure are analysed globally in Fig. 3.

**Fig. S9. Relative abundances of *Tara* Oceans diatom plastid glycolysis meta-genes.** Plots show relative abundances of meta-genes that group with **(i, iii)** cpEnolase and **(ii, iv**) cpPGAM1 sequences over individual size fractions of **(i, ii)** surface and **(iii, iv)** DCM meta-transcriptome **(left)** and -genome **(right)** data. These data, which allows us to identify whether specific trends are observed in different kinds of diatom cells, ranging from the nano to micro-metre scale, support global trends observed across from 0.8-2000 μm filtered size fractions and surface layers shown in Fig. 3.

**Fig. S10. Normalised atitudinal regressions of *Tara* cpEnolase and cpPGAM1 sequences.** Scatterplots of *Tara* Ocean expression patterns of sequences assigned phylogenetically to diatom cpEnolase and cpPGAM1A against station latitude. Abundances are shown for 0.8-2000 μm surface and DCM sample meta-transcriptome data, and are normalised relative to **(A**) total diatom metaT abundances at each station and **(B)** the corresponding metaG abundances for diatom cpEnolase and cpPGAM1A. Best-fit (linear) regression lines are provided for each depth. In each case, a significant positive correlation between latitude and relative expression is observed, consistent with global distributions observed in Fig. 3.

**Fig. S11. Total relative abundances of meta-genes phylogenetically reconciled to diatom PGAM2 in 0.8-2000 μm filtered surface samples.** Plots showing (**A**) meta-transcriptome and (**B**) meta-genome data, showing effective congruence between both, in contrast to the high latitudinal abundance specific to meta-transcriptome data for diatom cpEnolase and cpPGAM1 as per Fig. 3.

**Fig. S12. Occurrence and mean coverage depth of *Tara* Oceans MAGs divided by chloroplast glycolysis state.** This figure shows mapped distributions for *Tara* meta-genome assembled genomes (MAGs (Delmont, Gaia et al. 2022)) pertaining to (**A**) diatoms, (**B**) chlorophytes, and (**C**) other algal groups (haptophytes, pelagophytes, dictyochophytes, chrysophytes, bolidophytes, cryptomonads) containing members with inferred lower half plastidial glycolysis. MAGs are divided by the inferred occurrence of which chloroplast-targeted enolase and PGAM enzymes could be inferred using combined RbH to *Phaeodactylum* queries, PFAM annotation, and in silico targeting prediction (TargetP, WolfPSort, PredAlgo, HECTAR, ASAFind): (i) detection of plastid-targeted homologues of both enzymes; (**ii**) detection of cpEnolase or cpPGAM only; (**iii**) detection of neither. Bubble sizes correspond to the mean vertical coverage of meta-gene reads recruited to each MAG as a proxy of abundance. In each case the linear correlation and P-value (two-tailed *t*-test) of correlation between mean vertical mapped depth and absolute latitude is provided. Notably there is a positive correlation between the retrieval of either plastid glycolysis protein and greater mapped read depth at high latitudes in diatoms, and also the retrieval of both plastid glycolysis proteins and greater mapped read depth at high latitudes in chlorophytes, although no such trend is observed in other algal groups. The latitudinal associations observed for diatom MAG abundances support expression trends shown in Fig. 3, although the chlorophyte MAG abundances point to the presence of novel plastid glycolytic pathways absent from the cultured species shown in Fig. 1.

**Fig. S13. Genotypes of *P. tricornutum* glycolysis knockout lines. A:** alignments of the two CRISPR regions targeted for mutagenesis of cpEnolase (Phatr3_J41515) and cpPGAM1A (Phatr3_J17086), and the genotypes obtained from Sanger sequences of homozygous CRISPR knockouts obtained for each gene. **B:** average relative expression level of each mutated gene, assessed by quantitative RT-PCR with two primer combinations and normalised against two housekeeping genes (RNA polymerase II and TATA binding protein), expressed as a % of the relative expression levels calculated in two empty vector expression controls. One-way *t*-test significance levels of the knockdown of gene expression in each knockout line compared to the empty vector controls are provided. Knockout lines shown in this figure were used for growth and integrative ‘omic analyses as per **Figs. 3-6**.

**Fig. S14. Absolute and individual growth phenotypes of cpEnolase and cpPGAM1A CRISPR-Cas9 knockout mutant lines. A:** growth curves of knockout lines, shown as per Fig. 4, but with absolute as opposed to logarithmic cell concentrations B: scatterplot showing the average and standard deviation relative growth rates for each cell line studied under 19C CL (vertical) and 19C LD (horizontal axis). Each point corresponds to an individual line, with genotype indicated by point colours, and standard deviations of growth rates by error bars. Despite individual variances in growth rate between lines, knockout lines show consistently slower growth than empty vector controls under both conditions, particularly 19C CL.

**Fig. S15. Measured photo-physiology of glycolysis knockout lines A:** Curves for **(i-ii)** relative electron transport (rETR) of photosystem II fitted as a function of light intensity) and **(iii-iv)** photoprotective non-photochemical quenching (NPQ) fitted as a function of E. Separate values are shown for cultures in CL **(i, iii)** and LD **(ii, iv)** growth conditions. Data points are the mean between the average values (*n*=2-4) measured in each strain within each genotype (Control = 2, cpEnolase complemented = 2, cpPGAM1A complemented = 3, cpEnolase knockout = 6, cpPGAM1A knockout = 3). **B:** Violin plots of PSII functional absorption cross-section (σPSII), measured with a MINIFIRe spectrometer for glycolysis mutant versus control lines under each growth conditions. Significantly different values observed for knockout and complementation mutants relative to control lines (one-way ANOVA, P < 0.05) are asterisked, with asterisk colour corresponding to the line considered. Each boxplot includes all measured/ fitted values for each strain within a mutant line. The absence of clear photosynthetic defects contrasts with the diminished growth of knockout lines, as per Fig. 4.

**Fig. S16: Bar plots of the mean and standard deviation of the ratios of 39 metabolites assessed by GC-MS in plastid glycolysis mutant lines under the three tested experimental conditions.** Data support the Volcano Plots shown in Fig. 6. Metabolites are sorted in ranked decreasing accumulation in mutant lines over all three conditions. Metabolites inferred to be differentially accumulated (one-way ANOVA) in each mutant line and condition are asterisked.

**Fig. S17. Lipid accumulation profiles under 19C LD conditions**. **A:** Volcano Plots showing (horizontal axis) log_2_ accumulation ratios and (vertical axis) –log_10_ one-way ANOVA, two-tailed Pvalues for separation of mean proportions of specific fatty acids, across all fatty acids observed in a specific lipid class in glycolysis mutants versus control lines, supporting the global scatter- and violin plots shown in Fig. 7. Specific lipids that show extreme (P < 0.01) differences in accumulation between both mutant genotypes and control lines are labelled, and coloured by lipid class. **B:** Bar plots showing total DGTA lipid class distributions in all three lines under these conditions. These data suggest limited changes in glycolysis mutant lipid architecture, barring a probable over-accumulation of *sn-*1 C16 in gycolysis mutant lipid pools, and corresponding under-accumulation of *sn-*1 C20 in mutant DGTA pools.

**Fig. S18. Lipid accumulation profiles under 19C CL conditions. A:** Volcano plots showing (horizontal axis) log_2_ accumulation ratios and (vertical axis) –log_10_ one-way ANOVA, two-tailed Pvalues for separation of mean proportions of specific fatty acids, across all fatty acids observed in a specific lipid class in glycolysis mutants versus control lines, and **B: bar plots** of SQDG and DGTA accumulation in lines harvested under **19C CL.** Data are shown as per **Fig. S17** and support global scatter- and violin plots shown in Fig. 7. These data suggest similar changes in glycolysis mutant lipid architecture to **19C LD**, including probable over-accumulations of *sn-*1 C16 in lieu of C20 and C14 in cpEnolase and cpPGAM1A mutant SQDG and MGDG pools.

**Fig. S19. Lipid accumulation profiles under 8C CL conditions.** Volcano plots showing (horizontal axis) log_2_ accumulation ratios and (vertical axis) –log_10_ ANOVA Pvalues for separation of mean proportions of specific fatty acids, across all fatty acids observed in a specific lipid class in cpEnolase mutants versus control lines, and cpEnolase mutants versus cpPGAM1A mutants harvested under 8C CL conditions. Data are shown as per **Fig. S17** and support global scatter- and violin plots shown in Fig. 7. No sigificantly differentially accumulated (P < 10^-05^) lipids were observed in corresponding comparisons of cpPGAM1A mutants and control lines. These data suggest specific overaccumulations in short-chain *sn-*1 MGDG, and *sn*-2 SQDG, and C20 *sn-1* DGTA in cpEnolase mutants compared to other lines.

**Fig. S20. Schematic diagram of the reaction kinetics measured for *P. tricornutum* cpEnolase and cPGAM1A enzymes**. The measured activities of this assay are shown in Fig. 8. Common enzymes are shown in green, enzymes unique to the glycolytic assay in blue, and enzymes unique to the gluconeogenic assay in red. Reaction intermediates and reversible substrates are shown in yellow and gray respectively.

**Supplemental Dataset S2. Transcriptional and localisation patterns of ptPGAM and ptEnolase genes in *Phaeodactylum tricornutum*.**

**Supplemental Dataset S3. *Tara* Oceans analysis of diatom plastidial glycolysis.**

**Supplemental Dataset S4. Genotyping, growth dynamics and photophysiology in *P. tricornutum* plastidial glycolysis mutant lines.**

**Supplemental Dataset S5. Differentially and conditionally expressed genes in *P. tricornutum* plastidial glycolysis mutants.**

**Supplemental Dataset S6. Lipid and metabolite profiles of *P. tricornutum* plastidial glycolysis mutant lines, and measured reaction kinetics of expressed enzymes.**

