## Supplemental Figures for "Complementary environmental analysis and functional characterization of a plastid diatom lower glycolytic-gluconeogenesis pathway"

**Short Title: Probing Diatom Chloroplast Lower-Half Glycolysis**

Richard G. Dorrell[1,2,3], Youjun Zhang[4,5], Yue Liang[6,7], Nolwenn Gueguen[8], Tomomi Nonoyama[1,9], Dany Croteau[10], Mathias Penot[1,2,3], Sandrine Adiba[1], Benjamin Bailleul[10], Valérie Gros[8], Juan José Pierella Karlusich[1,2,11], Nathanaël Zweig[1,2], Alisdair R. Fernie[5,6], Juliette Jouhet[8], Eric Maréchal[8] and Chris Bowler[1,2]

[1]Institut de Biologie de l'ENS (IBENS), Département de Biologie, École Normale Supérieure, CNRS, INSERM, Université PSL, 75005 Paris, France
[2]CNRS Research Federation for the study of Global Ocean Systems Ecology and Evolution, FR2022/Tara Oceans GOSEE, 3 rue Michel-Ange, 75016 Paris, France
[3]Laboratory of Computational and Quantitative Biology (LCQB), Institut de Biologie Paris-Seine (IBPS), CNRS, INSERM, Sorbonne Université, 75005 Paris, France
[4]Center of Plant Systems Biology and Biotechnology, 4000 Plovdiv, Bulgaria
[5]Max-Planck-Institute of Molecular Plant Physiology, Am Mühlenberg 1, 14476, Potsdam-Golm, Germany
[6]Key Laboratory of Seed Innovation, Institute of Genetics and Developmental Biology, Chinese Academy of Sciences, Beijing, China
[7]Center of Deep Sea Research, Institute of Oceanology, Center for Ocean Mega-Science, Chinese Academy of Sciences, Qingdao 266071, China
[8]Laboratory for Marine Mineral Resources, Pilot National Laboratory for Marine Science and Technology, Qingdao 266237, China
[9]Laboratoire de Physiologie Cellulaire et Végétale, CNRS, Univ. Grenoble Alpes, CEA, INRAE, IRIG, 17 rue des Martyrs, 38000 Grenoble, France
[10]Division of Biotechnology and Life Science, Institute of Engineering, Tokyo University of Agriculture and Technology, 2-24-16, Naka-cho, Koganei, Tokyo 184-8588, Japan
[11]Institut de Biologie Physico-Chimique (IBPC), Université PSL, 75005 Paris, France
[12]Present address: FAS Division of Science, Harvard University, Cambridge, MA, USA

### Supplemental Figures and Datasets

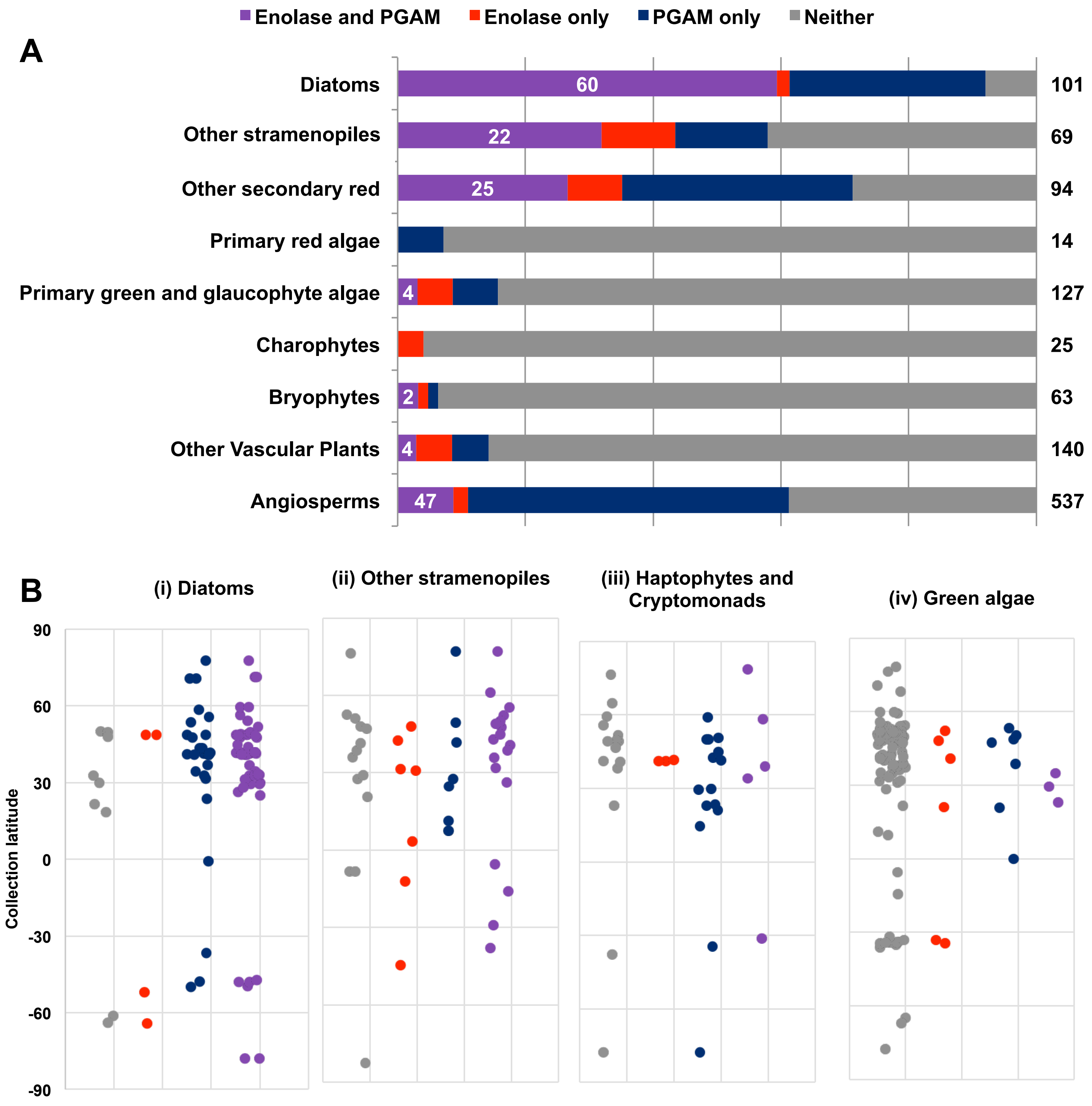


**Fig. S1. Distribution of lower plastid glycolysis-glucneogenesis across photosynthetic eukaryotes. A:** Occurrence of plastid-targeted enolase and PGAM enzymes across 1,673 plant and algal genomes and transcriptomes, inferred using reciprocal BLAST best hit of *P. tricornutum* query enzymes as per **Fig. 1B** and **1C,** PFAM domain annotations, and in silico targeting predictions with TargetP and PredAlgo (primary chloroplast bearing lineages) and HECTAR and ASAFind (secondary lineages). **B:** scatterplots of collection site latitude for (**i)** diatoms, (ii) other stramenopiles, **(iii)** cryptomonads and haptophytes and **(iv)** green algae with detectable enolase and PGAM enzymes, divided by presence of inferred plastid-targeted isoforms. Notably, diatoms lacking both plastid-targeted glycolysis enzymes do not occur outside of low and intermediate latitudes (50°N in the northern and 60°S in the southern hemisphere) compared to other groups, which show no significant association between plastid glycolysis and latitude. The data in this figure were subselected for the phylogenies shown in **Fig. 1** and support the latitudinal correlations revealed by Tara analysis of **Fig. 3.**

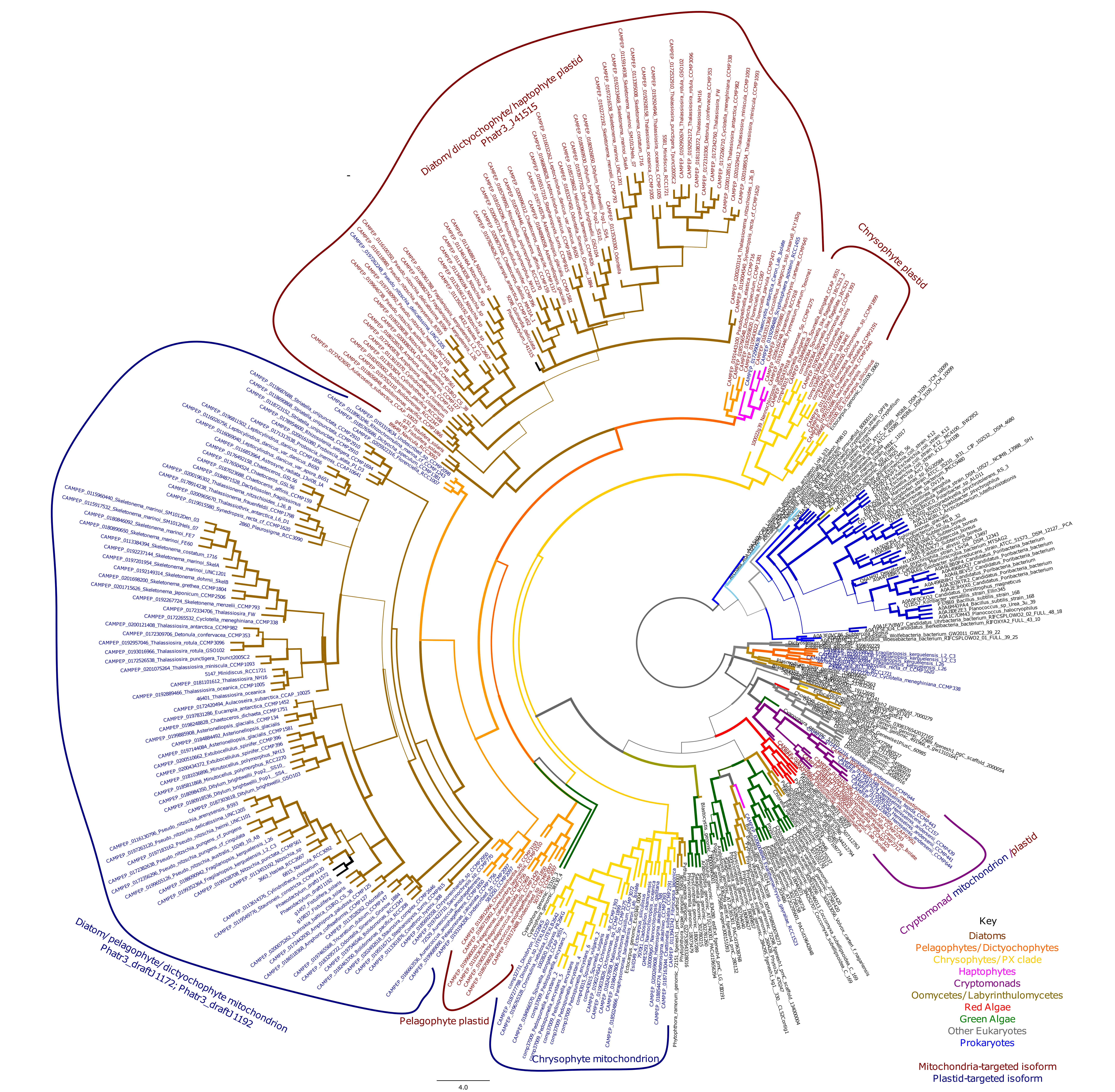


**Fig.S2**. **Consensus topology of a 380 taxa x 413 aa alignment of Enolase sequences.** Sequences represent a sample all organelle-targeted isoforms from cryptomonads, haptophytes and stramenopiles and representatives from a densely-sampled dataset of 151 taxonomic groups across the tree of life (Dorrell et al., 2021). The tree topology shown is the consensus of the best-scoring rAxML trees identified using three substitution matrices: GTR, JTT, and WAG. Branch thickness corresponds to frequency of consensus tree topology recovery in individual trees; branches are coloured by taxonomic affiliation; and tips (cryptomonads, haptophytes and stramenopiles only) by predicted *in silico* localisation. Individual clades (considering both taxonomic origins and inferred localisation) of organelle-targeted enolase isoform are labelled with coloured brackets.

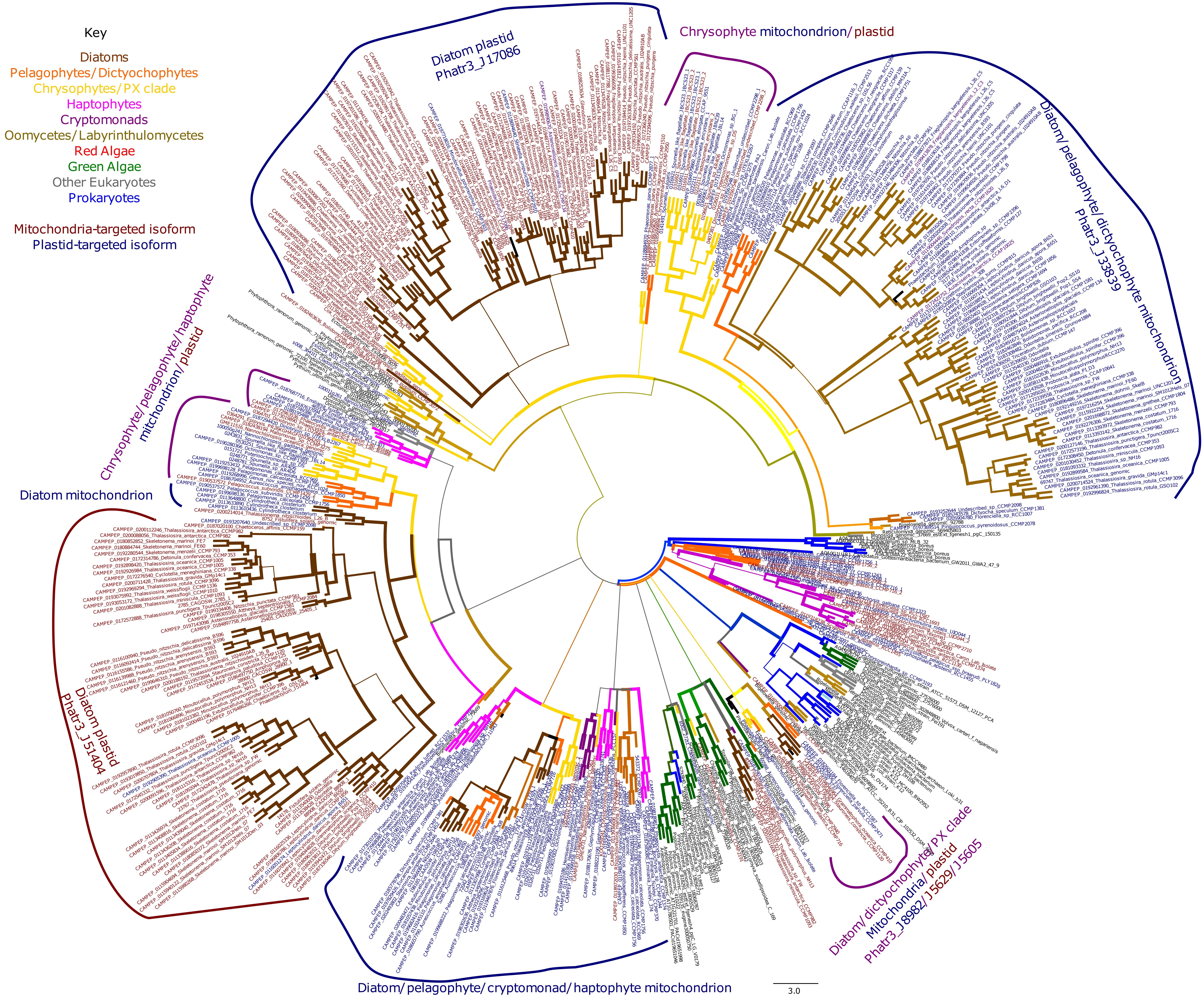


**Fig. S3. Consensus topology of a 220 aa x 560 taxa alignment of PGAM isoform 1 sequences.** Data are shown as per **Fig. S1**, extending on the topology shown in **Fig. 1C**.

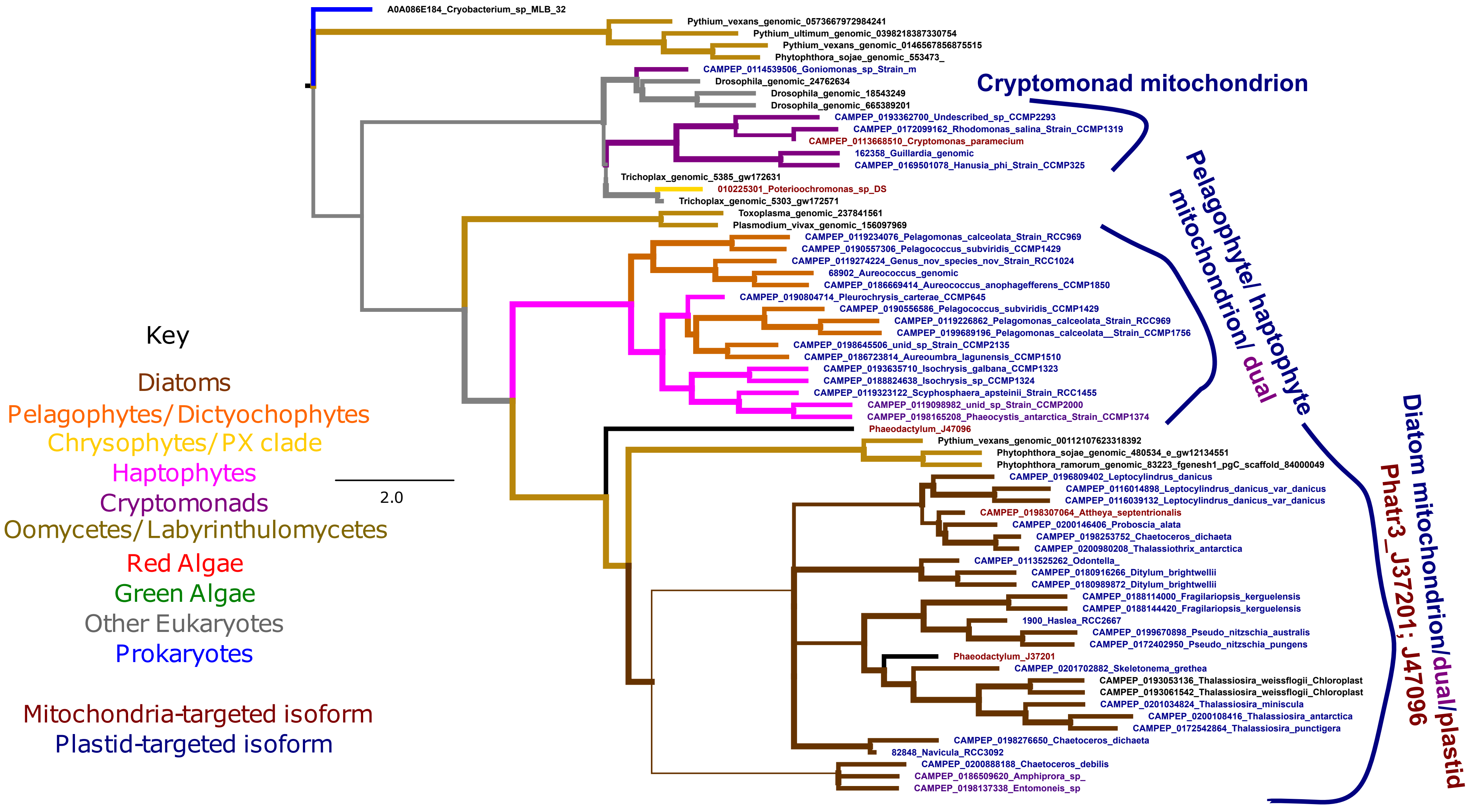


**Fig. S4. Consensus phylogeny of a 235 aa x 66 taxa alignment of PGAM isoform 2 sequences.** Data are shown as per **Fig. S1**, forming a complement to the PGAM1 topologies shown in **Figs. 1C** and **S3**.

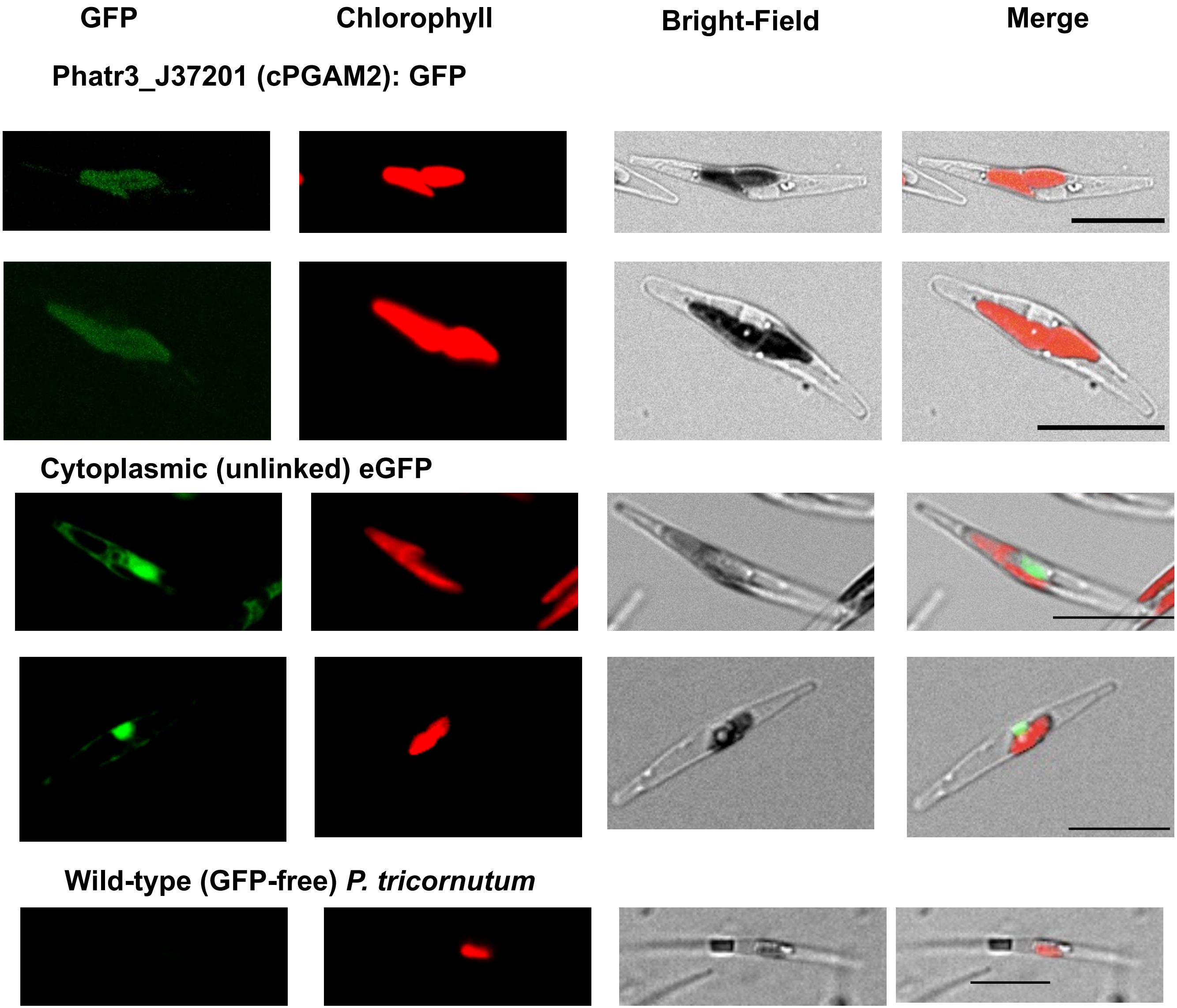


**Fig. S5. PGAM2 and control confocal microscopy images for *P. tricornutum* plastid glycolysis proteins.** Images complement those shown in **Fig. 2A.**

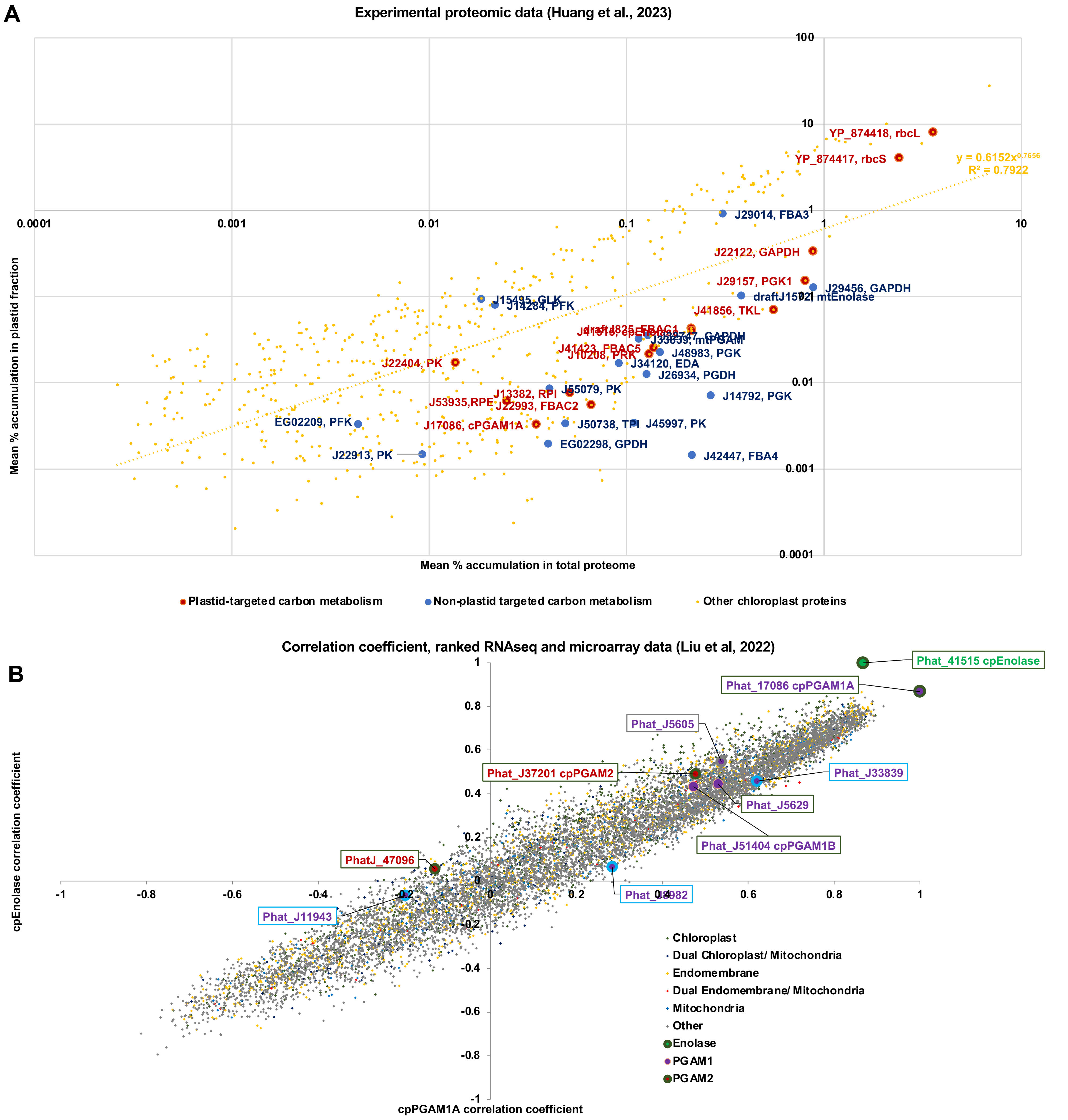


**Fig. S6**. **Relative abundances and transcriptional coordination of plastidial enolase, PGAM1 and PGAM2. A:** relative % accumulation of different plastid- and mitochondrial-associated carbon metabolism proteins (large points) and all known plastid-encoded or inferred plastid-targeted proteins (small points, with power-law trendline) for experimental proteomics of *Phaeodactylum tricornutum* total cell proteomes (horizontal axis) or plastid-enriched protein fractions (vertical axis), following Huang et al, 2023. Only proteins detected in at least two replicates of plastid-enriched and total cell fractions are included, and abundances are renormalized based on the total abundances of these proteins. Both cpEnolase (Phatr3_J41515) and cpPGAM1A (Phatr3_J17086) are detected in the plastid-associated proteome fraction with similar frequencies to other plastid carbon metabolism proteins, whereas other predicted plastid-targeted PGAMs (e.g. cpPGAM1B, Phatr3_J51404; cPGAM2, Phatr3_J37201) are not. **B:** Spearman correlation coefficients of all genes across the version 3 annotation of the *Phaeodactylum tricornutum* genome against cpEnolase and cpPGAM1A; calculated from ranked transcript data (Liu et al, 2022) from two merged datasets, PhaeoNet (Ait-Mohamed et al., 2020) and DiatomPortal (Ashworth et al., 2016). Genes are coloured by inferred in silico localisation; and other annotated enolase and PGAM enzymes in the *Phaeodactylum* genome are shown as large, labelled points. Notably, while cpEnolase and cpPGAM1A show a strong positive (r > 0.8) coregulation to one another, other PGAM isoforms (e.g. cpPGAM1B, cpPGAM2) show less apparent coregulation to either of these two genes. These data support the experimental localisations shown in **Fig.2A**, and identify cPGAM1A as the PGAM most likely to work cooperatively with cpEnolase from the homologues shown **in Fig. 2B**.

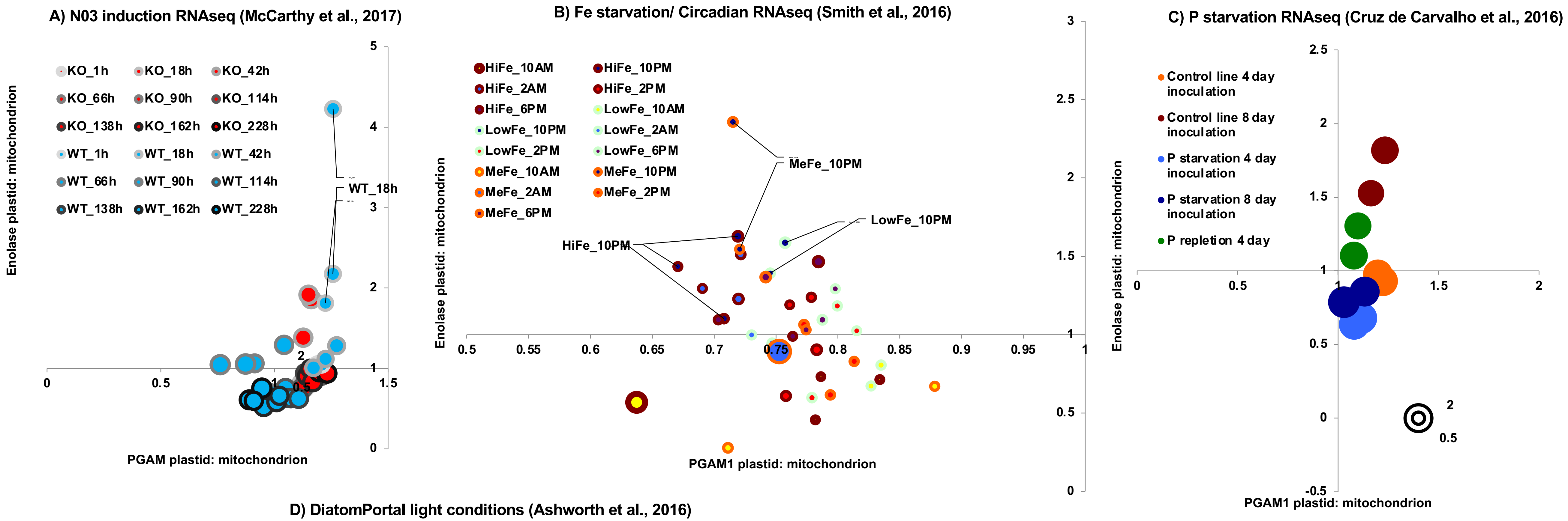


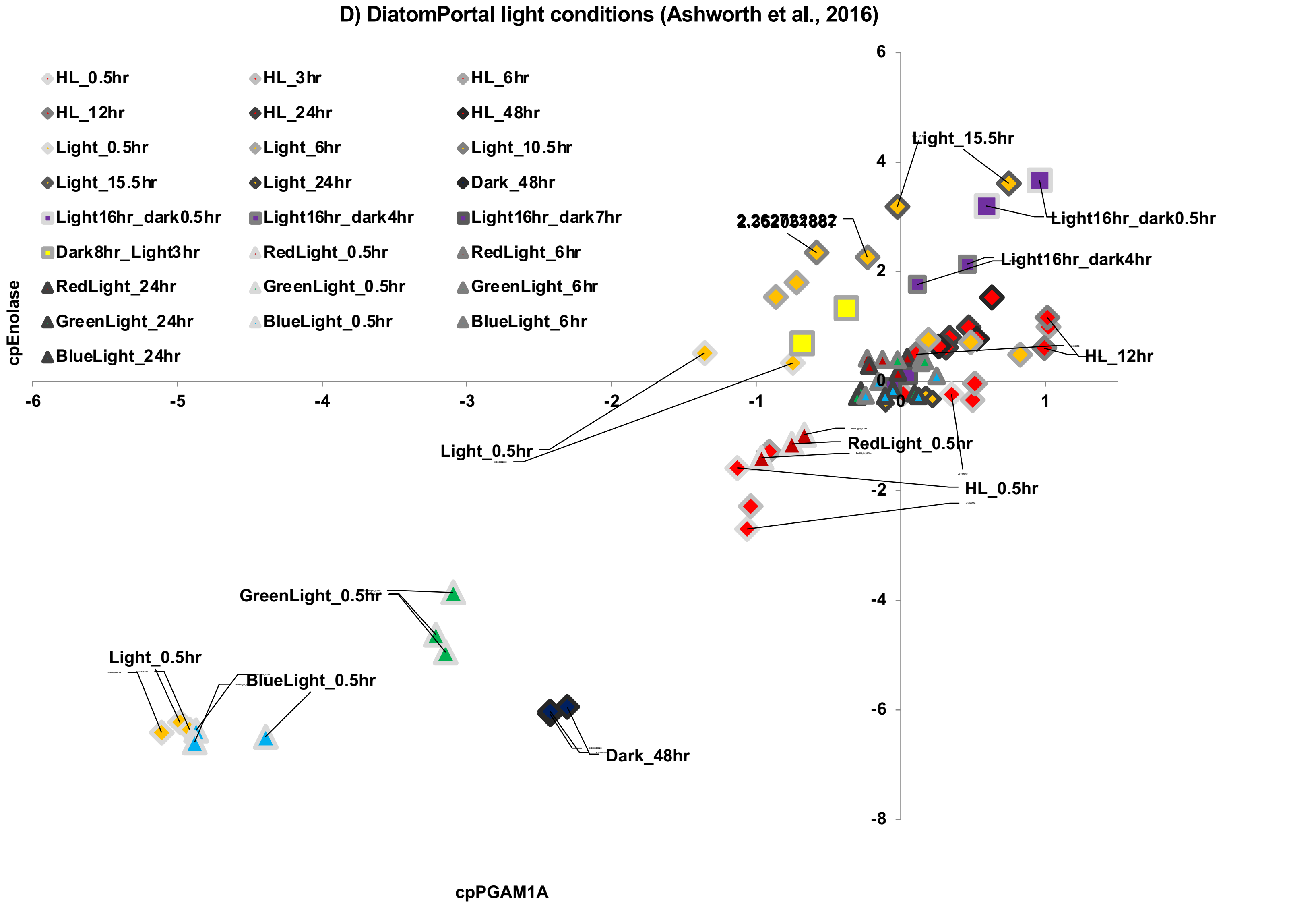


**Fig. S7. Relative transcriptional regulation of cpEnolase and cpPGAM under nutrient, light and temperature stress conditions. A-C:** the relative ratio of plastid to mitochondria-targeted gene isoform expression for enolase (vertical axis), PGAM1 (horizontal axis), and PGAM2 (bubble size) in published *Phaeodactylum* RNAseq data subject to nitrate limitation (McCarthy et al., 2017), phosphate starvation and addition (Cruz de Carvalho et al., 2016) and Circadian Fe enrichment and limitation (Smith et al., 2016). While limited transcriptional responses are identifiable in response to changing N, P or Fe availability, the relative transcription of plastidial to mitochondrial-targeted enolase is substantially greater in RNA sampled in long day (> 12h post-illumination) conditions in **C. D:** relative fold changes in plastidial enolase and PGAM1 expression in published microarray data assembled in Ashworth et al., 2016, under different illumination conditions. Both Enolase and PGAM1 show substantial downregulation in dark-incubated (48h) and short post-illumination-incubated (< 0.5h) cultures, and show the greatest positive fold expression changes respectively > 16h and > 12h post-illumination. Specific data points showing either very high or low cpEnolase and cpPGAM1A expression are labelled. These data support the qRT-PCR and *Tara* Oceans observations of correlation of diatom cpEnolase and cpPGAM expression in **Fig. 2** and **Fig. 3**.

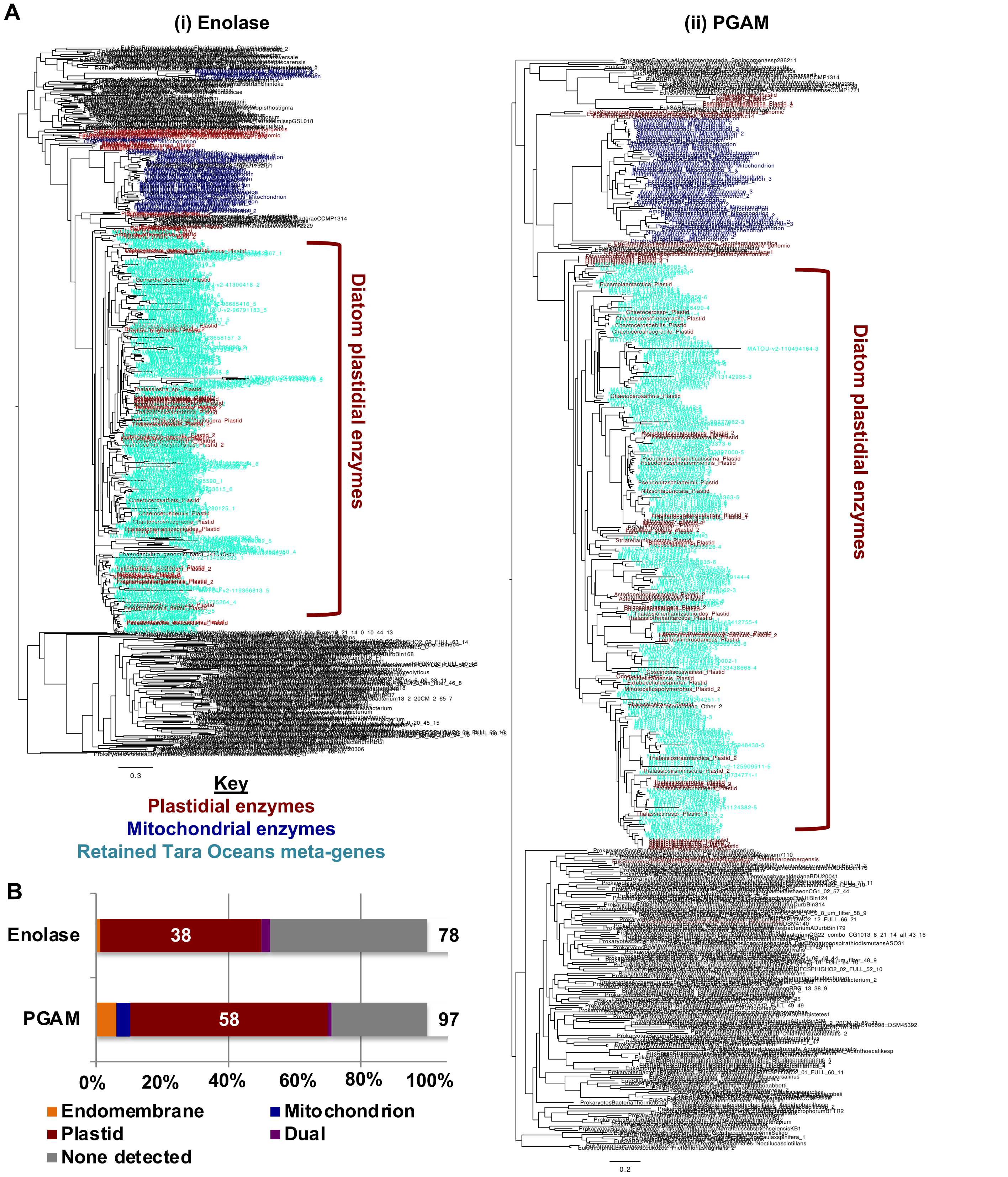


**Fig. S8. Identification of Tara Oceans homologues of diatom plastid-targeted enolase and PGAM enzymes**. **A:** Consensus rAxML JTT topologies of the phylogenetically verified *Tara* Oceans homologs of diatom plastidial enolase and PGAM enzymes and cultured species sequences, demonstrating reconciliation of retained homologs within monophyletic clades containing exclusively diatom plastidial isoforms amongst cultured species. **B:** *in silico* targeting predictions of all retrieved homologs inferred by BLAST alignment to be probably N-terminally complete, showing a strong enrichment in homologs with predicted plastid-targeting sequences. Sequences shown in this figure are analysed globally in **Fig. 3**.

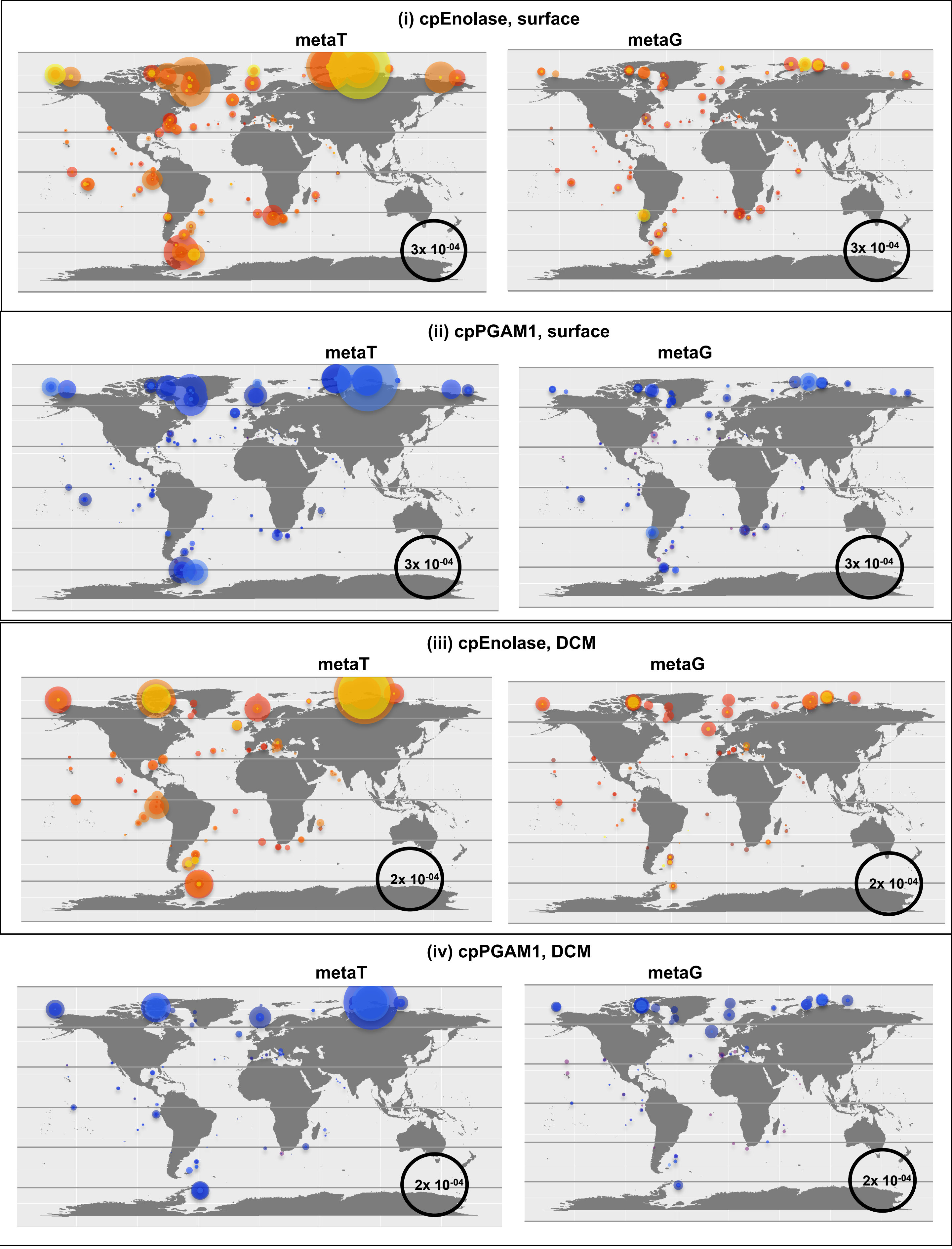


**Enolase:** 0.8-3/5 • 5-20 • 20-180 • 180-2000 µm

**PGAM:** 0.8-3/5 • 5-20 • 20-180 • 180-2000 µm

**Fig. S9. Relative abundances of *Tara* Oceans diatom plastid glycolysis meta-genes.** Plots show relative abundances of meta-genes that group with (**i, iii**) cpEnolase and (**ii, iv**) cpPGAM1 sequences over individual size fractions of (**i, ii**) surface and (**iii, iv**) DCM meta-transcriptome (left) and -genome (right) data. These data, which allows us to identify whether specific trends are observed in different kinds of diatom cells, ranging from the nano to micro-metre scales, support global trends observed across from all (unfiltered) size fractions and surface layers shown in **Fig. 3**.

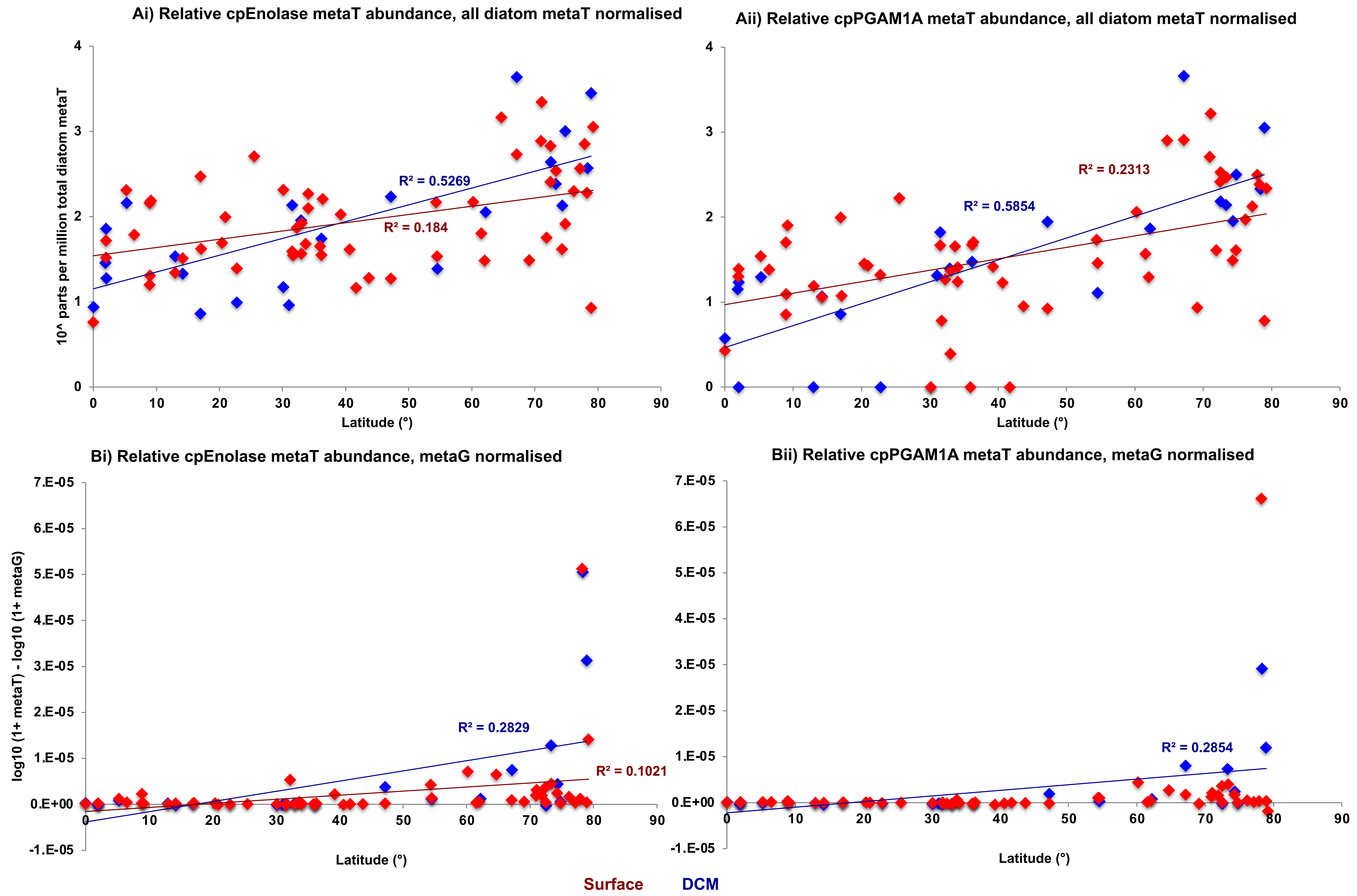


**Fig. S10. Normalised atitudinal regressions of *Tara* cpEnolase and cpPGAM1 sequences.** Scatterplots of *Tara* Ocean expression patterns of sequences assigned phylogenetically to diatom cpEnolase and cpPGAM1A against station latitude. Abundances are shown for 0.8-2000 μm surface and DCM sample meta-transcriptome data, and are normalised relative to **(A)** total diatom metaT abundances at each station and **(B)** the corresponding metaG abundances for diatom cpEnolase and cpPGAM1A. Best-fit (linear) regression lines are provided for each depth. In each case, a significant positive correlation between latitude and relative expression is observed, consistent with global distributions observed in **Fig. 3**.

(i) Relative PGAM2 metaT abundances

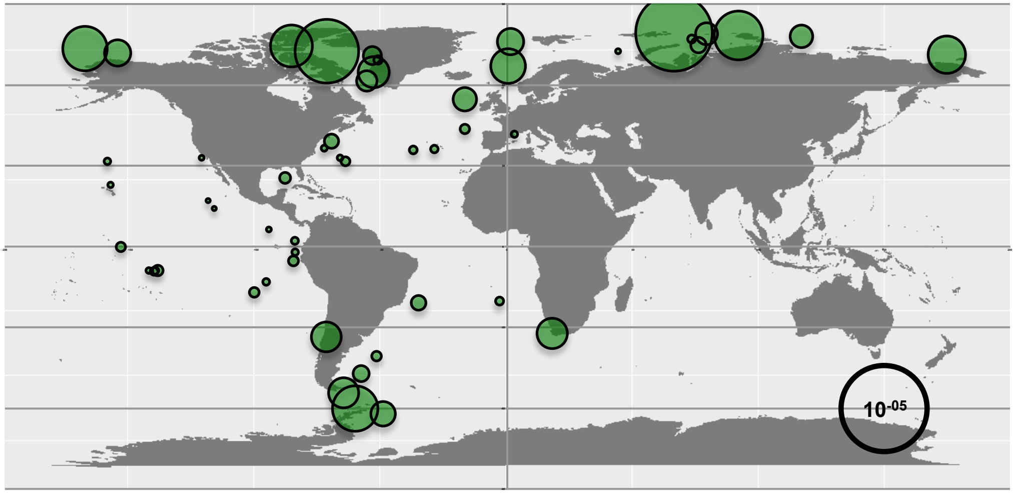


(ii) Relative PGAM2 metaG abundances

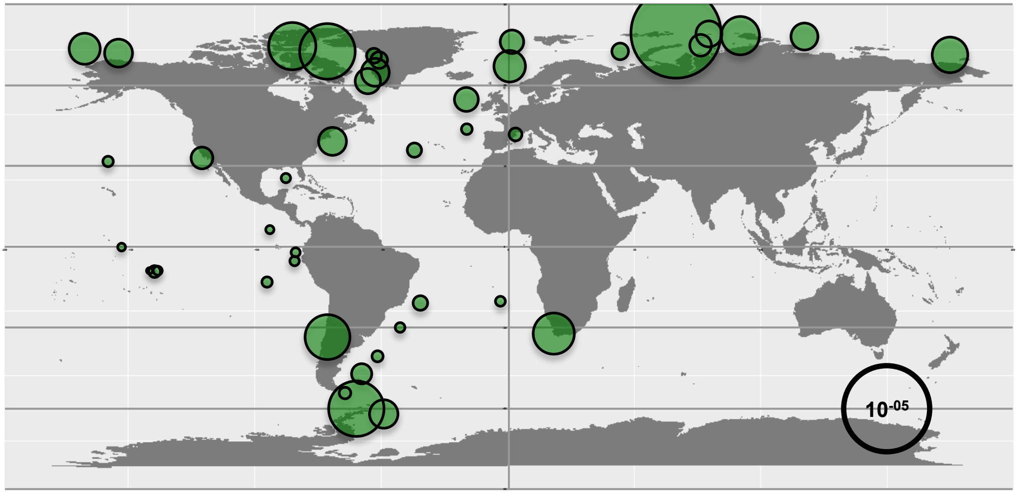


**Fig. S11. Total relative abundances of meta-genes phylogenetically reconciled to diatom PGAM2 in 0.8- 2000 µm filtered surface samples.** Plots showing **(i)** meta-transcriptome and **(ii)** meta-genome data, showing effective congruence between both, in contrast to the high latitudinal abundance specific to meta-transcriptome data for diatom cpEnolase and cpPGAM1 as per **Fig. 3.**

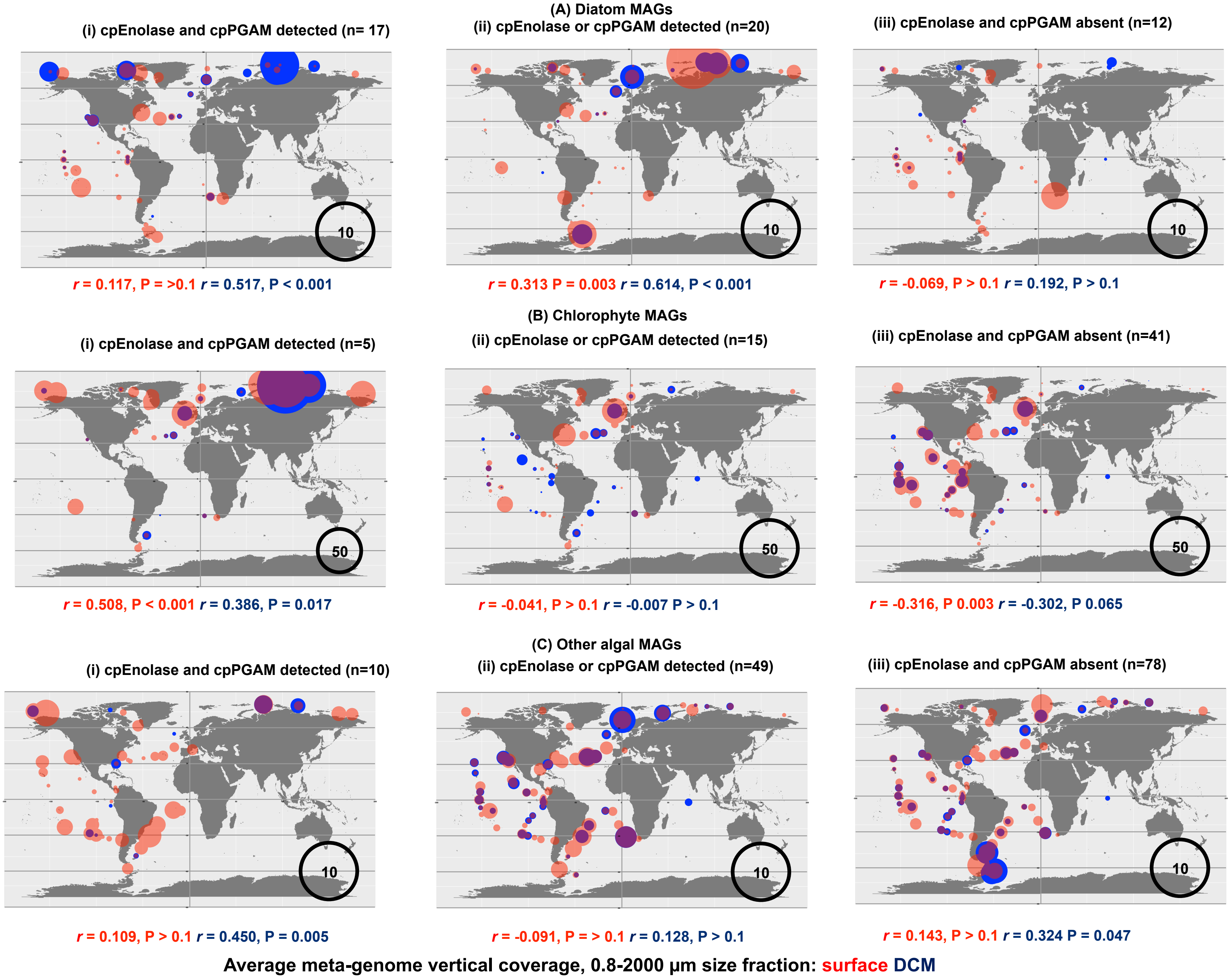


**Fig. S12. Occurrence and mean coverage depth of *Tara* Oceans MAGs divided by chloroplast glycolysis state.** This figure shows mapped distributions for *Tara* meta-genome assembled genomes (MAGs, Delmont et al., 2022) pertaining to (**A**) diatoms, (**B**) chlorophytes, and (**C**) other algal groups (haptophytes, pelagophytes, dictyochophytes, chrysophytes, bolidophytes, cryptomonads) containing members with inferred lower half plastidial glycolysis. MAGs are divided by the inferred occurrence of which chloroplast-targeted enolase and PGAM enzymes could be inferred using combined RbH to *Phaeodactylum* queries, PFAM annotation, and in silico targeting prediction (TargetP, WolfPSort, PredAlgo, HECTAR, ASAFind): (i) detection of plastid-targeted homologues of both enzymes; (**ii**) detection of cpEnolase or cpPGAM only; (**iii**) detection of neither. Bubble sizes correspond to the mean vertical coverage of meta-gene reads recruited to each MAG as a proxy of abundance. In each case the linear correlation and P-value (two-tailed *t*-test) of correlation between mean vertical mapped depth and absolute latitude is provided. Notably there is a positive correlation between the retrieval of either plastid glycolysis protein and greater mapped read depth at high latitudes in diatoms, and also the retrieval of both plastid glycolysis proteins and greater mapped read depth at high latitudes in chlorophytes, although no such trend is observed in other algal groups. The latitudinal associations observed for diatom MAG abundances support expression trends shown in **Fig. 3**, although the chlorophyte MAG abundances point to the presence of novel plastid glycolytic pathways absent from the cultured species shown in **Fig. 1**.

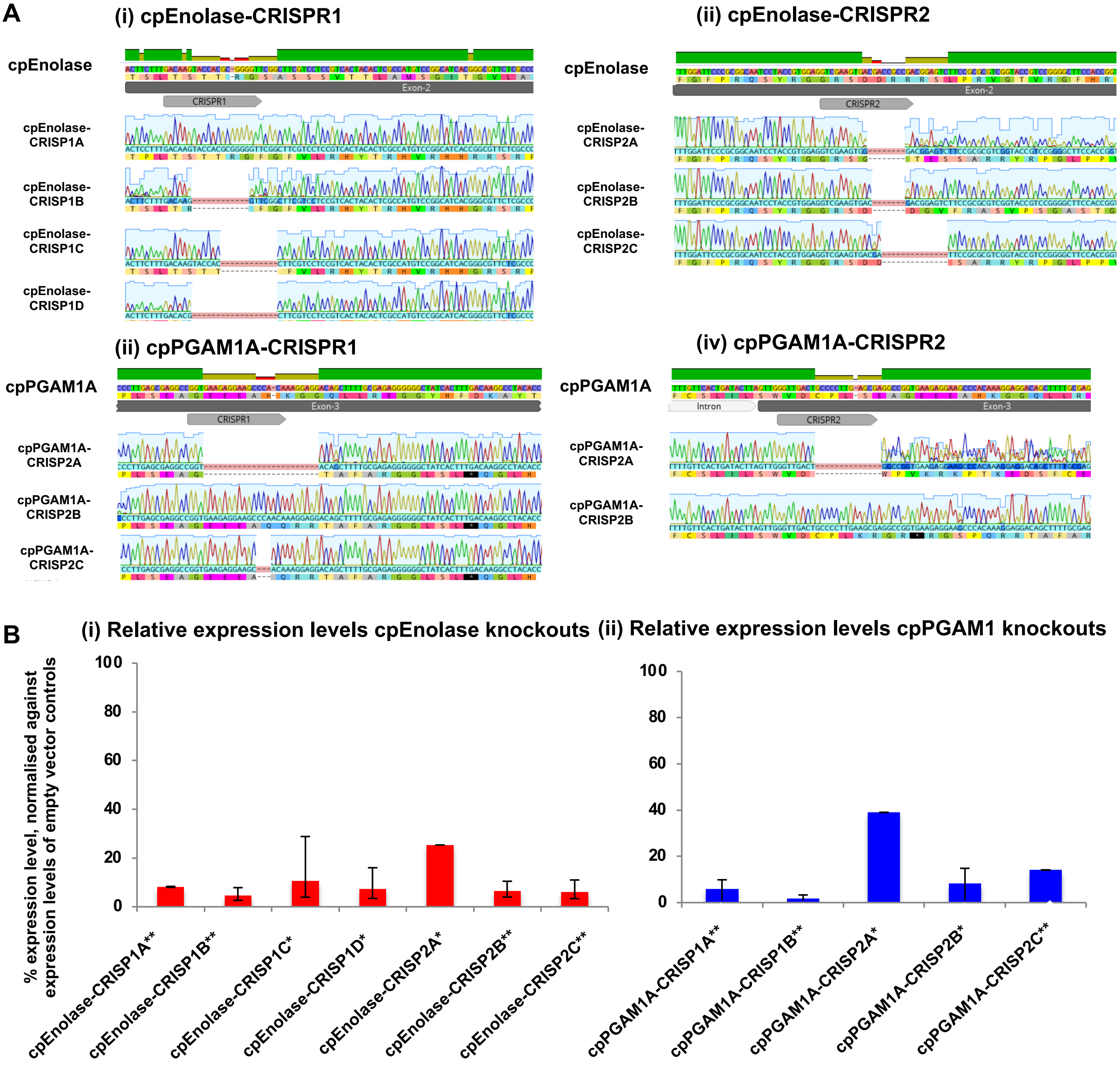


**Fig. S13. Genotypes of P. tricornutum glycolysis knockout lines. A:** alignments of the two CRISPR regions targeted for mutagenesis of cpEnolase (Phatr3_J41515) and cpPGAM1A (Phatr3_J17086), and the genotypes obtained from Sanger sequences of homozygous CRISPR knockouts obtained for each gene. **B:** average relative expression level of each mutated gene, assessed by quantitative RT-PCR with two primer combinations and normalised against two housekeeping genes (RNA polymerase II and TATA binding protein), expressed as a % of the relative expression levels calculated in two emcpy vector expression controls. One-way *t*-test significance levels of the knockdown of gene expression in each knockout line compared to the empty vector controls are provided. Knockout lines shown in this figure were used for growth and integrative ‘omic analyses as per **Figs. 4-7**.

*** Significant to P < 0.05**
**** Significant to P < 0.01**

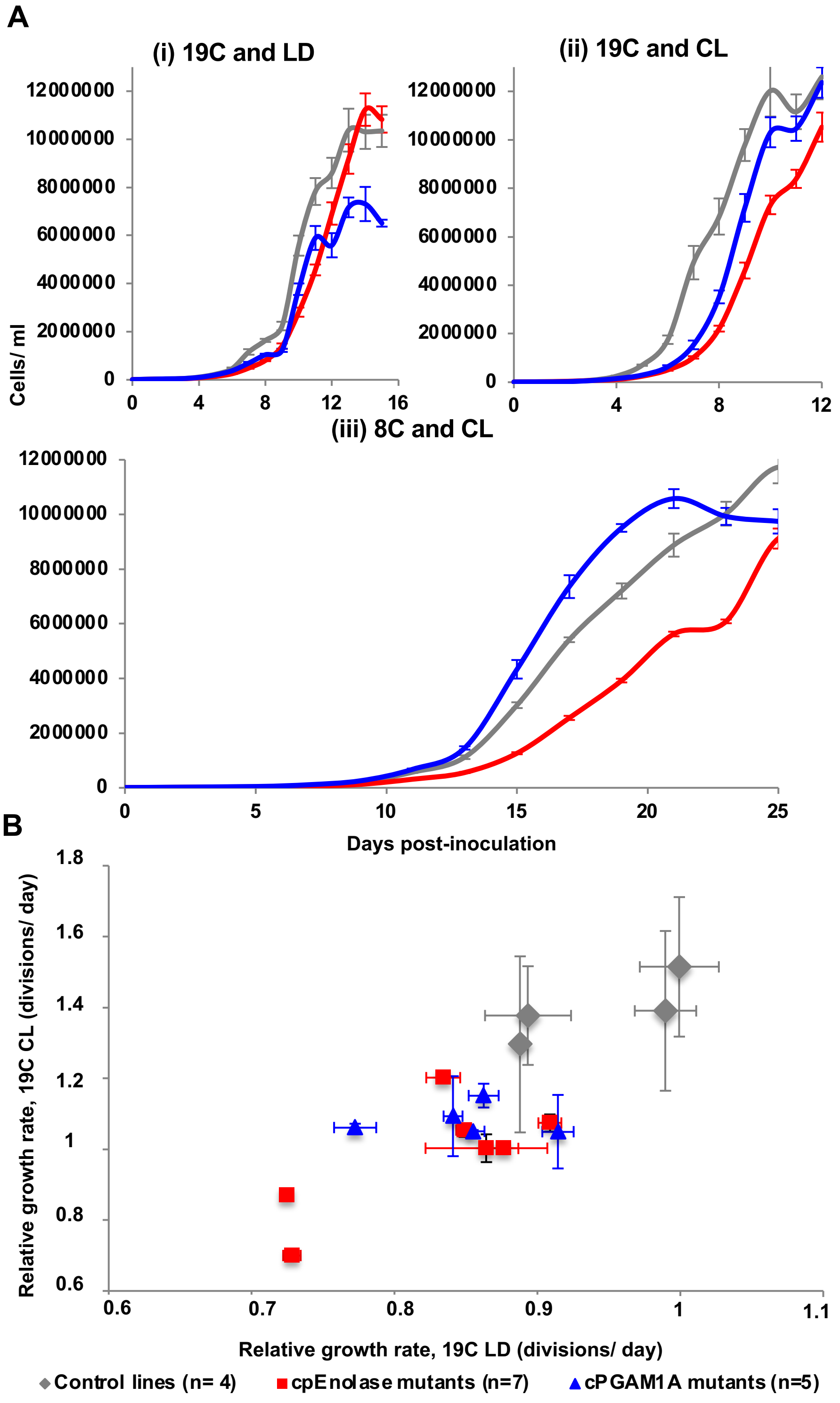


**Fig. S14. Absolute and individual growth phenotypes of cpEnolase and cpPGAM1A CRISPR-Cas9 knockout mutant lines. . A:** growth curves of knockout lines, shown as per **Fig. 4,** but with absolute as opposed to logarithmic cell concentrations **B:** scatterplot showing the average and standard deviation relative growth rates for each cell line studied under 19C CL (vertical) and 19C LD (horizontal axis). Each point corresponds to an individual line, with genotype indicated by point colours, and standard deviations of growth rates by error bars. Despite individual variances in growth rate between lines, knockout lines show consistently slower growth than empty vector controls under both conditions, particularly 19C CL.

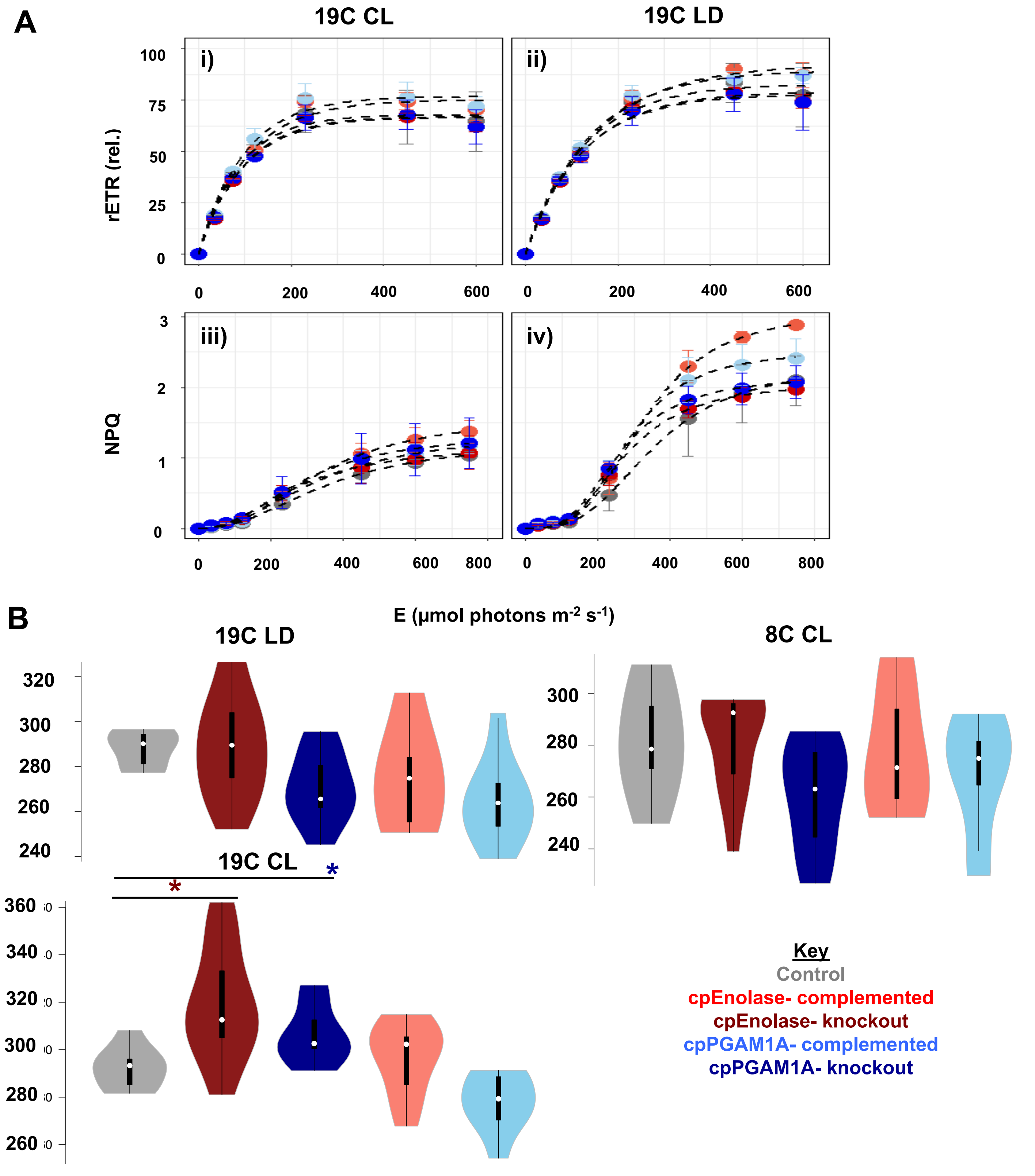


**Fig. S15. Measured photo-physiology of glycolysis knockout lines. A:** Curves for **(i-ii)** relative electron transport (rETR) of photosystem II fitted as a function of light intensity (E) and **(iii-iv)** photoprotective non-photochemical quenching (NPQ) fitted as a function of E. Separate values are shown for cultures in CL **(i, iii)** and LD **(ii, iv)** growth conditions. Data points are the mean between the average values (*n*=2-4) measured in each strain within each genotype (Control = 2, cpEnolase complemented = 2, cpPGAM1A complemented = 3, cpEnolase knockout = 6, cpPGAM1A knockout = 3). **B:** Violin plots of PSII functional absorbtion cross-section (σPSII), measured with a MINIFIRe spectrometer for glycolysis mutant versus control lines under each growth conditions. Significantly different values observed for knockout and complementation mutants relative to control lines (one-way ANOVA, $P < 0.05$) are asterisked, with asterisk colour corresponding to the line considered. Each boxplot includes all measured/ fitted values for each strain within a mutant line. The absence of clear photosynthetic defects contrast with the diminished growth of knockout lines, as per **Fig. 4**

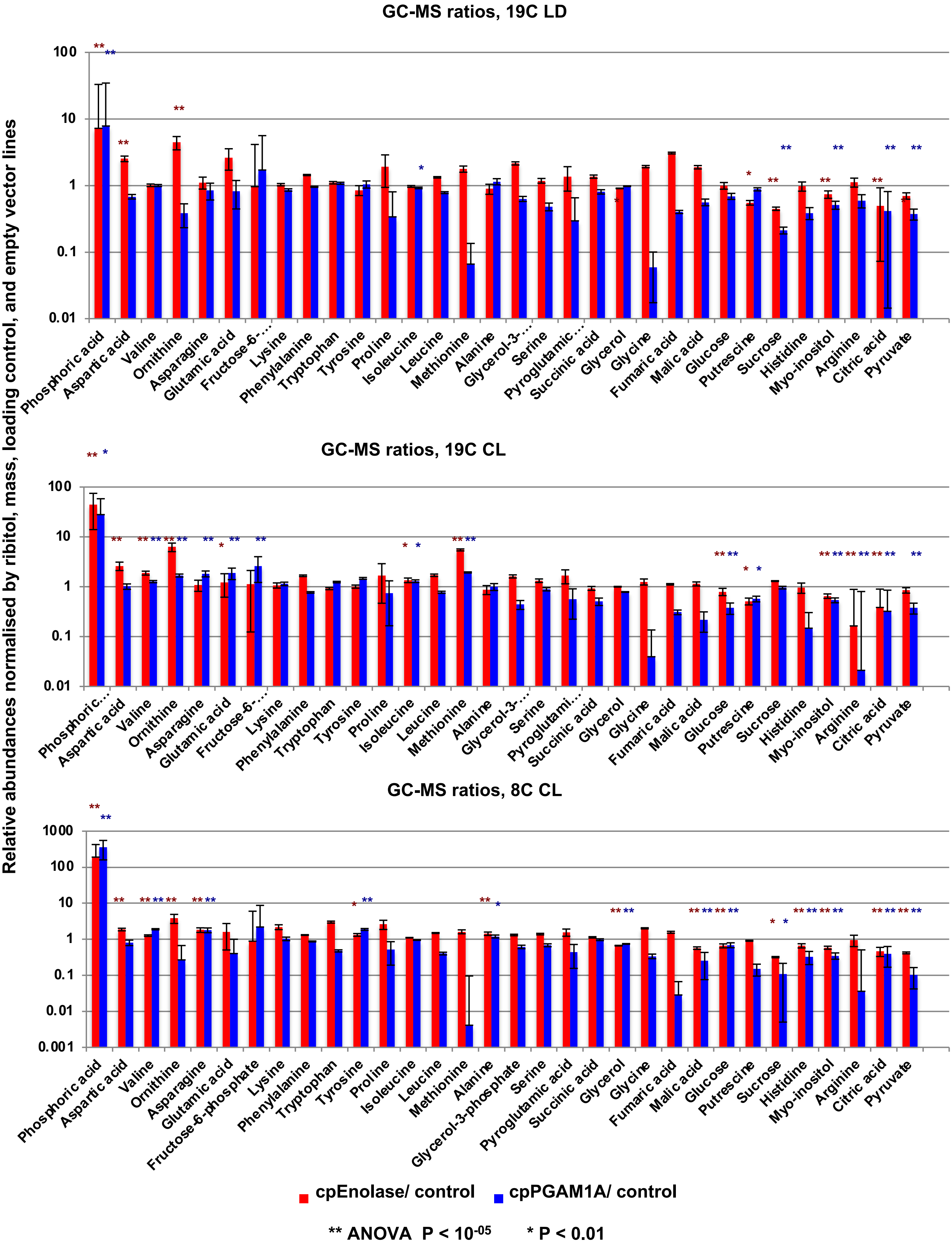


**Fig. S16. Bar plots of the mean and standard deviation of the ratios of 39 metabolites assessed by GC-MS in plastid glycolysis mutant lines under the three tested experimental conditions.** Data support the Volcano plots shown in **Fig. 6**. Metabolites are sorted in ranked decreasing accumulation in mutant lines over all three conditions. Metabolites inferred to be differentially accumulated (one-way ANOVA) in each mutant line and condition are asterisked.

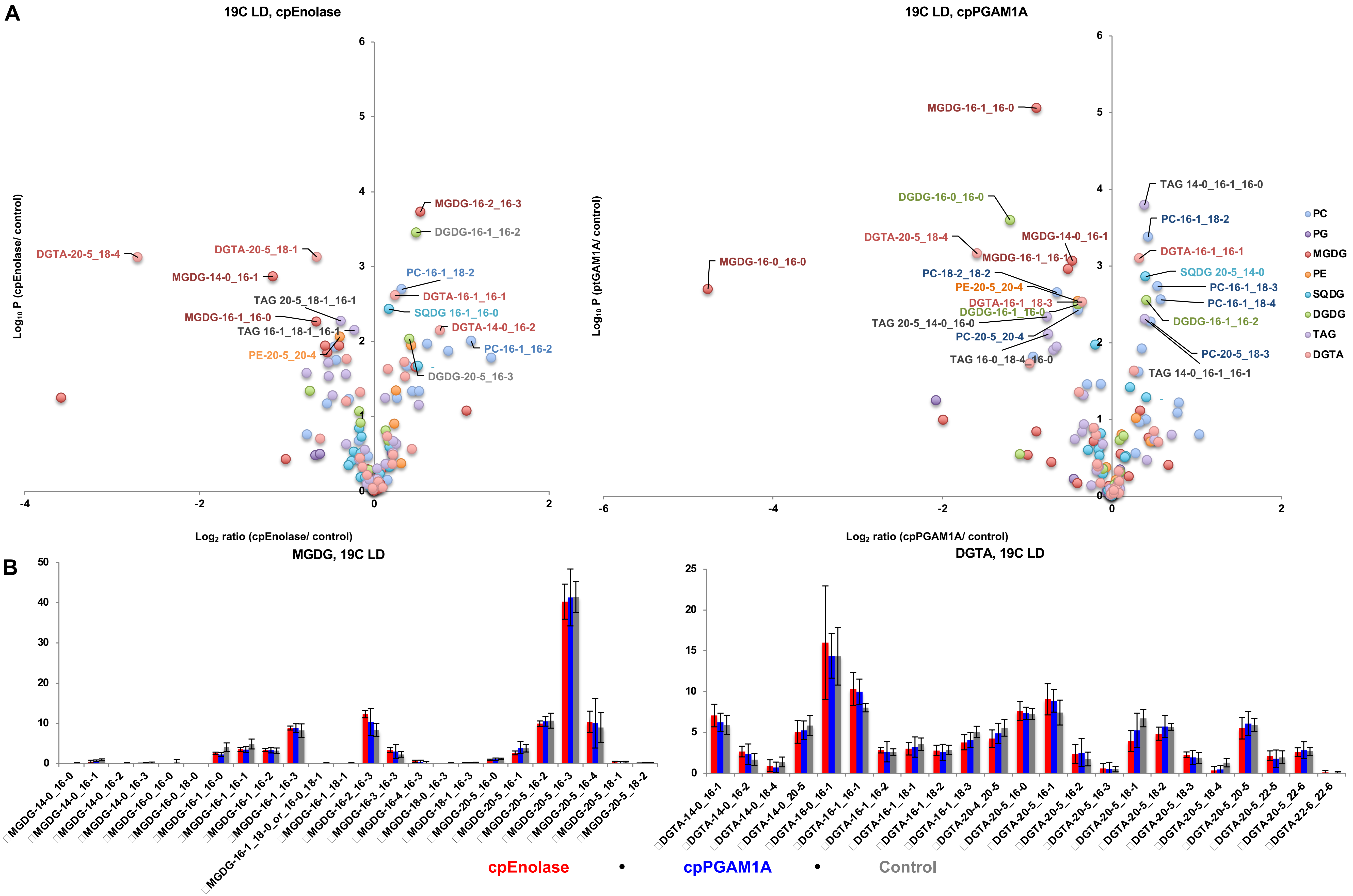


**Fig. S17. Lipid accumulation profiles under 19C LD conditions. A:** Volcano plots showing (horizontal axis) $\log_2$ accumulation ratios and (vertical axis) $-\log_{10}$ one-way ANOVA, two-tailed Pvalues for separation of mean proportions of specific fatty acids, across all fatty acids observed in a specific lipid class in glycolysis mutants versus control lines, supporting the global scatter- and violin plots shown in **Fig. 7.** Specific lipids that show extreme ($P < 0.01$) differences in accumulation between both mutant genotypes and control lines are labelled, and coloured by lipid class. **B:** Bar plots showing total DGTA lipid class distributions in all three lines under these conditions. These data suggest limited changes in glycolysis mutant lipid architecture, barring a probable over-accumulation of *sn*-1 C16 in gycolysis mutant lipid pools, and corresponding under-accumulation of *sn*-1 C20 in mutant DGTA pools.

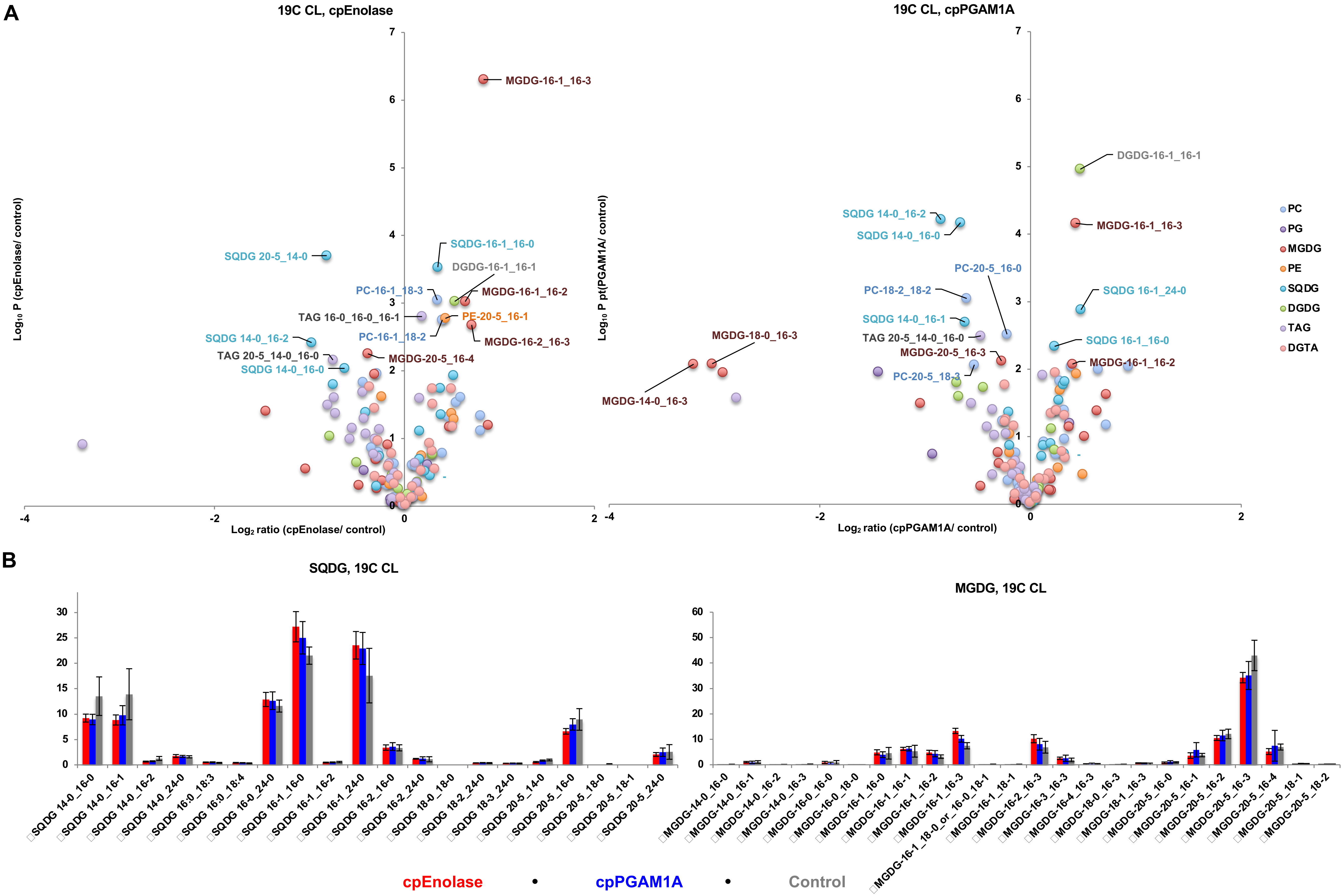


**Fig. S18. Lipid accumulation profiles under 19C CL conditions. A:** Volcano plots showing (horizontal axis) $\log_2$ accumulation ratios and (vertical axis) $-\log_{10}$ one-way ANOVA, two-tailed Pvalues for separation of mean proportions of specific fatty acids, across all fatty acids observed in a specific lipid class in glycolysis mutants versus control lines, and **B: bar plots** of SQDG and DGTA accumulation in lines harvested under **19C CL.** Data are shown as per **Fig. S17** and support global scatter- and violin plots shown in **Fig. 7.** These data suggest similar changes in glycolysis mutant lipid architecture to **19C LD**, including probable over-accumulations of *sn*-1 C16 in lieu of C20 and C14 in cpEnolase and cpPGAM1A mutant SQDG and MGDG pools.

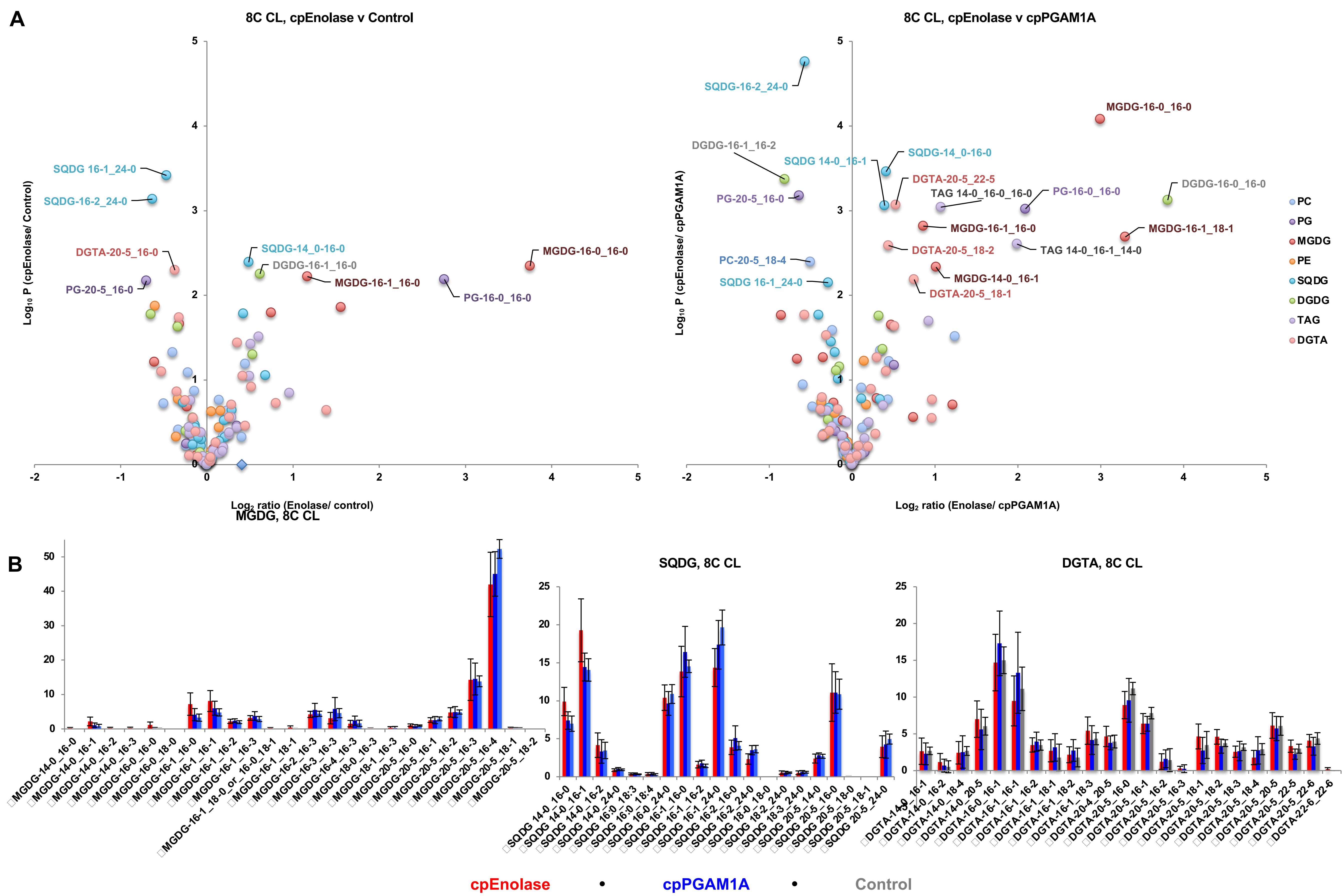


**Fig. S19. Lipid accumulaiton profiles under 8C CL conditions.** Volcano plots showing (horizontal axis) $\log_2$ accumulation ratios and (vertical axis) $-\log_{10}$ ANOVA Pvalues for separation of mean proportions of specific fatty acids, across all fatty acids observed in a specific lipid class in cpEnolase mutants versus control lines, and cpEnolase mutants versus cpPGAM1A mutants harvested under 8C CL conditions. Data are shown as per **Fig. S17** and support global scatter- and violin plots shown in **Fig. 7.** No sigificantly differentially accumulated ($P < 10^{-05}$) lipids were observed in corresponding comparisons of cpPGAM1A mutants and control lines. These data suggest specific overaccumulations in short-chain *sn*-1 MGDG, and *sn*-2 SQDG, and C20 *sn-1* DGTA in cpEnolase mutants compared to other lines.

A

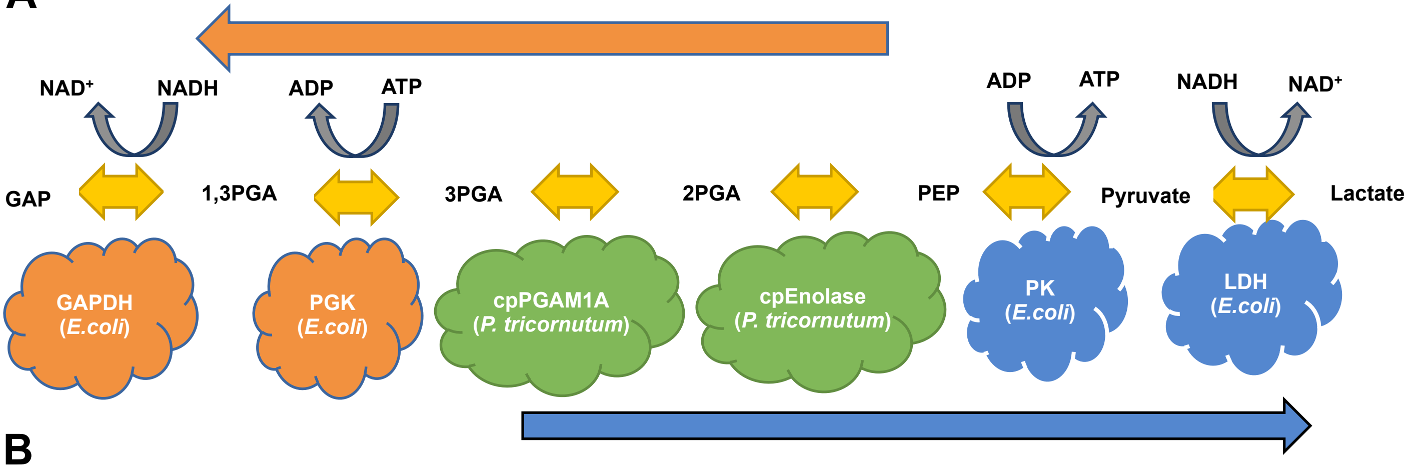


B

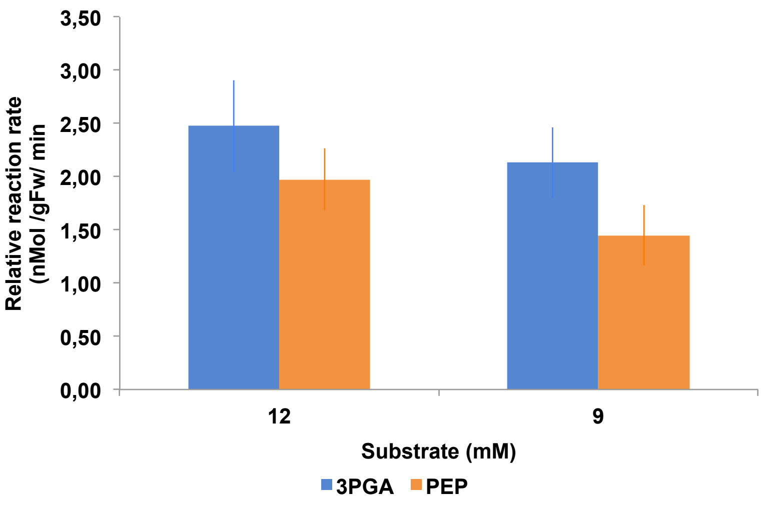


**Fig. S20. Reaction kinetics measured for *P. tricornutum* cpEnolase and cPGAM1A enzymes**. **A:** reaction pathway performed. The measured activities of this assay are shown in **Fig. 8.** Common enzymes are shown in green, enzymes unique to the glycolytic assay in blue, and enzymes unique to the gluconeogenic assay in red. Reaction intermediates and reversible substrates are shown in yellow and gray respectively. **B:** measured reaction rates (V) in each direction with 12mM and 9mM substrate. Error bars = one standard deviation

**Supplemental Dataset S1. Phylogenetic diversity of Enolase and PGAM sequences from across the tree of life.**

**Supplemental Dataset S2. Transcriptional and localisation patterns of ptPGAM and ptEnolase genes in *Phaeodactylum tricornutum*.**

**Supplemental Dataset S3. *Tara* Oceans analysis of diatom plastidial glycolysis.**

**Supplemental Dataset S4. Genotyping, growth dynamics and photophysiology in *P. tricornutum* plastidial glycolysis mutant lines.**

**Supplemental Dataset S5. Differentially and conditionally expressed genes in *P. tricornutum* plastidial glycolysis mutants.**

**Supplemental Dataset S6. Lipid and metabolite profiles of *P. tricornutum* plastidial glycolysis mutant lines, and measured reaction kinetics of expressed enzymes.**
